# Pro-apoptotic and anti-invasive properties underscore the tumor suppressing impact of myoglobin on subset of human breast cancer cells

**DOI:** 10.1101/2022.06.30.498102

**Authors:** Mostafa A. Aboouf, Julia Armbruster, Markus Thiersch, Franco Guscetti, Glen Kristiansen, Peter Schraml, Anne Bicker, Ruben Petry, Thomas Hankeln, Max Gassmann, Thomas A. Gorr

**Author notes:** Corresponding author: PD. Dr. Thomas A. Gorr, Institute of Veterinary Physiology, Vetsuisse Faculty, University of Zurich, Winterthurerstrasse 260, 8057 Zurich, Switzerland.

## Abstract

Expression of myoglobin (MB), well known as the oxygen storage and transport protein of myocytes, is a novel hallmark of the luminal subtype in breast cancer patients and correlates with better prognosis. The mechanisms by which MB impacts mammary tumorigenesis are hitherto unclear. We aimed to unravel this role, by using CRISPR/Cas9 technology to generate MB-deficient clones of MCF7 and SKBR3 breast cancer cell lines and subsequently characterize them by transcriptomics plus molecular and functional analyses. As main findings, loss of MB, at normoxia, upregulated the expression of cell cyclins and increased cell survival while it prevented apoptosis in MCF7 cells. Also, MB-deficient cells were less sensitive to doxorubicin but not ionizing radiation. Under hypoxia, loss of MB enhanced partial epithelial to mesenchymal transition, thus augmenting the migratory and invasive cell behavior. Notably, in human invasive mammary ductal carcinoma tissues, MB and apoptotic marker levels were positively correlated. In addition, MB protein expression in invasive ductal carcinomas was associated with a positive prognostic value, independent of the known tumor suppressor p53. In conclusion, we provide multiple lines of evidence that endogenous MB in cancer cells by itself exerts novel tumor-suppressive roles through which it can reduce cancer malignancy.

## Introduction

Myoglobin (MB) is well known as the heme-binding globin of heart and skeletal muscles where it is present in the µM (terrestrial mammals) to mM (apnoe diving mammals) concentration range. There, MB functions as a temporary store of O_2_ and buffers short phases of exercise-induced increase in aerobic metabolism, by supplying O_2_ to the myocytes’ mitochondria [1-3]. MB was also reported to scavenge/detoxify harmful reactive oxygen species (ROS) [4] as well as to maintain nitric oxide (NO^•^) homeostasis in cardiomyocytes by either scavenging or producing the radical, in normoxic or hypoxic environments, respectively [5, 6]. Both ROS and NO^•^ are important cell signaling mediators, implying that MB broadly regulates mammalian cell behavior via various cell signaling pathways. Recently, MB was discovered to also be expressed in different tumor types, including multiple epithelial cancers (e,g, breast cancer, prostate cancer, non-small cell lung cancer (NSCLC) etc.) as well as in leukemic bone marrow [7-12]. Being expressed at sub-µM levels in these tumor types, however, it is still unclear whether this low-level abundance is sufficient for the effective oxygenation of cells. While the prominent occurrence of MB in muscle cells is known to prevent apoptosis and to help detoxifying chemotherapeutics such as doxorubicin to non-lethal metabolites [13], these features still have to be demonstrated for malignant tissues. At concentrations that lie perhaps 1000 times below those of myocytes, MB in cancer cells might indeed harbor a set of novel and rather catalytic functions.

A strong hint to a new set of MB functions might be deduced from its ectopic expression site. In healthy breast parenchyma of human subjects, and in the sub-cohort of MB-positive breast and prostate human cancer [8-10], MB was expressed not in the contractile myoepithelial layer but exclusively in the inner secretory luminal epithelium, whose malignant transformation marks the origin of the vast majority of primary human breast carcinomas [14]. Increasing evidence strengthens the notion that oxygenated MB (oxymyoglobin, MBO_2_) contributes to the production and secretion of lipid components into the milk by binding and transporting fatty acids [15, 16], perhaps also in contexts of non-malignant luminal cells, i.e. those of breast tissues in healthy “mice and men” [17]. Beyond the mammary gland, we recently demonstrated that presence of MB in the brown adipose tissue (BAT) of mice appears to link oxygen and lipid based thermogenic metabolism, for example by shifting the lipid droplet (LD) equilibrium towards higher counts of smaller droplets (i.e. towards a browning phenotype) [18]. Importantly, MB in breast epithelia is, despite its low expression level, still of clinical relevance. In mammary carcinomas, MB was found to be expressed in around 40% of invasive ductal carcinomas where it is positively correlated with a higher degree of tumor cell differentiation (luminal subtype), estrogen receptor positivity (ER+), and a significantly better prognostic outcome in ER+ or ER-breast cancer patients [19]. Hence, MB has been validated as a novel luminal marker for breast cancer where it adds to the prognostic value of ER. In prostate cancer, MB was associated with androgen receptor expression, markers of tumor hypoxia and a trend towards a prolonged recurrence-free survival [20]. In agreement with these observations, MB expression was detected in almost every second tumor from human patients with HNSCC, again correlating with favorable overall survival [21]. Conversely, overexpression of MB in NSCLC was associated with poor prognosis [22], suggesting a tumor-type specific role of MB in cancer cells.

At molecular level, the expression of MB in human breast cancer (BrCa) cells is controlled by mitogenic stimuli or oxidative stress [7]. We previously discovered three MB transcripts predominantly expressed in BrCa cells and breast tumors [13], which are under the control of a hypoxia-inducible enhancer/promoter [19] that can be silenced by hormonal treatment [23]. MB’s tumor suppressive biology was underscored by its transcriptional regulation in human BrCa cells [17, 20, 21]. In addition, in LNCaP prostate cancer cells, MB was directly or indirectly involved in induction of apoptosis, TP21 expression, diminished migratory capacity, and possible induction of cell cycle arrest [22]. Yet, in MDA-MB 468 breast cancer cells, MB increased migration [22] and its impact on apoptosis was inconclusive. Whether the opposing observations are attributed to differences in MB expression levels, tumor type or existence of other functional mediators (such as hormonal receptors or p53 status) need to be explored. In the current study, we, therefore, aimed to analyze MB’s role in governing various aspects of cellular vitality and therapeutic responsiveness specifically of breast cancer cells. We designed our experiments to decipher the role of MB in human BrCa cell lines MCF7 and SKBR3. We used CRISPR/Cas9 to generate MB-WT and MB knockout (MB-KO) cells, with which we analyzed the effect of the endogenous MB expression on apoptosis, cell proliferation, migration and invasion, yet, in ways not directly related to the facilitated diffusion or storage of O_2_. The present study provides evidence along with mechanistic insights on the pivotal role played by endogenous MB in tumor suppression and opens the door to a potentially new anti-cancer intervention.

## Materials and Methods

### Cancer cells lines and cell culture

Human MCF-7, epithelial adenocarcinoma-derived breast cancer cells, were obtained from (ATCC #HTB-22) and cultured in Eagle’s Minimum Essential Medium (EMEM) (ATCC #30-2003) supplemented with 10% heat inactivated fetal bovine serum (FBS) (gibco #10270-106) and 1% Penicillin/Streptomycin (gibco #15140-122). Human SKBR3 epithelial adenocarcinoma-derived breast cancer cells were also obtained from (ATCC #HTB-30) and were cultured in Dulbecco’s Modified Eagle Medium (DMEM) (gibco #31885023) supplemented with 1% Glutamax (gibco #35050061), 10% FBS and 1% Pen/Strep. Both cell lines are authenticated by ATCC and the short tandem repeat (STR) profile for either one is as follows: MCF7 (Amelogenin:X; CSF1PO:10; D13S317:11; D16S539:11,12; D5S818:11,12; D7S820:8,9; TH01:6; TPOX:9,12; vWA:14,15) and SKBR3 (Amelogenin:X; CSF1PO:12; D13S317:11,12; D16S539:9; D5S818:9,12; D7S820:9,12; TH01:8,9; TPOX:8,11; vWA:17). For passaging and seeding cells, 0.25% Trypsin-EDTA (gibco #25200-072) was used to detach the adherent cells after washing with warm phosphate buffer saline (PBS). To specifically study the impact of MB in cells adapted to long-term hypoxia, mimicking tumors, the cells were incubated for 72h at hypoxic vs. normoxic conditions. As MB can be either oxygenated or deoxygenated, experiments for cell lines were conducted in room air (normoxia, Nx) and 0.2% O_2_ (severe hypoxia) where MB is mostly deoxygenated [24]. Normoxic (21% O_2_) incubation was performed at standard conditions (37°C, room air with 5% CO_2_) in Heracell 240 incubator (Heraeus #H240-CO_2_). Hypoxic incubations were performed in glove box (Coy Laboratory #8302050) at standard 37°C with 5% CO_2_.

### Generation of CRISPR/Cas9-mediated knockouts of MB

#### Preparation of CRISPR plasmids against human MB

The online “CRISPR Design Tool”, as described by Cong et al., 2015 [25], was used to identify suitable target sites in exon 2 in the human MB gene. Exon 2 of the *MB* gene was selected because it encodes the functionally critical heme pocket residues (E7 distal and F8 proximal histidine residues) [26] (Supplemental Fig. 1A). sgRNAs with the lowest rate of off-target effect score were designed with overhangs (indicated as bold in sequences provided in Supplemental Table 3) to facilitate ligation to CRIPSR/Cas9 plasmids and ordered from Microsynth (Switzerland). These plasmids are based on all-in-one pSpCas9(BB)-2A-Puro (px459) (Addgene #62988) V2.0 and pSpCas9n(BB)-2A-Puro (px462) V2.0 (Addgene #62987 [27]) that express Cas9 protein from *S. pyogenes* and include the scaffold of the sgRNA [27]. They both contain a selection marker for puromycin and offer bacterial resistance to ampicillin. px459 expresses the wild type gene for the Cas9 nuclease and therefore only one plasmid, containing one sgRNA is needed to lead Cas9 to the target sequence and introduce a double strand break. px462 on the other hand, expresses the Cas9n mutant, which does not introduce double strand breaks but nicks one strand of DNA. Here two plasmids with two different sgRNA are required in order to guide Cas9n to two different target sites to achieve a double strand break. sgRNA oligonucleotides were ligated to plasmids using T4 DNA ligase (Promega #M1801) and BbsI restriction enzymes (NEB #R0539L) according to manufacturer. Ligated plasmids were cloned into 5-a competent *E. coli* (NEB #C2987I) cultivated on LB-selection plates with ampicillin (1:1000) (Sigma Aldrich). PCR using U6 primer (annealing to vector backbone) and anti-sense sgRNA (annealing to cloned insert) was performed to identify bacterial colonies that successfully incorporated ligated and not empty plasmids. These specific colonies were amplified by culturing in liquid LB-medium with ampicillin (1:1000) followed by plasmid isolation using QIAprep Spin Miniprep Kit (Qiagen #27104), according to the manufacturer. Isolated CRISPR plasmids were further double checked for incorporating sgRNAs by i) sequencing (Microsynth, Switzerland) ii) failure to linearize upon digestion with BbsI due to lack of the enzyme’s restriction site.

**Figure 1.**
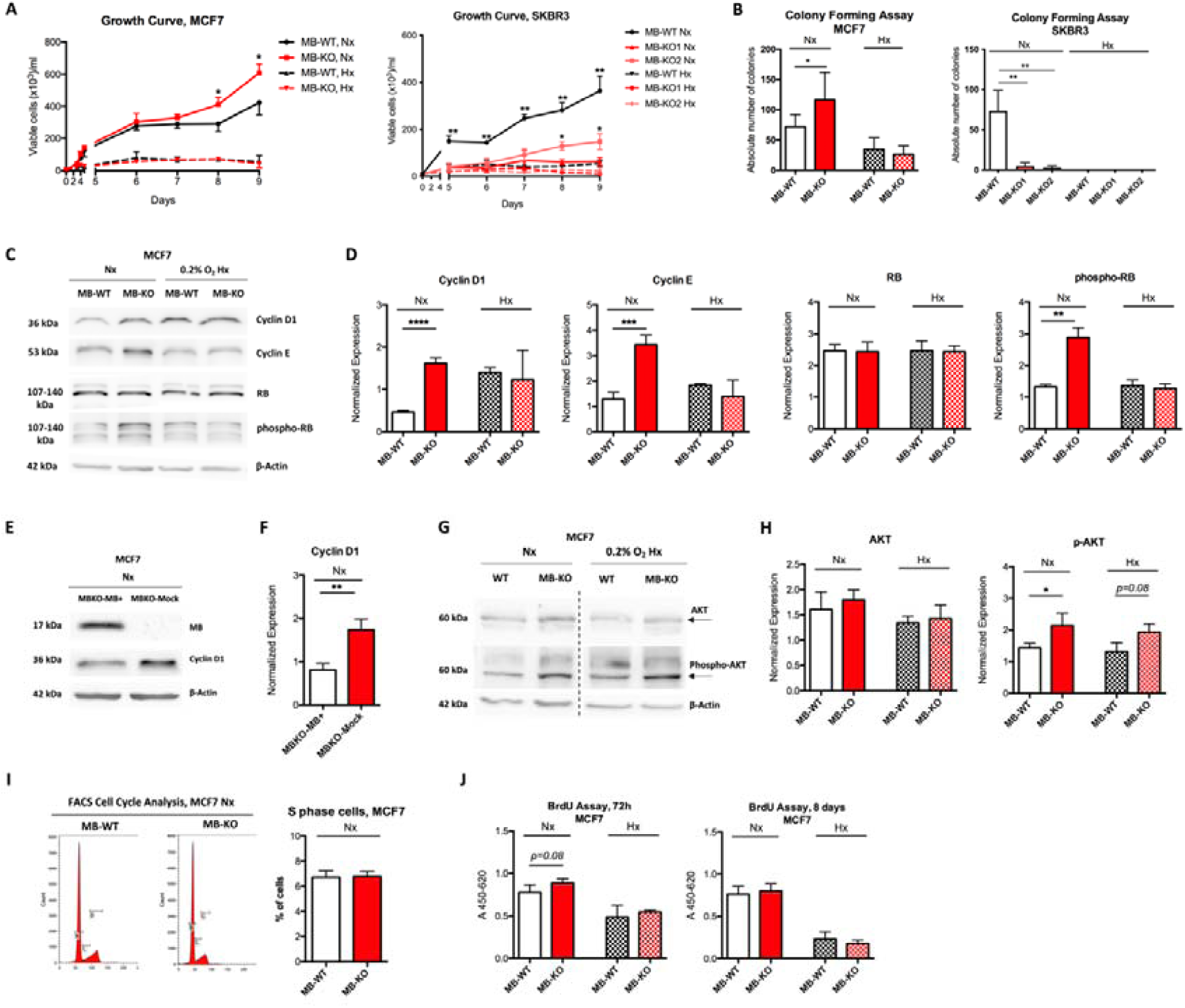
Myoglobin impacts breast cancer cell survival. **(A)** Growth curves of MB-KO vs. MB-WT of MCF7 (left panel) and SKBR3 (right panel) at Nx and 0.2% O_2_ Hx, respectively. Cells were stained by trypan blue, dead cells defined as blue stained were excluded from counting. N=3. **(B)** Colony assay performed using MB-WT and MB-KO clones of MCF7 (left panel) and SKBR3 (right panel). Cells were seeded at low confluency and incubated in normoxia (Nx) or hypoxia (Hx, 0.2% O2) for 9 days. Colonies (>50 cells) stained by crystal violet were counted using a binocular. N=5. (**C**) Representative Western Blotting image (n= 4) of whole tissue lysate of MB-WT and MB-KO clones of MCF7 cells cultured at Nx or 0.2%O_2_ Hx for 72h and stained for cyclin-D1 (33 kDa), cyclin E (53 kDa), retinoblastoma (RB) (130 kDa), 16phospho-retinoblastoma (p-RB) (130 kDa) and β-actin (42 kDa) used as loading control. (**D**) Band intensity of cyclin D1, cyclin E, RB and 16phospho-RB proteins after Western Blotting, from MB-WT (white) and MB-KO (red) MCF7 cells at Nx (empty bars) and 0.2%O_2_ Hx (dashed bars), was quantified using MCID Analysis 7.0 and normalized to β-actin (n=4). (**E**) Representative Western Blotting image (n= 3) of whole tissue lysate of MCF7 MB-KO cells transfected with MB+ or mCherry plasmids and stained for cyclin-D1 (33 kDa) and β-actin (42 kDa) used as loading control. (**F**) Band intensity of cyclin D1 after Western Blotting, from MB-KO transfected with MB+ (white) or mCherry plasmids (red) cells at Nx was quantified using MCID Analysis 7.0 and normalized to β-actin (n=3). (**G**) Representative Western Blotting image (n= 3) of whole tissue lysate of MB-WT and MB-KO clones of MCF7 cells cultured at Nx or 0.2% O_2_ Hx for 72h and stained for AKT (60 kDa 17phospho-AKT (p-AKT) (60 kDa), and β-actin (42 kDa) used as loading control. (**H**) Band intensity of AKT and 17phospho-AKT after Western Blotting, from MB-WT (white) and MB-KO (red) cells at Nx (empty bars) and 0.2%O_2_ Hx (dashed bars), was quantified using MCID Analysis 7.0 and normalized to β-actin (n=3). (**I**) FACS analysis of cell cycle status of MCF7 cells (B marks G1 phase cells, C marks S phase and D marks G2/M phase cells) after culturing at Nx for 72h and staining with Pyronin-Y and Hoechst stains and quantification of proportions of cells at S phase (n=3). (**J**) BrdU incorporation assay of MB-WT (white) and MB-KO (red) clones of MCF7 cells cultured at Nx (empty bars) or 0.2% O_2_ Hx (dashed bars) for 72h or 8 days. Absorbance of clones with anti-BrdU antibody for 4.5h, measured at 450nm and corrected at 620nm (n=4). Data are presented as mean and standard error of mean and analyzed by Student’s-t test except SKBR3 data in panel (**B**) by One-way ANOVA with Tukey’s multiple comparison post-hoc analysis *p<0.05, **p<0.01, ***p<0.001

### Generation of MB knockout clones of MCF7 and SKBR3 cells

We generated single cell clones for generating MB knock out cells with the purpose of exploiting cell models with identical genetic background that only differ in MB expression. The single cell clones were selected for the CRISPR-knock out approach based on their highest basal as well as the hypoxia-regulated expression of MB (Supplemental Fig. 1B and C, red square mark). These clones were further tested for transfectability using pmCherry-N1 control vectors (Takara Bio #632523). 200’000 cells of selected monocolonies of either cell line were seeded in 6-well plates and transfected on next day with CRISPR plasmids using Lipofectamine 2000 (Invitrogen by Life Technologies #11668019) according to manufacturer instructions. 24h post-transfection, cells were selected using 2M puromycin for 48 h for MCF7 and SKBR3. Surviving cells were seeded on single cell bases for screening to identify knockout monoclonal cells. DNA was extracted using homogenization buffer (50mM KCl, 10mM Tris/Hcl pH 8.3, 10mM Gelatin, 0.045% NP-40, 0.045% Tween 20) with 50mM proteinase K (Sigma Aldrich #3115879001). PCR was conducted using primers that flank exon 2 of the human MB gene (Supplemental Table 3) and products were screened by running on 10% PAGE against that of control wildtype clone. Clones that showed multiple bands, indicating heteroduplexes, were sequenced and further analyzed using the online TIDE tool [28]. This tool precisely determines the spectrum and frequency of targeted mutations (insertions or deletions) [29]. Candidate clones that showed no trace of a wildtype allele presence were further inspected for separate allelic modifications by cloning of purified aforementioned PCR product into pGEM-T vectors (Promega #A1360) and transformation to NEB 5-a competent *E. coli* according to manufacturer instructions. Cloned plasmids were isolated as described above and sequenced (Microsynth, Switzerland). DNA sequences were again analyzed by TIDE tool and aligned to the wildtype MB gene sequence (NCBI nucleotide blast). The resulting modifications in the protein sequence were analyzed with the Expasy Translation Tool [30] and NCBI protein blast.

### Total Protein Extraction and Western Blotting

Cells were lysed in RIPA buffer (50 mM Tris/Hcl pH 8, 150 mM NaCl, 1% NP-40, 0.5% Na deoxycholate, 1 mM EDTA, 0.1% SDS) in the presence of Protease Inhibitor Cocktail Set III, EDTA-Free, according to the manufacturer instructions (Merck millipore, #539134). Protein concentration was determined using the Pierce BCA assay (Thermo Scientific, #23228, #23224). SDS-PAGE (Bio-Rad, USA) was used for separation of proteins which were transferred to nitrocellulose blotting membrane (GE Healthcare, #10600002). Following washing with 0.05% Tris buffered saline-Tween (TBST), membranes were blocked in 5% skimmed milk (or 5% FBS for phospho-proteins) in 0.05% TBST for 1 hour at room temp. Next, membranes were incubated at 4°C overnight with following antibodies: rabbit anti-myoglobin (FL-154) polyclonal antibody (Santa Cruz, #sc-25607) 1:200; mouse anti-p53 (DO-1) monoclonal antibody (Santa Cruz, #sc-126) 1:200; mouse anti-cyclin D1 monoclonal antibody (Santa Cruz, #sc-8396) 1:200; mouse anti-cyclin E (HE12) monoclonal antibody (Santa Cruz, #sc-247) 1:200; mouse anti-Rb (IF8) monoclonal antibody (Santa Cruz, #sc-102) 1:200; mouse anti-Phospho-Rb (B-4) monoclonal antibody (Santa Cruz, #sc-514031) 1:200; mouse anti-ERα (F-10) monoclonal antibody (Santa Cruz, #sc-8002) 1:200; rabbit anti-PI3K p85 polyclonal antibody (Cell Signaling, #4292) 1:1000; rabbit anti-AKT polyclonal antibody (Cell Signaling, #9272) 1:1000; rabbit anti-Phospho-AKT (Ser473) polyclonal antibody (Cell Signaling, #9271) 1:1000; rabbit anti-Smad2/3 polyclonal antibody (Cell Signaling, #5678) 1:1000; rabbit anti-Phospho-Smad2 (Ser245/250/255) polyclonal antibody (Cell Signaling, #3104) 1:1000; rabbit anti-Phospho-Smad1 (Ser463/465)/Smad5 (Ser463/465)/ Smad9 (Ser465/467) (D5B10) monoclonal antibody (Cell Signaling, #13820) 1:1000; mouse anti-E-Cadherin (36) monoclonal antibody (BD Biosciences, #610182) 1:5000; mouse anti-HIF-1α (mgc3) monoclonal antibody (Abcam, #ab16066) 1:1000; rabbit anti-HIF-2α (EPAS1) polyclonal antibody (Novus, #NB100-122) 1:1000; rabbit anti-AIF polyclonal antibody (Abcam, #ab1998) 1:1000; rabbit anti-VDAC1/Porin polyclonal antibody (Abcam, #ab15895) 1:1000; rabbit anti-JNK Pan Specific polyclonal antibody (R&D, #AF1387) 0.2 ug/ml; mouse anti-ATG5 (7C6) monoclonal antibody (nanoTools, #0262) 0.5 ug/ml; mouse anti-LC3 (5F10) monoclonal antibody (nanoTools, #0231) 0.5 ug/ml; mouse anti-p27 (F-8) monoclonal antibody (Santa Cruz, #sc-1641) 1:200; mouse anti-p21 (F-5) monoclonal antibody (Santa Cruz, #sc-6246) 1:200; mouse anti-cyclin A (H-3) monoclonal antibody (Santa Cruz, #sc-271645) 1:100; mouse anti-PCNA (PC10) monoclonal antibody (Santa Cruz, #sc-56) 1:200; mouse anti-beta-actin monoclonal antibody (Sigma Aldrich, #A5441) 1:5000. After washing, membranes were incubated with HRP-conjugated secondary antibodies: donkey-anti-rabbit 1:5000 (Amersham, #NA934V) or goat-anti-mouse 1:5000 (Santa Cruz, #sc-2031), for 1 hour at room temp. After washing, bands were visualized by Fujifilm LAS-3000 Chemiluminescent imager using SuperSignal^TM^ West Femto (Thermo Scientific, #34095). For quantification purposes, bands peak intensity was analyzed by MCID Analysis 7.0 software.

### RNA extraction and real time PCR

Cultured cells were washed twice with 1x PBS followed by lysis and RNA extraction using ReliaPrep RNA Cell Miniprep System (Promega, #Z6011), according to the manufacturer. First strand cDNA was synthesized following the manufacturers protocol using RevertAid First Strand cDNA Synthesis Kit (Thermo Scientific, #K1622). A final cDNA concentration of 5ng/µl was used for semi-quantitative real time PCR analysis performed in Thermocycler ABI7500 Fast (Applied biosystems), using PowerUp SYBR Green Master Mix (Applied biosystems, #A25778). Primers were designed with Primer3 and are listed in (Supplemental Table 3). Primers were validated first by qRT-PCR via i) melting curve analyses (mode integrated in the 7500 Fast Real-Time PCR System to confirm having one single peak corresponds to desired product) as well as ii) on acrylamide gels to confirm size and purity of PCR products iii) Sanger sequencing of some of PCR products. DDCt method was used to calculate mRNA expression levels [31, 32].

### Trypan Blue Viability assay

In 6-well plate scale, 10’000 cells were seeded in duplicates using 10% FBS containing culture medium. The next morning, cells were incubated at 21% or 0.2% O_2_. From day 3 to 8, cells were trypsinized, then mixed with 0.4% trypan blue solution in PBS (Sigma Aldrich #T6146) in a 1:1 ratio and counted by hemocytometer (Neubau Chamber). Only viable cells (white) were counted and plotted as growth curve over a time period of 9 days while dead cells (blue) were excluded.

### Colony forming assay

The clonogenic assay was used to evaluate reproductive viability and survival of cells, so the mitotic capacity of a single cell at different conditions and treatments was assessed [33]. 250 cells were seeded in 20% FBS containing culture medium in either 6-well plate (for Dox or TAM treatment) or T25 angled neck culture flask with vent/close cap (TPP #90025) (for radiation treatment). The next morning, 5 nM or 10 nM of doxorubicin as well as 2.5 mM of tamoxifen was added, and cells were incubated at 21% or 0.2% O_2_ environments for 9 days. Solvent treated cells were used as a control. After overnight incubation, caps were closed and all flasks including untreated controls were transferred to the irradiation facility. Normoxic or hypoxic cells designated to irradiation were treated with 3Gy of radiation. Flasks were moved back to 21% or 0.2% O_2_ incubators where they were vented and incubated for 9 days. Formed colonies were washed and then fixed/stained using a mixture of 0.5% crystal violet and 6% glutaraldehyde overnight at room temperature. Plates or flasks were then washed with water several times and then air dried. Counted colonies were defined as aggregates of at least 50 cells. Number of colonies were normalized to those of untreated controls.

### BrdU proliferation assay

The BrdU assay was performed using the Cell Proliferation ELISA BrdU assay (Roche #11647229001) according to manufacturer. Briefly, 20’000 cells/well were cultured in triplicates, in 96-well pates using complete culture medium and incubated for 72 h at 21% normoxic or 0.2% O_2_ hypoxic environments. After incubation, 10 mM of BrdU labelling solution was added and cells were incubated for 4.5 h at 37°C. Labelling medium was removed and cells were fixed, followed by staining with anti-BrdU-POD working solution for 90 min at room temperature. Wells were rinsed three times with 1X PBS followed by color development by adding substrate and 1 M H_2_SO_4_ stop solution. Photometric detection was carried out, within 5 min of adding stop solution, at 450 nm (reference wavelength is 690 nm) by Multiskan RC plate reader (Thermo Labsystems).

### MTT assay

The MTT (Methyl thiazolyl diphenyl-tetrazolium bromide) reduction assay combines the measurement of cell proliferation with the assessment of mitochondria activity. Mitochondrial dehydrogenases of only viable cells will convert MTT to formazan product within the mitochondria. 20’000 cells/well were cultured, in triplicates, in 96-well plates and incubated overnight. Cells were incubated at 21% or 0.2% O_2_ with or without (0.05, 0.1 and 0.3 uM) doxorubicin (Dox) for 72 h. 500 mM of freshly prepared MTT reagent (Sigma Aldrich #M5655) was added to each well and incubated for 4h at 37°C in the dark. Medium was removed and 100µl of dimethyl sulfoxide (DMSO, Sigma Aldrich) solvent was added to dissolve purple precipitate of formazan. Absorbance was measured at 570 nm (reference wavelength is 620 nm) by a Multiskan RC plate reader (Thermo Labsystems).

### Migration assay

Migratory capacity of cancer cells was assessed by scratch assay. In 6-well plates, 1.2 million cells were cultured per well, in duplicates, in serum-reduced medium to minimize proliferation and hence observe migration only. After overnight incubation, a wound was swiftly created into the cells confluent monolayer and cells were incubated at 21% or 0.2% O_2_ for 48 h. Pictures were taken from the same 3 spots (predefined by reference lines drawn on the bottom of the plates) at 0, 24 and 48 h from wound generation. Images were captured using an inverted Axiocam HR microscope coupled with a CCD camera (10x magnification) and analyzed by ImageJ software. The area of the wound was measured and the average of the 3 images at each time point was calculated. Scatter plot was created as % of normalized wound area as function of time of wound generation.

### Cell cycle status analysis

Protocol was adapted from Kim and Sederstrom [34]. It differentiates resting (G0) from proliferating cells by determining total RNA content of cells, as G0 cells have much lower levels of RNA compared to (G1-S-G2-M) cells. In presence of Hoechst 33342, which stains double stranded DNA, and Pyronin Y, which exclusively stains RNA, both signals were quantified by flow cytometry (FACS) analysis. Briefly, 1 million cells were incubated in 10-cm plates, in duplicates, overnight in serum free culture medium in order to align cells’ cycle status. After 72 h of incubation in 10% FBS culture medium, cells were harvested, washed and fixed by adding pre-chilled 70% ethanol dropwise while vortexing, followed by 2h incubation at -20. Following multiple cycles of washing and centrifuging, cells were stained using 2mM Hoechst 33342 (Sigma Aldrich #14533) and 4mM Pyronin Y (Sigma Aldrich #83200) double staining solution. Samples were incubated in dark for 20 min followed by fluorescence analysis by Gallios Flow Cytometer 561 ready (Beckman Coulter). UV (355 nm) and blue (488 nm) lasers were used for excitation as well as 461 nm and 575 nm filter set for Hoechst 33342 and Pyronin Y, respectively.

### Annexin-V and NAO-based FACS analysis of apoptosis

A key event in the development of apoptosis is oxidation of the mitochondrial inner membrane key lipid cardiolipin (CL), followed by the lipids externalization to the outer mitochondrial membrane (OMM), which is subsequently followed by the release of cytochrome c from the mitochondria into the cytoplasm. 10-N-Nonyl acridine orange (NAO), a fluorophore that forms a stable complex selectively with reduced form of CL, but not with oxidized CL, can be used for flow cytometry analysis of the extent of surface of the inner mitochondrial membrane (IMM) [35]. Therefore, cell populations with negative NAO fluorescence correspond to the population of apoptotic cells, i.e. those cells whose pool of CL exists as oxidized lipid in the OMM [36] As second apoptosis assay, we used the conventional Annexin V staining, which binds to phosphatidylserine exposed in the outer membrane that precedes the loss of membrane integrity which accompanies the later stages of cell death. The stain was either accompanied by live/dead stain of propidium iodide (PI). 100’000 cells were incubated, in duplicates, at 21% O_2_ for 72 h, followed by incubation with 1 mM of the apoptosis inducing agent staurosporin (STS) for 6 h. Cells were harvested, washed and stained for either 100 nM NAO (Sigma Aldrich) (for 30 min in dark at 37°C) or Annexin V-FITC (Promokine #PK-CA577-K101-100) (5 ml for 5 min in dark at room temperature), followed by fluorescence analysis (Ex = 488 nm; Em = 530 nm) using Gallios Flow Cytometer 561 ready (Beckman Coulter).

### Invasion assay

The assay protocol for MCF7 cells was adapted from [37]. 50 ml of Matrigel basement membrane matrix (Qiagen #354234) prepared in serum free culture medium (1:8 ratio to reach final concentration of 1.2 mg/ml) was used to coat 8µm PET membrane Transwell inserts in 24-well plates (Corning, #3464) to mimic extracellular matrix. The Transwell inserts were incubated with Matrigel at 37°C overnight to solidify. 100’000 cells in reduced serum culture medium were placed in upper chamber and incubated for 48 h at 21% or 0.2% O_2_, with 10% serum medium placed in the lower chamber as chemoattractant. Inserts were then washed twice with 1X PBS then cells were fixed and stained simultaneously in mixture of 0.5% crystal violet and 6% glutaraldehyde overnight at room temperature. Inserts were washed with water several times and then air dried. Cells on upper surface of the membrane (non-invading cells) were removed by moistened cotton swap and invading cells exist on the lower surface were imaged by Digital Microscope VHX-6000 (Keyence, United States). 5 random pictures were captured from each insert and crystal violet signal was quantified using ImageJ software.

### ROS measurement

The DHE (Dihydroethidium) assay kit (abcam #ab236206) was used to measure ROS directly in living cells. The kit is specific for superoxide and hydrogen peroxide and was used according to the manufacturer’s instructions. 10’000 cells/well were seeded, in triplicates, in 96-well plate and incubated at 21% or 0.2% O_2_ for 72 h. Antimycin A, an inhibitor of complex III of the mitochondrial electron transport chain, was included as a positive control for ROS generation. N-acetyl Cysteine was included as an antioxidant negative control. Fluorescence using excitation/emission wavelength of 500/590 was measured by Infinite 200 Pro fluorescent multi-chromate plate reader (Tecan, Switzerland).

### Immunofluorescence

50’000 cells were cultured on cover slips placed in 24-well plates and incubated at standard culture conditions overnight. After attaching, plates were incubated at 21% or 0.2% O_2_ for 72 h. Cells were fixed in 4% paraformaldehyde then washed and permeabilized in 0.1% triton/PBS followed by washing. Cells were then blocked in 10% normal goat serum (NGS) in PBS for 1h at room temperature followed by staining with: mouse anti-ERα (F-10) monoclonal antibody (Santa Cruz, #sc-8002) 1:50; mouse anti-HIF-1α (mgc3) monoclonal antibody (Abcam, #ab16066) 1:200; rabbit anti-HIF-2α (EPAS1) polyclonal antibody (Novus, #NB100-122) 1:200; rabbit anti-AIF polyclonal antibody (Abcam, #ab1998) 1ug/ml, antibodies in 3% NGS in PBS overnight at 4°C in wet chamber. Cover slips were then washed in PBS-Tween and incubated with Alexa 546-a-mouse secondary antibody (Thermo Fisher) 1:500 in 3% NGS in PBS, and counterstained by DAPI (Sigma Aldrich) 1:1000 in PBS for 1h at room temperature. Slides were mounted using Fluorescent Mounting Medium (Dako #S3023), sealed with nail polish and kept at 4°C in the dark. Images were then taken using a Zeiss Imager Z2 fluorescent microscope coupled with a CCD camera.

### Myoglobin expression

RNA from MDA-MB468 cells was extracted with the RNeasy mini kit (Qiagen) and converted to cDNA with Superscript III (Thermo Scientific). The CDS of the MB gene was amplified with the KAPA HiFi PCR Kit (Peqlab) and flanked with restriction enzyme target sites. PCR products were cleaned with the Wizard SV Gel and PCR Clean Up Kit (Promega) and cut with XhoI and BamHI or HindIII und BamHI (Thermo Scientific). After vector dephosphorylation, MB CDS were ligated in XhoI and BamHI digested pmCherry-C1 (Takara Bio #632524) or in HindIII and BamHI digested pmCherry-N1 (Takara Bio #632523) with T4 DNA ligase (Thermo scientific). Plasmids were transformed into DH10B cells by electroporation and plated on 50mg/µl Kanamycin agar. Single clones were amplified in LB media, isolated with the Gene JET Plasmid Miniprep Kit (Thermo Scientific) and approved by Sanger sequencing (Starseq). For overexpression experiments, cells were seeded in 12-well plates to reach 70% confluency on the next day. For each well, 3.3 μg DNA of either plasmids was added to 150 μl Opti-MEM reduced serum media (Gibco #31985062) then mixed thoroughly with 9.9 ul Fugene HD transfection reagent (Promega #E2311). After 10 mins, 150 μl of the complex was added to each well and left for 6 h before changing medium with complete growth medium and incubation at 21%, 1% and 0.2% O_2_ for 48 h. Whole cell lysates and RNA were subsequently isolated from all clones. Pictures were taken on days 1 and 2, to confirm successful transfection.

### RNA sequencing

#### RNA quality control and sequencing

RNA was quality-checked on an Agilent RNA nano bioanalyzer, yielding RNA integrity numbers between 9.6 and 10. RNA was quantified with the Qubit RNA BR Assay (Thermo Fischer). Illumina sequencing libraries were generated with the NEB Next Ultra II directional RNA library prep Illumina kit. 50 bp single-end Illumina sequencing was performed on an Illumina HiSeq2500 sequencing platform (NGS core facility, University Mainz, Germany). Prior to RNA-Seq, clone authenticity was approved by Sanger Sequencing (Starseq, Mainz).

#### Mapping against reference genome

RNA-Seq data were filtered for adapter sequences and trimmed to a quality score greater 20 with the program package bbduk (BBTools suite; https://sourceforge.net/projects/bbmap/). Reads shorter than 20bp were discarded. The genome index file was created in STAR version 2.7.3a [38] with a sjdbOverhang of 38 using the aforementioned reference GRCh38 and the annotation release 98 (http://ftp.ensembl.org/pub/release-98/gtf/homo_sapiens/Homo_sapiens.GRCh38.98.gtf.gz). The cleaned reads were mapped with the STAR aligner version 2.7.3a [38] to the human genome GRCh38, downloaded from ENSEMBL (ftp://ftp.ensembl.org/pub/current_fasta/homo_sapiens/dna/ftp://ftp.ensembl.org/pub/current_fasta/homo_ sapiens/dna/Homo_sapiens.GRCh38.dna.primary_assembly.fa.gz). A multimap filter was applied with a cut-off of 20, the maximum mismatch allowance was set to 0.04. The minimum overhang for non-annotated splice junctions was set to 8, the minimum overhang for annotated splice junctions was set to 1. Count data files were generated from each mapping to determine differential gene expression between samples with DeSeq2, which is part of the Bioconductor package for R [39]. All R analyses were done in RStudio Version 1.0.153. Values were normalized using the rlog function and converted to log2 values.

#### Gene ontology analyses

The Webtool g:Profiler [40] and corresponding Ensembl IDs were applied for the over-representation analysis (ORA) of differentially expressed genes (DEGs) with an adjusted p-value <0.05. The analysis was performed using the Gene Ontology (GO) biological process and KEGG pathways database [41], yielding a Generic Enrichment Map and a Gene Matrix Transposed file. Both were processed with the EnrichmentMap Cytoscape App 3.3.1 in Cytoscape 3.8.2. The Network was created with an edge Cutoff at a Jaccard-Index of 0.25. The resulting Network was MCL clustered with clusterMaker2 Cytoscape App 1.3.1 with the similarity coefficient for edge weight and annotated with the AutoAnnotate Cytoscape App 1.3.3. For clustering, Cluster was selected (part of the). CoSE (Compound Spring Embedded) was chosen as the layout algorithm and the network was scaled for better distribution. For an additional visualization of the ORA results, the Enrichment fold change was calculated using the GMT file downloaded from the g:Profiler website as the background list. The percentage of differentially expressed genes (DEGs) associated with a specific term was divided by the percentage of background list genes with the same term. The results were plotted in R using ggplot2.

### RNA-Seq data will be available at EBI, project number PRJEB51094

Human cancer tissue array staining was subjected to antigen retrieval at 98 °C for 20 min in EDTA buffer (pH 9.0) in a steamer and incubated with a rabbit anti-cleaved caspase 3 antibody (Cell Signaling, 9664L) 1:150 overnight at 4 °C followed by anti-rabbit 594 (Invitrogen A11012) 1:400 for 1h at room temp. Subsequently the array was incubated with a monoclonal rabbit anti-MB (Abcam, ab77232) 1:400 for 1h at room temp. followed by anti-Rb 488 (Invitrogen, A11008) 1:500 for 1h at room temp. and DAPI for 15 min. Slides were fully scanned (NanoZoomer 2.0-HT; Hamamatsu, Hamamatsu City, Japan) and images of individual cores were captured. For immunohistochemical analysis of p53 our study included tissue microarrays of invasive breast cancer of patients diagnosed at the Institute of Surgical Pathology (University Hospital, Zurich, Switzerland), as described [8]. Tissue sections were processed using automated immunohistochemistry platforms (BOND, Labvision (Fremont, CA, USA) and Benchmark, Ventana (Tucson, AZ, USA)) using anti-p53 (DO7) antibody and scanned for further evaluation.

## Results

We generated MB-knockout (MB-KO) breast cancer cells of the MCF7 and SKBR3 human cell line and compared their molecular and functional profiles to those of matching MB-wildtype (MB-WT) cells as a control. The two breast cancer cell lines were chosen for studying the tumor-specific effects of MB in a p53 wildtype / ER-positive (MCF7) and a p53 gain-of-function (GoF) / ER-negative (SKBR3) background, respectively. As detailed in material and methods, all KO constructs were based on single cell clones with the purpose of exploiting cell models with identical genetic background that only differ in *MB* with specific knockout hit in each allele of the *MB* gene on genomic DNA and the expected resultant truncated protein (Supplemental Fig 2). We detected no MB protein expression by Western blotting in both MB knock out clones (Supplemental Fig 1B and C). In wildtype controls, MB protein was upregulated during prolonged (72h) severe (0.2% O_2_) hypoxia in MCF7 but not in SKBR3 WT cells.

**Figure 2.**
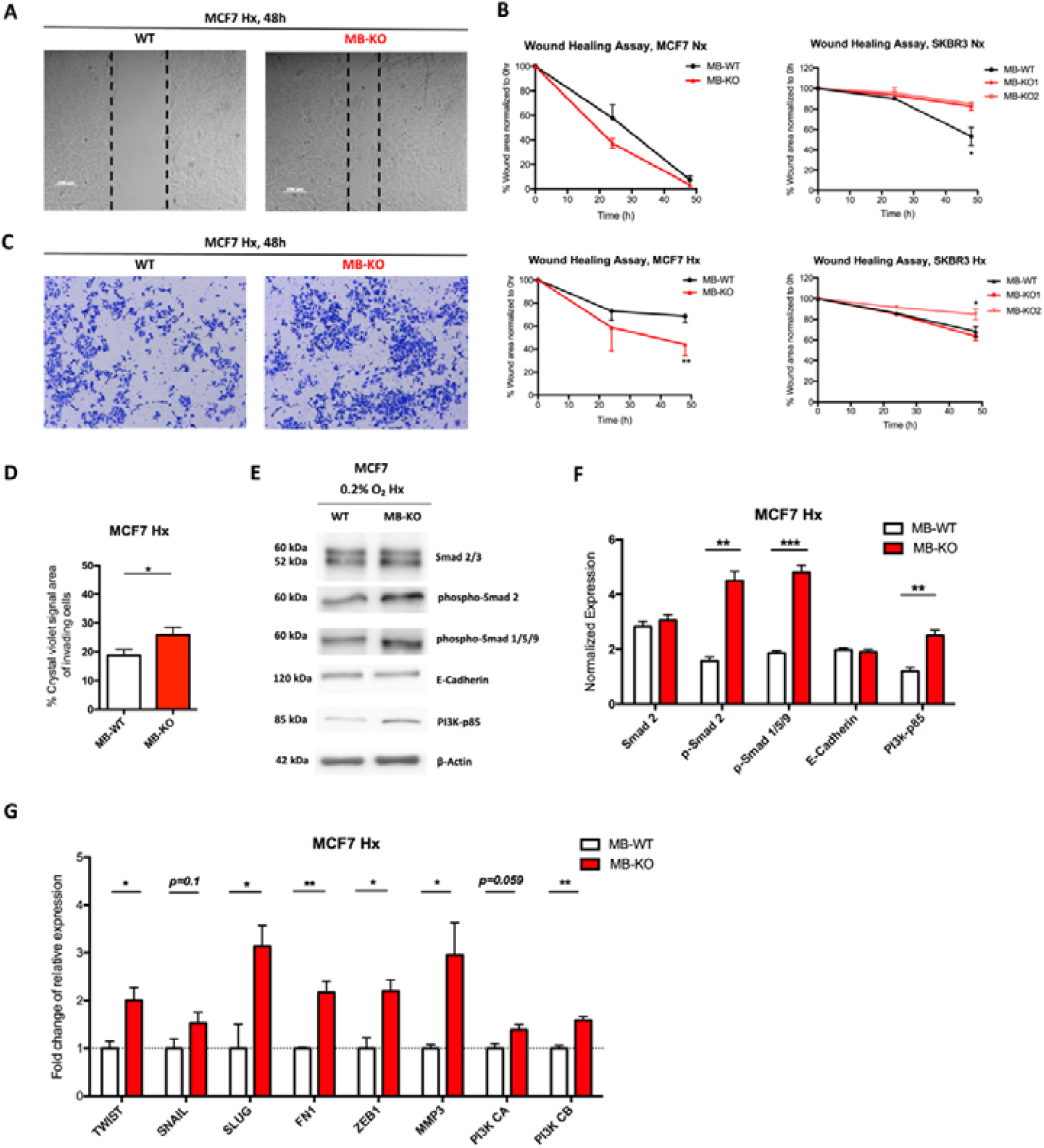
Loss of myoglobin increases migratory capacity of hypoxic MCF7 cancer cells. (**A**) Representative images of Wound healing assay (n= 3) for MCF7 clones grown at 0.2% O_2_ Hx in serum reduced medium for 48h. A scratch was initiated into a monolayer of each cell line and cells were incubated in normoxia (Nx) or hypoxia (Hx, 0.2% O_2_). The % of scratch area was measured and calculated after 24 and 48h. (**B**) Scatter plot of % scratch area (normalized to 0h) against time of incubation. MCF7 (left panels) and SKBR3 (right panels) wild type (MB-WT,white) and MB knockout (MB-KO, red) clones (n=3). (**C**) Representative images of Transwell invasion assay showing only invading wildtype and MB-KO MCF7 cells through layer of Matrigel after culturing at 0.2% O_2_ Hx in Transwell for 48h then fixed/stained with 0.6% glutaraldehyde and crystal violet solution. (**D**) Quantitative representation of % of crystal violet signal area of invading cells of MB-WT (white) and MB-KO (red) of MCF7. 5 random images from each well were captured and analyzed by ImageJ and average of signal area was calculated (n=3). (**E**) Representative Western Blotting image (n= 3) of whole tissue lysate of MB-WT and MB-KO clones of MCF7 cells cultured at 0.2% O_2_ Hx for 72h and stained for Smad2/3 (60 and 52 kDa, respectively 19phospho-Smad2 (60 kDa 19phospho-Smad 1/5/9 (60 kDa), E-Cadherin (120 kDa), PI3K-p85 (85 kDa) and β-actin (42 kDa) used as loading control. (**F**) Band intensity of Smad2, 19phospho-Smad2 19phospho-Smad 1/5/9, E-Cadherin and PI3K-p85 proteins after Western Blotting, from MB-WT (white bars) and MB-KO (red bars) cells at 0.2% O_2_ Hx was quantified using MCID Analysis 7.0 and normalized to β-actin (n=3). (**G**) Relative mRNA expression levels of genes that regulate epithelial to mesenchymal transition: *TWIST, SNAIL, SLUG, FN1, ZEB1, MMP3, PI3K CA and Pi3K CB*, quantified by qPCR and normalized to β-actin (*ACTB*) mRNA expression levels, from MB-WT (white) and MB-KO (red) clones of MCF7 cells cultured at 0.2% O_2_ Hx for 72h. (n=3 per group). Data are shown as bar graph with mean and standard error of mean and analyzed by Student’s t-test *p<0.05; **p<0.01, ***p<0.001.

### Myoglobin impacts breast cancer cell survival

Whether endogenous, rather than overexpressed, MB expression helps regulating BrCa cell survival has never been investigated so far. To answer this, we analyzed the growth of cancer cell clones in normoxic (Nx, air atmosphere) and hypoxic (Hx, atmosphere with 0.2% O_2_) cultures using the Trypan blue viability exclusion assay. Nx, but not Hx, MB-KO MCF7 cells grew 33% faster than WT cells (Fig. 1A), suggesting oxygenated MB (MBO_2_) to either slow down cell proliferation or to reduce survival of cancer cells. In contrast, the loss of MB in SKBR3 cells lowered proliferation/survival under both, normoxic and hypoxic conditions (Fig. 1A). In addition, a clonogenic assay supported the results obtained from growth curves. Under Nx conditions, the loss of MB increased the number of colonies by 163% in MCF7 cells while lowered it by 680% in SKBR3 cells (Fig. 1B). Due to the proliferative arrest of cells under severe hypoxia [23], far less colonies were formed at 0.2% O_2_ than under Nx conditions. We thus concluded that loss of MB increased MCF7, but decreased SKBR3, cell viability and potentially proliferation. To more specifically investigate the beneficial impact of MB basal expression on limiting cancer cell proliferation, we analyzed the expression of cyclin D1 and cyclin E in MCF7 cells, both regulating the cell division process in murine mammary tumors [42]. The loss of MB in Nx MCF7 cells increased expression of cyclin D1 by 3.2 and cyclin E by 2.9 folds, respectively, (Fig.1C and D) correlating with an increased phosphorylation of retinoblastoma (RB) protein (Fig.1D). In contrast, the loss of MB in Hx MCF7 cells did not affect expression of cyclin D1 or E (Fig. 1C and D). Re-expressing MB in Nx KO cells resulted in diminished cyclin D1 expression (Fig.1 E and F) confirming MB-aided regulation of this specific cell cycle protein. As the expression of p27 and p21 proteins (Supplemental Fig. 3A-C) as well as cyclin A or PCNA (Supplemental Fig. 3D-F) was independent of MB expression, these data suggest MBO_2_ to alter the G1/S phase transition independently from the p53 downstream proteins. Although levels of p-AKT were slightly increased in both normoxic and hypoxic MB-KO MCF7 cells (Fig. 1G and H), loss of MB in normoxic MCF7 cells had no influence on the percentage of S phase cells population (Fig. 1I), suggesting that MB expression mainly regulates survival and not proliferation. The unchanged levels of uridine base uptake in normoxic MB-KO MCF7 cells cultured for either 3 or 8 days further supported that MB does not impact proliferation (Fig. 1J). In conclusion our data show that MB lessens the viability/survival, rather than reducing the proliferation, of oxygenated MCF7 cells.

**Figure 3.**
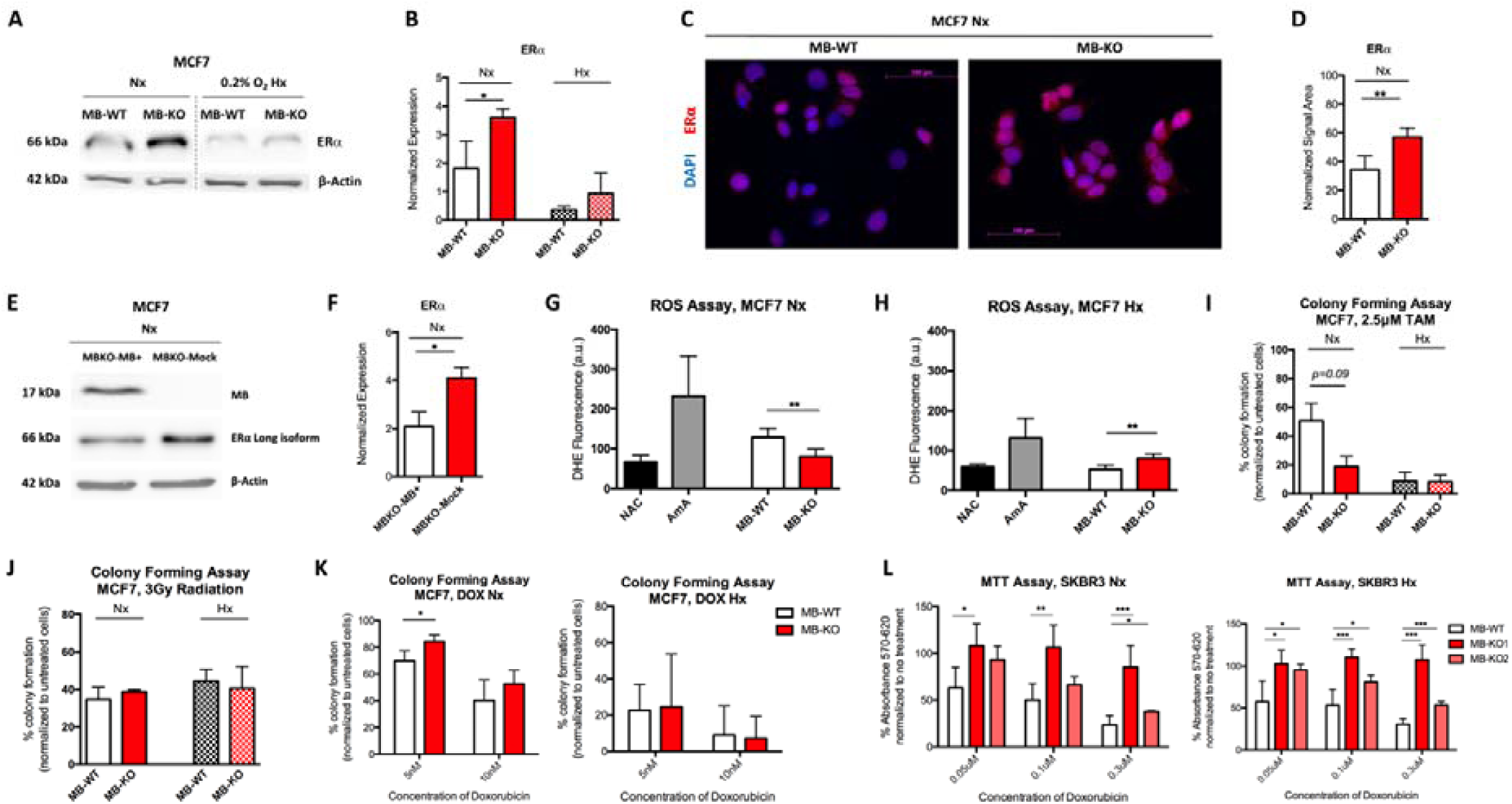
Myoglobin interferes with estrogen receptor expression, ROS generation and response to chemotherapeutic but not irradiation treatment. (**A**) Representative Western Blotting image (n=4) of whole tissue lysate of MB-WT and MB-KO clones of MCF7 cells cultured at normoxia Nx or 0.2% O_2_ Hx for 72h and stained for estrogen receptor alpha ERα (60 kDa) and β-actin (42 kDa) used as loading control. (**B**) Band intensity of ERα protein after Western Blotting, from MB-WT (white) and MB-KO (red) cells of MCF7 at Nx (empty bars) and 0.2%O_2_ Hx (dashed bars), was quantified using MCID Analysis 7.0 and normalized to β-actin (n=4). (**C**) Representative immunocytochemistry images of MB-WT and MB-KO MCF7 cells stained for ERα (red) and DAPI (blue) after culturing at normoxia for 72h. Scale bar is 100µm. (**D**) Quantitative analysis of ERα signal area as a percentage of nuclear signal area (n=4). (**E**) Representative Western Blotting image of whole tissue lysate of MCF7 MB-KO cells transfected with MB+ or mCherry plasmids and stained for estrogen receptor alpha (ERα) (60kDa) and β-actin (42 kDa) used as loading control. (**F**) Band intensity of ERα after Western Blotting, from MB-KO transfected with MB+ (white) or mCherry plasmids (red) cells at Nx was quantified using MCID Analysis 7.0 and normalized to β-actin (n=3). (**G**) and (**H**) Fluorescence assay of dihydroethidium (DHE) to measure ROS directly in living cells cultured at Nx or 0.2% O_2_ Hx for 72h, respectively. 10’000 cells/well were seeded, in triplicates, in 96-well plate and incubated at 21% or 0.2% O_2_ for 72 h. Antimycin A, an inhibitor of complex III of the mitochondrial electron transport chain, was included as a positive control for ROS generation. N-acetyl Cysteine was included as an antioxidant negative control. Fluorescence using excitation/emission wavelength of 500/590 was used (n=3). (**I**) and (**J**) Colony assay using MCF7 MB-WT (white) and MB-KO (red) cells seeded at low confluency and treated with 2.5 µM tamoxifen (TAM) (**H**) or 3Gy irradiation (**I**). Cells were then incubated in Nx (empty bars) or 0.2% O_2_ Hx (dashed bars) for 9 days before colonies (>50cells) were stained and counted (n= 4). Data are presented as % of colony forming normalized to untreated. (**K**) Colony assay using MCF7 MB-WT (white) and MB-KO (red) cells seeded at low confluency and treated with 5nM or 10nM Doxorubicin. Cells were then incubated at Nx or 0.2% O_2_ Hx for 9 days before colonies (>50cells) were stained and counted (n= 4). (**L**) MTT assay using SKBR3 clones. Cells were incubated in normoxia (Nx, left panel) or hypoxia (Hx, 0.2% O_2_, right panel) with 0.05, 0.1 and 0.3 µM doxorubicin for 72h. After the addition of MTT, absorbance was measured and normalized to cell number (determined by growth curve experiments) (n=4-6). Data are presented as mean and standard error of mean and analyzed by Student t-test (panels **B**, **D, G**, **H**, **I**, **J** and **K**), One-way ANOVA with Tukey’s multiple comparison post-hoc analysis (panel **L**) \**p*<0.05, **p<0.01, ***p<0.001

### Loss of myoglobin increases migratory capacity of hypoxic MCF7 cancer cells

Previous studies on MDA-MB-468 human breast cancer cells suggested that expression of MB might correlate with increased cell migration [17]. Thus, we investigated whether MB influences cell migration in MCF7 and SKBR3 cells by using the wound closure assay (Fig. 2A and B) and the invasion assay (Fig. 2C). Here again, the two cell backgrounds reacted differently upon the loss-of-function (LoF) of MB. While loss of MB in normoxic MCF7 cells did not alter cell migration, the loss of MB in normoxic SKBR3 cells decreased cell migration (Fig.2 B), aligning with previous studies [17]. However, at 0.2% O_2_, MB-KO MCF7 cells migrated almost 2 times faster than WT cells. SKBR3 MB-KO clones differed in cell migration (Fig.2B), thus yielding ambivalent results. The migration of MB-KO1 cells did not differ from WT SKBR3 but MB-KO2 cells migrated slower than WT SKBR3 cells. In addition, hypoxic MB-KO MCF7 invaded extracellular matrix-like substance significantly faster than WT cells (Fig. 2D) suggesting that expression of MB helps preventing cell migration and invasion in this background when O_2_ levels are low.

Since enhanced cell migration as seen in hypoxic MB-KO MCF7 cells is frequently associated with mesenchymal features of cells, we analyzed epithelial to mesenchymal transition (EMT) specifically in this background. EMT is a complex program that incorporates many pathways mediated by the transcription factors SNAIL, SNAI2 (or SLUG), TWIST and ZEB, whose differential expression in cancer has been shown to lead to EMT [43]. Fast migrating hypoxic MB-KO MCF7 showed indeed increased expression levels of EMT markers. The mean expression levels of 15hospho-Smad 2 and 15hospho-Smad 1/5/9 in hypoxic MB-KO cells were 2.3 and 2.5 times higher than in WT cells, respectively, while total Smad-2 and E-Cadherin protein levels did not differ (Fig. 2E and F). Interestingly, the mean expression levels of the p85 regulatory subunit of PI3K, previously shown to promote the motility activity of MCF7 cells [44], was 2 times higher in hypoxic MB-KO than in hypoxic control WT cells (Fig. 2E and F). Furthermore, hypoxic MB-KO MCF7 showed 3 times (*SLUG*), 2 times (*TWIST*) and 2.1 times (*ZEB1*) higher mean mRNA expression levels than hypoxic MB-WT cells (Fig. 2G). Mean mRNA levels of fibronectin 1 (*FN1*), which is associated with an invasive and metastatic breast cancer phenotype and EMT in MCF7 cells [45], were 2.2 times higher in MB-KO cells than in WT cells (Fig. 2G). Mean transcript levels of matrix metalloprotease 3 (*MMP-3* or stromelysin 1) in hypoxic MB-KO cells were 2.9 times higher than in hypoxic WT cells (Fig. 2G). In contrast, we observed no difference in MMP-9 or MMP-14 expression between hypoxic MB-KO and WT cells (not shown). Transcript levels of subunit B of PI3K, known to be dysregulated in MCF7 and to drive migration of cells [46], were 1.6 times higher in hypoxic MB-KO cells than in hypoxic WT controls (Fig. 2G). Our results indicate that many, but not all, EMT markers (e.g., E-Cadherin) are differentially up-regulated in hypoxic MB-KO MCF7 cells, implying that MB LoF in this cell background yields at least a partial EMT during hypoxia.

### Myoglobin interferes with estrogen receptor expression and ROS generation

MB adds to the prognostic value of ERα in human breast cancer patients [8]. Therefore, we analyzed if MB impacts the ERα expression in ER+ MCF7 cells (Fig. 3A), but refrained from doing so in ER-SKBR3 cells [47]. The mean expression levels of the nuclear long isoform of ERα in normoxic MCF7 MB-KO cells was 2 times higher than in normoxic WT cells (Fig. 3B), which was confirmed by immunofluorescence analyses that showed a higher number of ER positive cells (Fig. 3C and D). Re-expression of MB in the normoxic MB-KO cells resulted in downregulation of ERα expression, confirming our earlier observation (Fig. 1E and F). Because ERα protein can be destabilized by ROS [48, 49], we analyzed ROS levels (primarily superoxide and hydrogen peroxide) in normoxic (Fig.3G) and hypoxic (Fig.3H) MB-KO and WT MCF7 cells cultured for 72h. At normoxia, MB-WT cells showed 50% higher mean ROS levels than KO cells (Fig. 3G) in presence of positive (antimycin A: AmA) and negative (N-acetyl cysteine: NAC) assay controls. While levels of *SOD3* transcripts in MB-KO MCF7 cells were 10 times lower than in WT cells, other ROS detoxifying genes were not differentially regulated (Supplemental Fig. 4). However, this severe downregulation of *SOD3* by the loss of MB suggests the ERα destabilizing ROS in normoxic MB-WT cells to originate at sites outside of mitochondria. Unsurprisingly, 0.2% O_2_ hypoxia reduced mean ROS levels in WT cells by 2.4-fold relative to their normoxic levels, but not in MB-KO cells. MB-deficient MCF7 cells showed elevated ROS levels compared to WT controls under hypoxia (Fig. 3H), which might explain the normoxia-specific downregulation of ERα in MB-WT cells relative to MB-KO counterparts.

**Figure 4.**
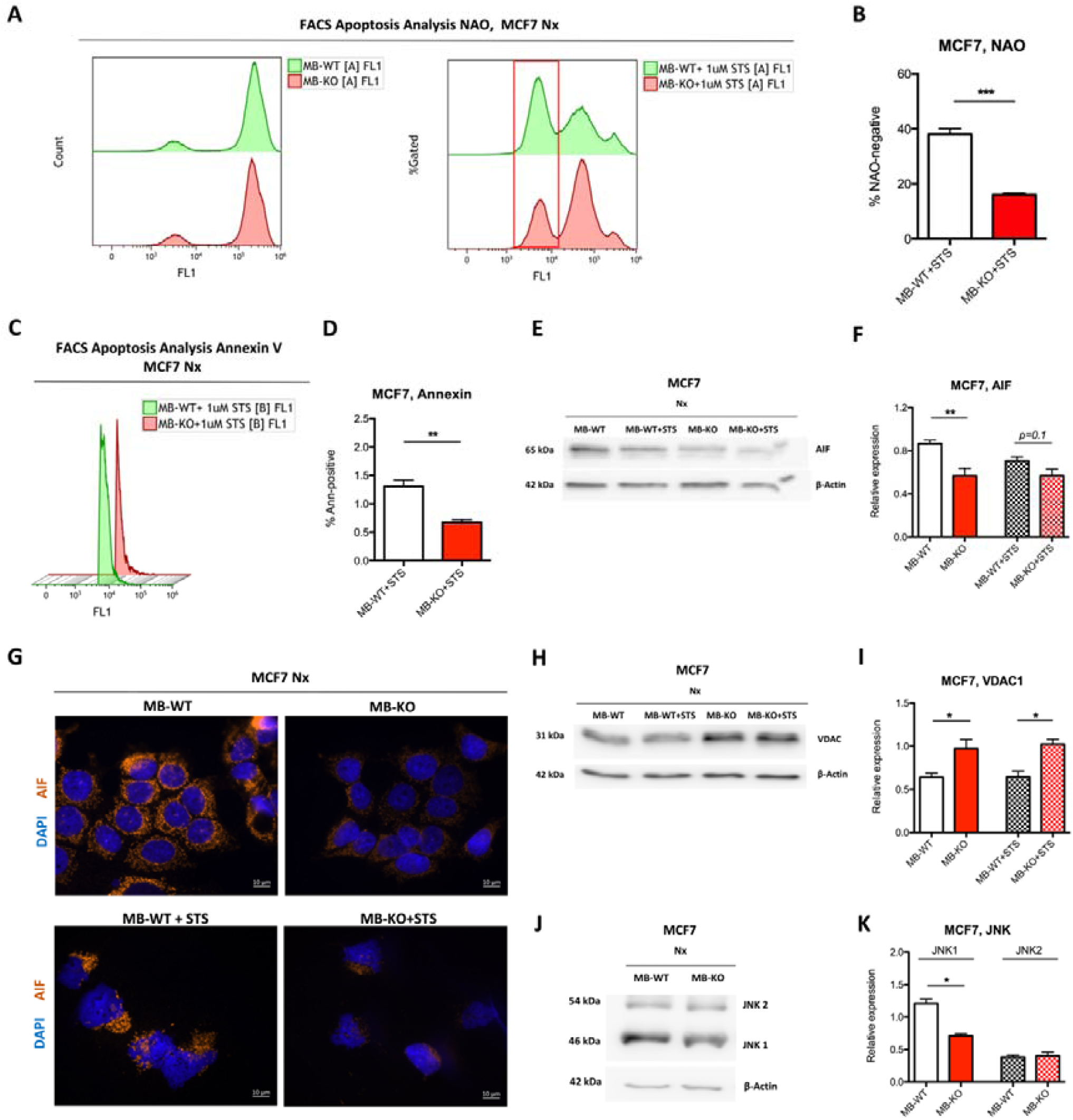
Myoglobin sensitizes breast cancer cells to apoptosis. (**A**) Representative FACS analysis of MB-WT and MB-KO clones of MCF7 cells stained by 10-N-Nonyl acridine orange (NAO), a fluorophore that forms a stable complex with reduced form of cardiolipin within the inner mitochondrial membrane of living non-apototic cells. 100’000 cells were incubated, in duplicates, at Nx for 72 h, followed by incubation without (left panel) or with 1 mM of the apoptosis inducing agent staurosporin (STS) for 6 h (right panel). Cell population with negative NAO fluorescence, marked with red box, correspond to the population of apoptotic cells. (**B**) Quantitative analysis of NAO-negative proportion of cells after with 1 mM of STS for 6 h (n=3). (**C**) Representative Annexin V staining based FACS analysis of MCF7 cells, incubated in duplicates at Nx for 72h, before treatment with 1 mM of the STS for 6 h, to detect cells at a later stage of apoptosis accompanied by live/dead stain of propidium iodide (PI). (**D**) Quantitative analysis of Annexin V-positive proportion of cells after with 1 mM of STS for 6 h (n=3). (**E**) and (**H**) Representative Western Blotting image (n=3) of whole tissue lysate of MB-WT and MB-KO clones of MCF7 cells cultured at normoxia Nx or Hx for 72h and treated or not with 1 mM of the apoptosis inducing agent staurosporin (STS) for 6 h, followed by staining for apoptosis inducing factor (AIF) (65 kDa) (**E**) or VDAC1 (31 kDa) (**H**) and β-actin (42 kDa) used as loading control. (**F**) and (**I**) Band intensity of AIF and VDAC1 proteins, respectively, after Western Blotting, from MB-WT (white) and MB-KO cells (red) of MCF7 cultured at Nx without (empty bars) and with treatment of 1 mM of STS for 6 h (dashed bars), was quantified using MCID Analysis 7.0 and normalized to β-actin (n=3). (**G**) Representative immunocytochemistry images of MB-WT and MB-KO MCF7 cells stained for AIF (orange) and DAPI (blue) after culturing at Nx for 72h (upper panels) or with treatment of 1 mM of STS for 6 h (lower panels). Scale bar is 10µm. (**J**) and (**K**) Representative Western Blotting image and band intensity analysis of whole tissue lysate of MB-WT (white) and MB-KO (red) clones of MCF7 cells cultured at normoxia Nx for 72h and stained for JNK1 (54 kDa), JNK2 (64 kDa) and β-actin (42 kDa) used as loading control. Data are shown as bar graph with mean and standard error of mean and analyzed by Student’s t-test *p<0.05; **p<0.01; ***p<0.001

### Myoglobin interferes with cancer cells’ response to chemotherapeutic but not ionizing irradiation treatment

We next tested whether the differential ER expression might influence response to anti-hormone therapy. Indeed, the clonogenic assay revealed a trend towards markedly fewer colonies of MB-KO cells treated with the selective ER modulator tamoxifen (TAM), as compared to WT cells under Nx but not at 0.2% O_2_ Hx (Fig. 3I). These results suggest that endogenously expressed MBO_2_ might contribute to regulating ERα expression in oxygenated breast cancer cells. In addition to antihormonal therapy, MB-containing particles were recently targeted towards cultured lung cancer cells and they improved intracellular O_2_ levels and thereby sensitized the cells to radiation therapy [50]. Contrary to the previous findings, our colony formation assays after 9 days showed no difference in the number of colonies when comparing normoxic or hypoxic MB-KO and MB-WT control cells in response to 3Gy irradiation (Fig. 3J). We also analyzed the response of MB-WT and MB-KO cells to doxorubicin (DOX) treatment, which is routinely used in treatment of several cancers including mammary carcinomas [51]. In normoxia, MB-KO MCF7 cells showed an elevated number of colonies compared to MB-WT cells in response to 5nM DOX (Fig. 3K). A similar, yet non-significant, trend was noted in response to 10 nM of DOX. Under hypoxic conditions, the colony formation of MCF7 WT and MB-KO cells did not differ at low or high concentrations of DOX (Fig. 3K), suggesting that only the presence of MBO_2_ sensitizes normoxic MCF7 cells to DOX treatment. Assessing the response of MB-proficient and -deficient SKBR3 cells to DOX was hampered by the complete lack of colony formation by this line. We, thus, needed to use the MTT assay to determine the MB-dependent sensitivity of these cells to DOX. Interestingly, MB-KO SKBR3 cells showed a higher MTT conversion rate than WT cells along increasing concentrations of Doxorubicin under both, normoxic and hypoxic conditions (Fig. 3L). Therefore, both oxy and deoxy MB can sensitize SKBR3 cells to DOX treatment. In conclusion, basal MBO_2_ abundance seems to interfere with hormonal receptor expression, perhaps via an intensified ROS generation, in oxygenated BrCa cells. Proficiency of MB in MCF7 and SKBR3 backgrounds sensitizes both cell types to treatment with DOX under high O_2_ concentrations, while it renders a TAM-based intervention of ERα-positive normoxic MCF7 cells far less effective. The status of endogenous MB in BrCa backgrounds allegedly has a strong influence on the outcome of different chemotherapeutic interventions. It does not, however, seem to impact the cells response to radiation therapy.

### Loss of myoglobin inhibits apoptosis in normoxic BrCa cells

Because loss of MB reduced ROS production in Nx MCF7 cancer cells and rendered these cells more resistant to DOX treatment (Fig. 3G and K), we next focused on the effect of MB-KO on apoptotic stimulation under Nx. Supplementation of human renal proximal tubule cells with ferrous myoglobin was previously shown to stimulate apoptosis [52]. In addition, presence of MB was demonstrated to stimulate apoptosis in prostate cancer cells [22]. We induced programmed cell death by incubation with 1µM staurosporin (STS) for 6h and quantified apoptotic cell populations by using flow cytometry with NAO staining (Fig. 4A) [35]. The mean number of apoptotic cells, denoted as completely negative for NAO-based fluorescence, of MB-KO MCF7 cells was 2 times lower than in MB-WT cells (Fig. 4B), suggesting that MBO_2_ indeed facilitates apoptosis. The NAO-intermediate pool of cells, located approximately one log below the positive population in Fig. 4A, right panel, were omitted in our quantification of apoptotic cells, as this pool presumably represents cells of mixed oxidized/reduced cardiolipin (CL) composition and/or of mixed location of CL to inner or outer mitochondrial membrane sites (see NAO-based FACS analysis in Materials and Methods section for more details). In a second apoptosis assay, we looked at flip of phosphatidylserine to the outer leaflet of the plasma membrane as an early indicator of programmed cell death. Here again, the mean number of Annexin-positive, apoptotic cells among the population of MB-KO MCF7 cells was 60% lower relative to WT cells after treatment with 1uM STS for 6h (Fig. 4C and D), further supporting that MB-deficiency greatly diminishes the frequency of apoptosis in oxygenated MCF7 cells. Because MCF7 cells do not to express caspase-3 [53, 54], which we verified (data not shown), we analyzed cellular levels of i) cleaved Lamin A, that is produced by cleaved caspase 6, as well as ii) cleaved caspase 7. No expression of cleaved Lamin A was detected in MCF7 cells by immunoblotting, nor could we detect a change in protein expression of cleaved caspase 7 in presence or absence of MB (data not shown). However, mean levels of apoptosis inducing factor (AIF) protein, which is a mitochondrial protein that induces caspase-independent apoptosis [55], was reduced by 33% in MB-KO than MB-WT cells (Fig. 4E and F). After treatment with STS for 6h, less nuclear fraction of the AIF protein was associated with the loss of MB as compared to MB-WT cells in conjunction with the weaker overall AIF expression in MB-negative cells (Fig. 4G). Additionally, MB-KO MCF7 cells up-regulated protein-level expression of the mitochondrial voltage dependent anion (VDAC1), a protein involved in metabolic apoptosis pathways [56], by 1.6 and 1.5 times relative to MB-WT cells in presence or absence of 1uM STS for 6h, respectively (Fig. 4H and I). Furthermore, the mean expression levels of JNK1, but not JNK2, in normoxic MB-KO cells were 2 times lower than in MB-WT cells (Fig. 5J and K) suggesting that lower ROS levels in normoxic MB-KO cells may not be sufficient to activate JNK1-dependent apoptosis [57]. In summary, our data show that MBO_2_ promotes apoptosis in oxygenated MCF7 cancer cells. In contrast to apoptosis, MB-KO MCF7 cells expressed 2.6-fold higher levels of the autophagy-related gene protein 5 (ATG5) as compared to MB-WT cells at normoxia but not at 0.2% O_2_ hypoxia for 72h (Supplemental Fig. 5A and B). The mean levels of the ATG5 and ATG12 complex, together recruiting microtubule-associated protein 1 light chain 3 (LC3), did, however, not differ under normoxia or hypoxia. Yet, the lipidated active form of LC3 was 3-fold higher in MB-KO MCF7 than in WT cells at normoxia but not at hypoxia (Supplemental Fig. 5A and B), while the cytoplasmic inactive form showed a similar increasing trend under normoxia (Supplemental Fig. 5 A and B). While the loss of MBO_2_ in normoxic MCF7 cancer cells prevents apoptosis, at the same time, it may activate the autophagy machinery. However, this notion must be further verified by incubating with autophagy inhibitors and inducers. Together, these changes might enhance cancer cell survival, if oxygen is sufficiently available.

**Figure 5.**
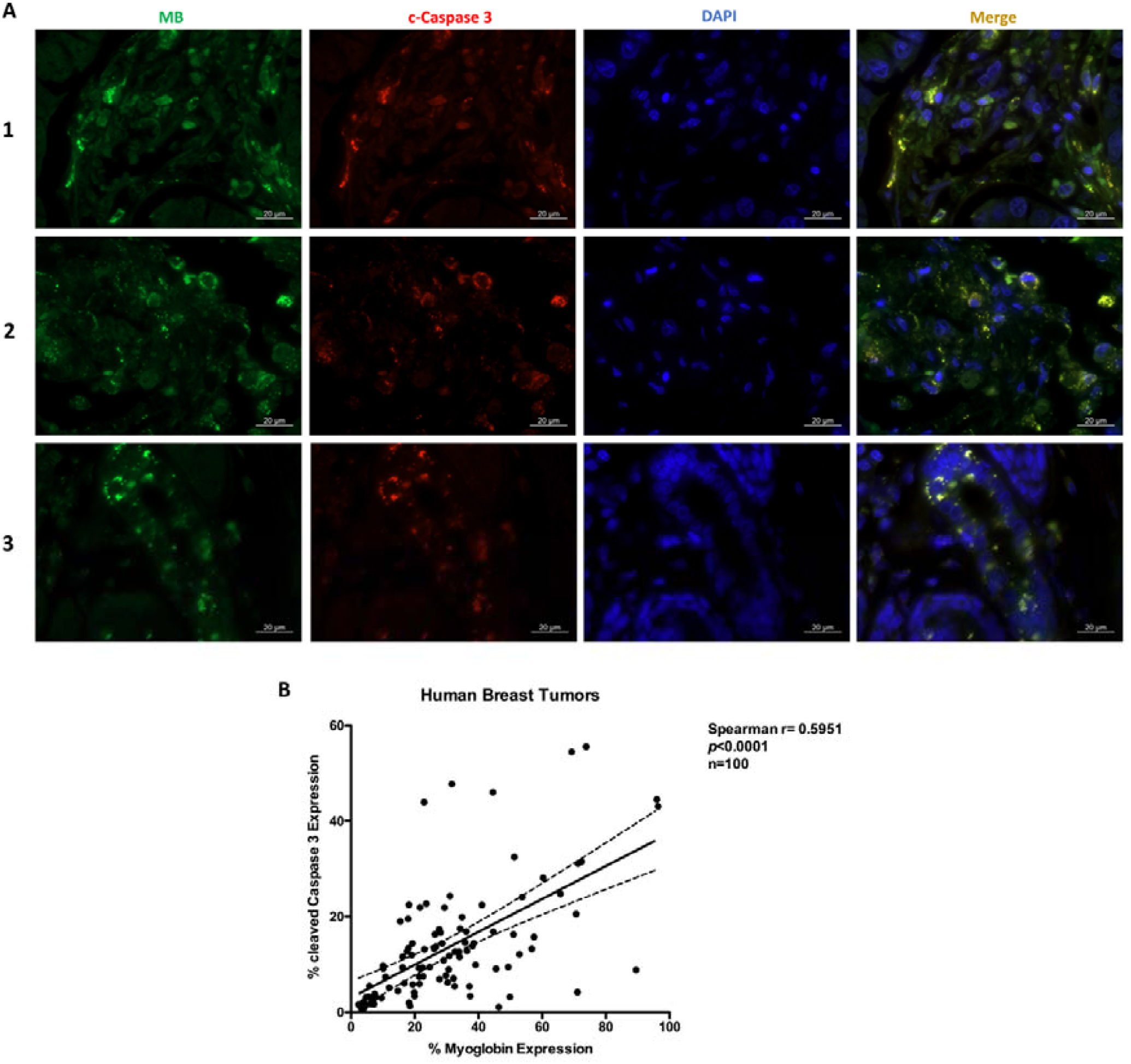
Expression of myoglobin in human invasive ductal carcinoma tumors correlates with increased cleaved caspase 3 expression *in vivo*. (**A**) Representative immunofluorescence image analysis of the expression of myoglobin (green), cleaved caspase 3 (red) and DAPI (blue) from breast tissue microarray of human invasive ductal carcinoma tumors that are estrogen and progesterone receptors positive. Scale bar is 20 µm. (n=100) (**B**) Correlation analysis of % of aforementioned stains signal areas normalized to tissue areas (n=100). Data are shown as regression analysis with spearman r correlation test *** p<0.0001

To determine whether our in vitro observations regarding apoptosis are also reflected in human cancer tissues *in vivo*, we measured MB and cleaved caspase 3 expression in tissue arrays from human invasive ductal carcinoma (Fig. 5A). The expression of MB in these tumor tissue cohort correlated positively with expression of cleaved caspase 3 (Fig. 5B). Taken together, these data are consistent with our observation that presence of MBO_2_ in breast cancer cells and tissues promotes apoptosis.

### Differential gene expression analysis by RNA-Seq

Genes differentially expressed between MB-KO and MB-WT MCF7 cells were analyzed by RNA-Seq after incubation at Nx and Hx (0.2% O_2_) for 72h (n=3). 281 genes were upregulated, and 128 genes were downregulated in response to MB-knockout in normoxic MCF7 cells (p adjust ≤0.05). Under Hx, 180 genes were up- and 81 genes down-regulated in MB-KO cells vs WT controls (p adjust ≤0.05). The overrepresentation analysis using g:Profiler revealed 12 GO-terms of biological processes for the normoxic comparison, while for the hypoxia data, a 100 GO-terms were enriched. Figure 6 shows the GO-terms clustered in Cytoscape. In the Nx dataset, overrepresented GO-terms are related to cell adhesion (‘cell adhesion’, ‘biological adhesion’ and ‘cell-cell adhesion via plasma-membrane adhesion molecules’) and cell migration (‘cell migration’). Additional GO-terms are related to developmental processes such as ‘digestive system development’, ‘animal organ development’ and ‘tube development’. GO-terms for the Hx dataset are related to signaling (e.g. ‘signal transduction’, ‘cell surface receptor signaling pathway’, ‘cell communication’) and to response to different stimuli (e.g. ‘response to external stimulus’, ‘response to organic substance’, ‘response to chemical’). Additional categories are related to cell motility, taxis and cell adhesion. Furthermore, genes related to ‘cell death’ and ‘programmed cell death’ were also overrepresented by the loss of MB. The terms ‘reactive oxygen species metabolic process’ and ‘regulation of fat cell differentiation’ appeared as well. Additional terms are related to general developmental processes (e.g. ‘anatomical structure development’, ‘system development’, ‘cell differentiation’) and to the development of the circulatory system (e.g. ‘tube development’,’ blood vessel development’,’ circulatory system development’). Full lists of all significantly differentiated genes at Nx and Hx are shown in (Supplemental Tables 1 and 2), including the already mentioned SLUG and SMAD genes (see Fig. 2).

**Figure 6.**
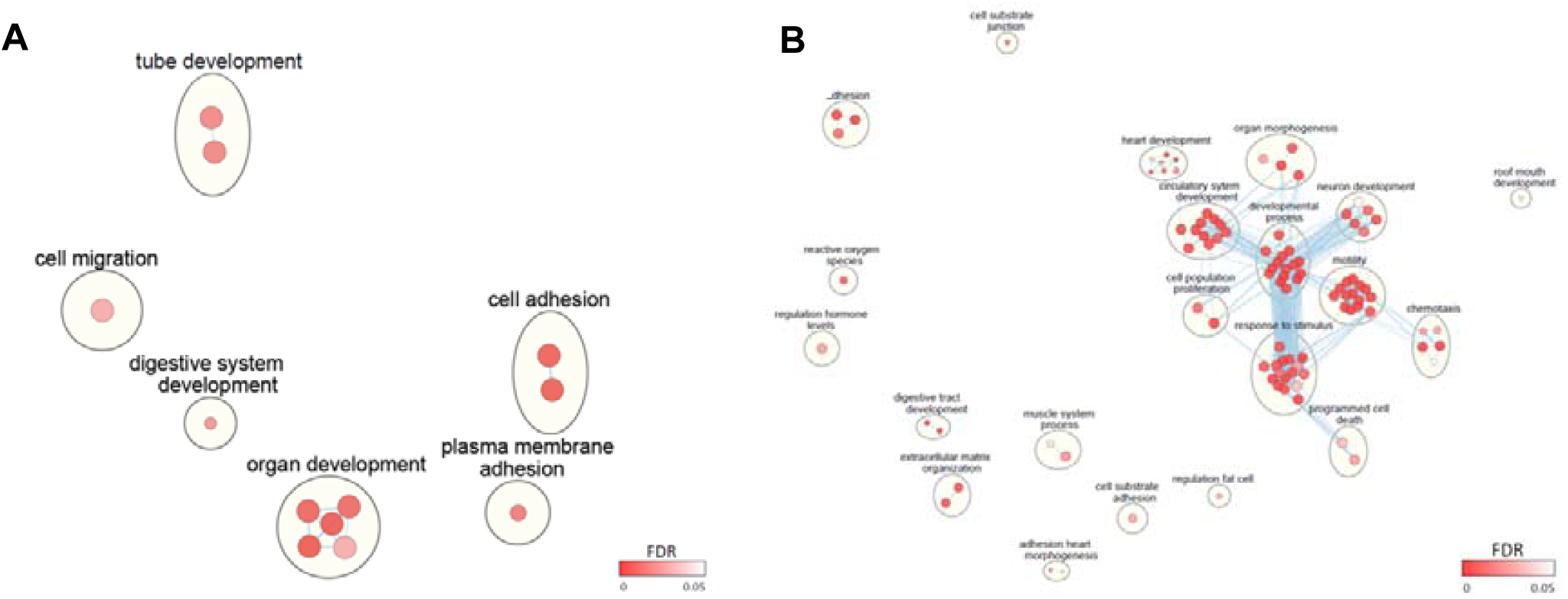
Enrichment maps for (A) normoxia and (B) hypoxia datasets. Each node represents a single GO-term. Blue lines are drawn between terms with a Jaccard-Index < 0.25. The FDR is represented by the node color. Annotations were created using AutoAnnotate and adjusted by hand. Clusters were arranged using CoSE algorithm, scaled and adjusted by hand.

### Breast cancers with mutant p53 have less MB and worse prognosis

To determine a possible correlation between MB and p53 expression, we determined p53 protein expression in tissue arrays from human invasive ductal carcinoma (n=288) that we have analyzed for MB expression in our previous work [8]. The expression of p53 in this tumor tissue cohort was heterogenous, varying between absent, weak, moderate and strong expression (Fig. 7A). 70.5% of tumors displayed WT p53 expression while 29.5% exhibited p53 mutation with either complete loss of expression or pathological accumulation of the protein. Patients with p53-WT tumors or MB positivity showed better prognosis and overall cumulative survival as compared to tumors with mutant p53 status or MB negative expression, respectively (Fig. 7B and C). MB expression was significantly associated with p53 status; 32.9% of p53-mutant tumors were devoid of MB while only 19.2% of p53-WT showed MB negativity (Table 1). 36.5% of p53 mutant tumors showed moderate to strong MB expression relative to 48.3% of p53-WT tumors. Taken together, p53 WT tumors accumulated more MB protein as compared to p53 mutant tumors. Whether prognostic value of MB is dependent on p53 status is currently unknown. Nevertheless, MB is associated with better prognosis in patients group stratified by p53, with visually more pronounced value in p53-mutant cases (Fig. 7D and E).

**Figure 7.**
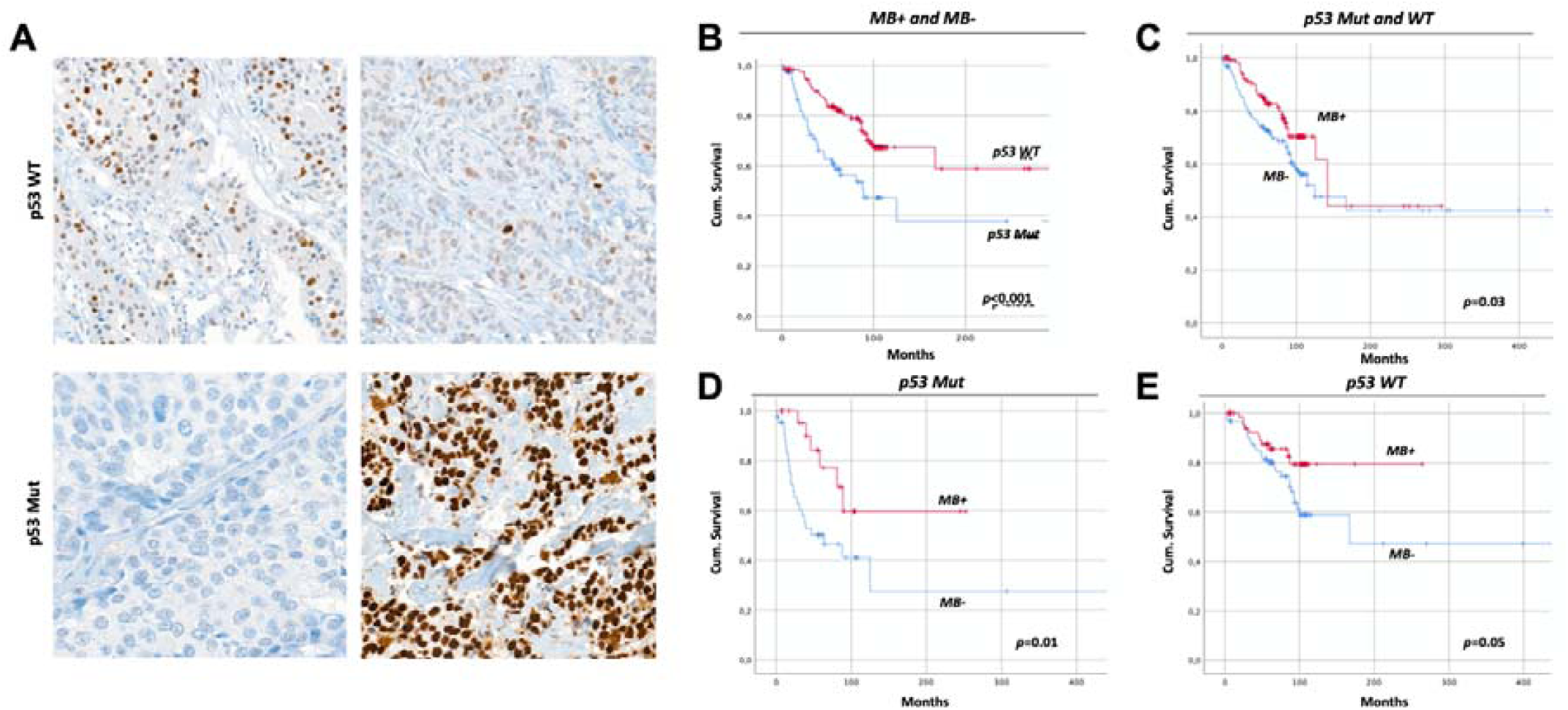
Myoglobin expression is linked to better prognosis independently of p53 status. (**A**) Representative image of immunohistochemistry staining of p53 in cohort of human invasive ductal carcinoma (n=288). Upper left and upper right panels show moderate and weak expression of wildtype (WT) p53, respectively. Lower left and lower right panels show complete expression loss and pathological accumulation of p53, respectively, displaying the two forms of p53 mutation (p53 Mut) as detected by immunohistochemistry. Scale bar is 200 mM. (**B**-**E**) Kaplan-Meier analyses of a cohort of 288 primary breast cancer cases displaying (**B**) tumors with p53 WT expression and their significantly improved cumulative overall survival prognosis compared with tumor cases with p53 Mut. (**C**) tumors with high MB expression (MB+) show a significantly improved cumulative overall survival prognosis compared with cases of low Mb expression (MB−) in mixed p53 background status. (**D**) and (**E**) MB+ tumors show significantly improved cumulative overall survival prognosis compared with MB− tumors, in p53 Mut as well as p53 WT cohorts, respectively.

**Table 1.** Correlation of myoglobin expression to p53 status.

| p53 Status | MB Expression n (%) |  |  |  |  | *P* |
| --- | --- | --- | --- | --- | --- | --- |
|  | No | Mild | Moderate | Strong | Total |  |
| Mutant | 28 (32.9) | 26 (30.6) | 23 (27.1) | 8 (9.4) | 85 (100) | 0.017 |
| Wildtype | 39 (19.2) | 66 (32.5) | 70 (34.5) | 28 (13.8) | 203 (100) |  |
| Total | 67 (23.3) | 92 (31.9) | 93 (32.3) | 36 (12.5) | 288 (100) |  |

## Discussion

The most widely recognized role of myoglobin (MB) is the storage and facilitated delivery of O_2_ to mitochondria in myocytes under hypoxia. However, the very low concentrations at which MB is estimated to be endogenously expressed in cancer cells seems to preclude this function [8]. Consistent with this, we also previously confirmed, through respirometry studies, that endogenous (ie. not overexpressed) MB does not play a significant role in (re)oxygenating breast cancer cells and may have other tumor suppressor functions. Here, genetically modified MCF7 breast cancer cells lacking MBO_2_ displayed enhanced survival with upregulated levels of G1/S cell cycle markers. When compared with MBKO, SKBR3 breast cancer cells without MB exhibited enhanced survival. When looking deeper into molecular mechanisms in the MCF7 background, we noticed that the loss of MBO_2_ in these cells was associated with fewer ROS being generated in normoxic cells. This resulted in a) upregulated protein levels of ERα, b) a sensitization of cells to tamoxifen-mediated inhibition of colony formation, and c) in desensitizing cells towards chemotherapy-induced apoptosis (e.g., via DOX treatment). MBO_2_ devoid cells upregulated autophagy proteins, possibly mediating the improved cell survival, yet to be further validated by experiments that involve autophagy inhibitors and inducers. Hypoxic MCF7 cells lacking MB showed upregulated EMT markers and increased motility along with increased invasive behavior, hence a heightened metastatic potential. The lack of MB combined with O_2_ limitation activated SMAD signaling but had no effect on the cellular response to irradiation or chemotherapy. The expression of MB in human tumor tissue cohort correlated positively with expression of cleaved caspase 3. We report on a novel set of roles and molecular mechanistic insights mediated by basal MB expression in breast cancer cells. The data presented in this study suggest MB to contribute to the tumor suppression in a subset of breast cancer cells (i.e. p53-WT, ERα-positive) and tumors by reducing estrogen hormone receptor expression, cellular migration as well as cell survival mechanisms. Moreover, our data suggest non-muscle MB to aid in predicting the metastatic behavior of breast cancer patients and therefore predict patient outcome. In addition, MB might serve as a therapeutic target with careful consideration of other factors as p53 and hormone receptor status of the tumor. Importantly, our studies demonstrate that those roles of MB are operant even with low basal expression levels.

### Myoglobin attenuates normoxic breast cancer cell survival and ERα signaling while enhances apoptosis and response to chemotherapy by increasing ROS levels

Loss of MB expression correlates with poor patient survival [8, 9] suggesting that MB is a tumor suppressor. While supraphysiological levels of MB after ectopic overexpression downregulate cyclin E and reduce the cell cycle progression of cancer cells [58], we show here that endogenous levels of MB in human MCF7 breast cancer cells antagonize cell survival rather than slowing proliferation. We show that the deficiency of the globin increased levels of both cyclin D1, which is a marker of poor prognosis in breast cancer patients [59], and of cyclin E, without increasing cellular proliferation. In contrast, loss of MB in LNCaP prostate cancer cells resulted in low p21 transcripts and increased proliferation [22] suggesting that MB may exert tumor-type specific functions. In addition, our previous results in MDA-MB 468 breast cancer cells using si/shRNA *MB* knockdown revealed decreased proliferation with MB downregulation. The differences may be explained by different cell line biology or different knockout mechanism [60]. The induced cyclin D1 levels in our MB-KO cells may also occupy a cell cycle independent role, e.g., by directly binding to the hormone-binding domain of ERα, mediating gene transcription also in the absence of estrogen [60]. Indeed, we observed induced ERα expression levels and increased cell survival in normoxic MB-KO cells, which might be either explained by the higher cyclin D1 levels and/or by the lower levels of ROS. Elevation of ROS levels can downregulate ERα [48, 49, 61]. ERα was reported to protect cancer cells from p53-mediated apoptosis induced by DNA damage [62] and from ROS-mediated apoptosis [63]. Interestingly, the Hx GO-term analysis shows terms related to signaling (e.g. ‘Regulation of signaling’, ‘signaling’ and ‘cell surface receptor signaling pathway’). Additionally, genes related to ‘regulation of hormone levels’ were overrepresented.

In addition to increasing ERα expression, the loss of MB in normoxic MCF7 cells prevented apoptosis and increased autophagy proteins and thus, increased cancer cell survival. We show that AIF levels and JNK activation were reduced in MB-KO cells. Presence of MB in the normoxic cells was associated with elevated ROS levels, along with a very strong compensatory overexpression of the *SOD3* antioxidant gene. This pattern might confine elevated ROS to sites other than mitochondria (i.e. cytosol), which could promote JNK signalling and downstream apoptosis in oxygenated cells [64]. The fact that presence of MB even promotes abundance of *SOD3* in hypoxic MCF7 needs to be further explored. We also show positive correlation of MB and apoptosis protein expression in human invasive ductal carcinoma tissue as proof of concept that our mechanistic findings may hold true in clinical datasets where myoglobin expression is known to correlate with improved outcome. Indeed, the terms ‘cell death’ and ‘programmed cell death’ were detected in the hypoxia dataset of our transcriptomic analysis. Additionally, we observed increased expression levels of VDAC1 in MB-KO cells. VDAC1, despite its role in mitochondrial metabolism [65], acts as a suppressor of apoptosis by downregulating caspases, p53, cytochrome c and growth in cancer cells [66] and yeast [67]. Recently, VDAC1 expression was correlated with poor prognosis in human breast cancer patients [68]. The loss of MB in normoxic MCF7 cells, however, not only reduces apoptosis, but may, in parallel, increase autophagy to eventually increase longevity of mammary carcinoma, and perhaps many cancers cell types [69-71], yet to be further studied and verified.

Upon closer inspection of MB’s impact on autophagy, we observed ATG5 and both cytoplasmic and lipidated (active) forms of LC3 to be upregulated in MB-KO cells. Loss of MB function in heart and brown adipose tissues was accompanied by lipid accumulation and shift in lipid droplets equilibrium from numerous and small droplets into fewer but bigger droplets, respectively [18]. Interestingly, MB overexpression in breast cancer cells was shown to promote degradation of the key autophagy regulator E3 ubiquitin ligase, parkin [58], which is required to mark dysfunctional mitochondria and organelles for autophagy. The increased survival of MB-KO MCF7 cells suggests that its expression might be a good predictor of treatment success in BrCa cells. Indeed, we show that the loss of MB lowers the efficacy of doxorubicin under normoxia. Doxorubicin targets cancer cells by interfering with DNA replication and, secondarily, by generating ROS [51, 72], which are reduced in MB-KO cells. However, the role of MB on cancer therapies *in vivo*, needs to be further studied. In contrast to doxorubicin, the lack of MB did not influence the radiation efficacy of MCF7 cells, although the targeted MB delivery to lung cancer cells enhanced O_2_ delivery and increased radiation-induced apoptosis [50]. In conclusion, our data suggest that presence of MBO_2_ in oxygenated breast cancer cells renders cancer cells less viable, mainly through its pro-apoptotic functions, and more susceptible to anti-cancer treatments.

### Myoglobin regulates migratory capacity of hypoxic cancer cells

MB-overexpressing lung cancer cells in an ectopically induced mouse model approached similar levels of MB protein as commonly observed in skeletal muscle and hence improved tumor oxygenation, enhanced tumor differentiation, reduced tumor growth as well as local invasion and spontaneous metastasis [73]. However, it is still a matter of debate whether the actual low-level expression of MB detected endogenously in tumors confers meaningful O_2_ storage or buffering capacities [8]. We performed our experiment at normoxic and hypoxic conditions to mimic the oxygenation heterogeneity of clinical tumors and to compare the cellular function of oxygenated (MBO_2_) vs. mainly deoxygenated MB in breast cancer (i.e. applying the Hill equation). Tumor hypoxia correlates with enhanced cancer cell migration. Indeed, hypoxic cells lacking MB showed increased migration and invasion capacity by acquiring some mesenchymal features of the EMT. We demonstrated that the loss of MB increased the migration of hypoxic, but not normoxic, MCF7 breast cancer cells through the activation of the SMAD signaling pathway and upregulation of MMP3 expression. Hypoxic MB-KO cells exhibited upregulated phospho-Smad 2 and phospho -Smad 1/5/9 proteins expression as well as upregulation of their downstream targets *TWIST*, *SNAIL*, *SLUG* and *ZEB1*. This notion is strongly supported by GO-terms related to motility under hypoxia in RNA-Seq analysis presented in this study. Additionally, multiple terms associated with adhesion were detected among genes overrepresented in the normoxic and hypoxic data set. Although no increase in migration under normoxia was observed, one GO-term related to migration and three related to adhesion were part of the normoxia results. Enrichment analyses in our recent transcriptomic analysis of three different epithelial cancer cells with basal or reduced MB expression revealed that the GO categories ‘cell motion’ and ‘cell migration’ were significantly overrepresented among the upregulated genes of MB knocked down LNCaP cells compared to WT cells [22]. Hypoxic MB-KO cells showed upregulation of *FN1*, a glycoprotein ECM component of the mesenchymal compartment that is not normally expressed by breast tissue, thus indicating the switching to an increased capacity of invasive and metastatic cancer cell behavior and induced EMT in MCF7 cells [45]. Complementary to that, *MMP3* was differentially upregulated in MB-KO cells, indicating active ECM degradation which is an important step during tumor metastasis. In contrast, other markers, such as E-Cadherin, did not change their expression level in correlation with the MB status, suggesting a partial rather than complete switch to an EMT phenotype. This finding might be clinically relevant, as cells, which have undergone a partial EMT, were shown to be highly enriched in lung metastases, while cells that have undergone a full EMT retain a more quiescent mesenchymal phenotype and do not colonize the lung, suggesting the *partial* EMT to contribute more to breast cancer metastasis [74].

### Myoglobin in MCF7 vs. SKBR3 and the hypothesized interplay with p53

In contrast to MFC7 cells, the loss of MB in SKBR3 breast cancer cells neither influences migration under hypoxia nor the cells survival under normoxia (see data summary in Table 2), suggesting that our findings may not only be specific for different tumor types but also for different subtypes of a cancer. MCF7 and SKBR3 cells differ in their ER, PR, HER2 status, with MCF7 representing the luminal subtype while SKBR3 mirror the HER2-overexpressing subtype [75]. Moreover, both cell lines diverge in their p53 status. While MCF7 cells express wt p53, SKBR3 cells express a mutated gain of function (GoF) p53^Arg175His^ variant [76]. p53 dysregulation triggers the development of around 52% of human mammary carcinoma [77-80]. Around one third of cases are associated with an inactivation of tumor suppressor p53 by spontaneous (somatic) mutation [76, 77, 81], making the loss of p53 the most common genetic defect related to a single gene in breast cancer [78, 79]. Despite the inability to directly drive p53 target gene expression, mutated p53 could function by interacting with other proteins. We previously showed multiple tumor suppressor p63 [82] target genes to be downregulated in MB positive MDA-MB 468 breast cancer cells due to interaction and inactivation of the mutated p53^Arg273His^ leading to enhanced cell survival and decreased apoptosis [22]. In addition, mutated p53 may act as a cofactor of anti-apoptotic genes [83]. MB-mediated activation and stabilization of the p53 mutation via increased NO·, ROS and HIF1α levels lead to a tumor-promoting effect of MB specifically for GoF p53 expressing cells (MDA-MB 468 and SKBR3) [84]. This might explain the increased survival and migratory capacity of MB-WT SKBR3 cells. The assumption that the p53 status could predict on how MB functions in cancer cells is supported by observations in the GoF p53^Arg273His^ -containing MDA-MB 468 breast cancer cells that showed less migration when MB was knocked down under hypoxia [17, 22]. However, MB expression in our human patient’s cohort favoured prognosis and overall patients’ survival independently of p53 status. The hormonal receptor status for these tumors were, however, not studied. In particular, the p53 status might be a better predictor of the role of MB in migration than the tumor type itself, because prostate p53 wt LNCaP show enhanced migratory phenotype when MB is knocked down [22]. Regarding the cells we used, this study shows that the loss of MB increases cell survival in p53 wt MCF7 cells and downregulates apoptosis in MCF7, as it previously diminished the expression of apoptosis-promoting genes in wt p53 LNCaP, Conversely, the impact of MB on cell survival in p53 GoF MDA-MB 468 was inconclusive [22], suggesting MB to play a role in p53-dependent apoptosis. On the other hand, the response to doxorubicin may not depend on the p53 status, since MB LoF in both SKBR3 and MCF7 cells increased the resistance to this chemotherapeutic intervention. Together, MB in breast cancer cells emerges to function in p53-dependent and -independent ways.

**Table 2.** Effect of MB expression in MCF7 and SKBR3 cells as compared to MB-KO cells.

| Effect (O<sub>2</sub>) | MCF7 | SKBR3 |
| --- | --- | --- |
| Growth curve (Nx) | ↓ | ↑ |
| Migration (Hx) | ↓ | inconclusive |
| Response to Doxorubicin (Nx) | ↑ | ↑ |
| Hormonal receptors [75] | ER+ PR+ | ER- PR- |
| p53 status [76] | WT | GoF Mutation |
| Tumor subtype [75] | Luminal A | HER2 overexpressed |

## Conclusion

In conclusion, our study established endogenously expressed MB to contribute to tumor suppression mechanisms in breast cancer cells and some developing tumors in ways depending on the cell’s oxygenation status. Normoxic breast cancer cells lacking MBO_2_ show an enhanced survival and more resistance to chemotherapy induced apoptosis, while MB-deficient hypoxic cancer cells display augmented migratory and invasive phenotypes. Our findings suggest MB to contribute to the tumor suppression in subset of breast cancer cells by regulating cell cycle markers, ROS generation and hormone receptor expression, controlling cellular motility as well as cancer cell survival and death mechanisms. Moreover, our data suggest non-muscle MB to aid in predicting the metastatic behavior and chemotherapeutic response of breast cancer patients and therefore predicting patient prognosis.

### Statistics

Statistics were performed using GraphPad Prism6 (GraphPad Software, San Diego, US). Normal distribution of data populations was analyzed with the Kolmogorov-Smirnov test. For comparing two groups, Student t-test was employed for parametrically distributed and a Mann-Whitney test for non-parametrically distributed data. For comparing between more than two groups of normally distributed data, one-way ANOVA with Tukey’s multiple comparison post-hoc analysis was used. SPSS was used to calculate the KM curves of p53mut vs. p53wt cases.

## Acknowledgement

The laboratory work was partly performed using the logistics of the Center for Clinical Studies at the Vetsuisse Faculty of the University of Zurich.

## Author contributions

M.A.A. performed all work related to MCF7 cells as well as human tumors and J.A. performed work related to SKBR3. M.A.A and J.A. generated knockout clones. M.A.A. designed experiments, analyzed and interpreted the data and wrote the manuscript. M.T. technically supervised generation of knockout cells and revised manuscript. F.G. provided expertise with human tumors staining and analysis. A.B. generated MB-mCherryplasmids. A.B., R.P., T.H. performed RNA-Seq analysis and RNA-Seq based genotyping of MCF7 clones. P.S. performed p53 staining. G.K. evaluated p53 staining. G.K., A.B., T.H. and M.G. provided intellectual input and revised the manuscript. T.A.G. conceptualized and supervised the project and revised the manuscript.

---

**Supplemental Fig. 1.**
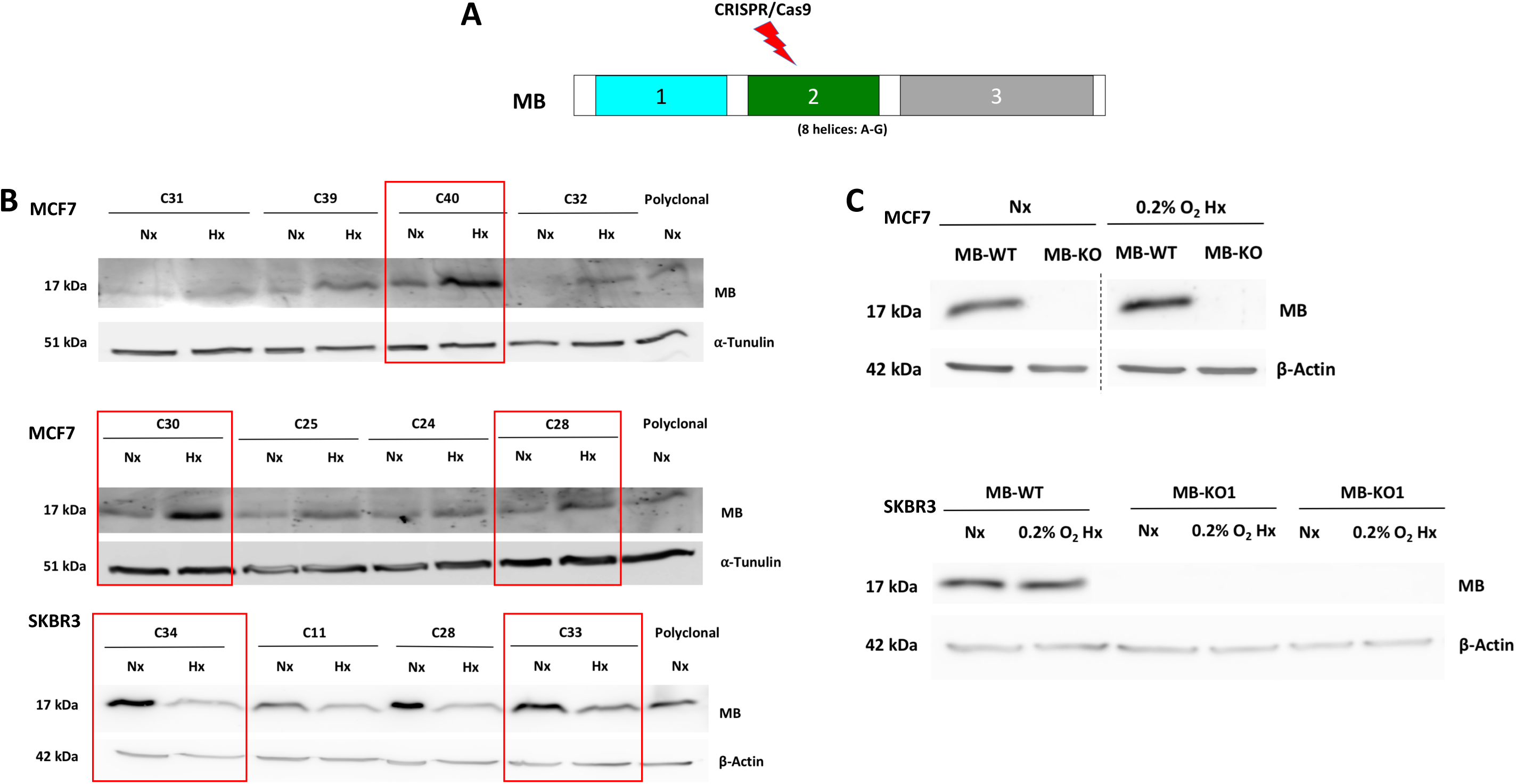

**Supplemental Fig. 2.**
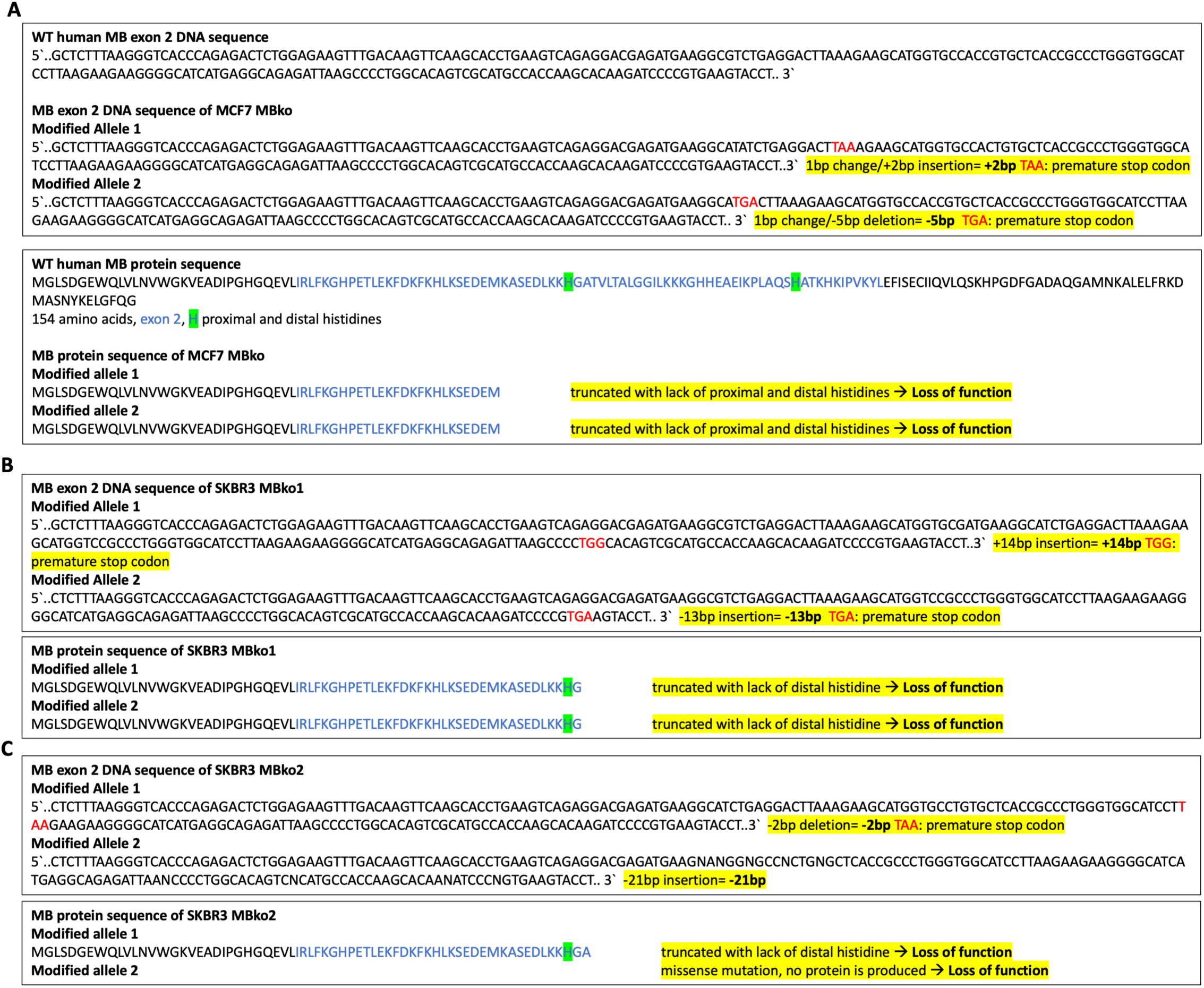

**Supplemental Fig. 3.**
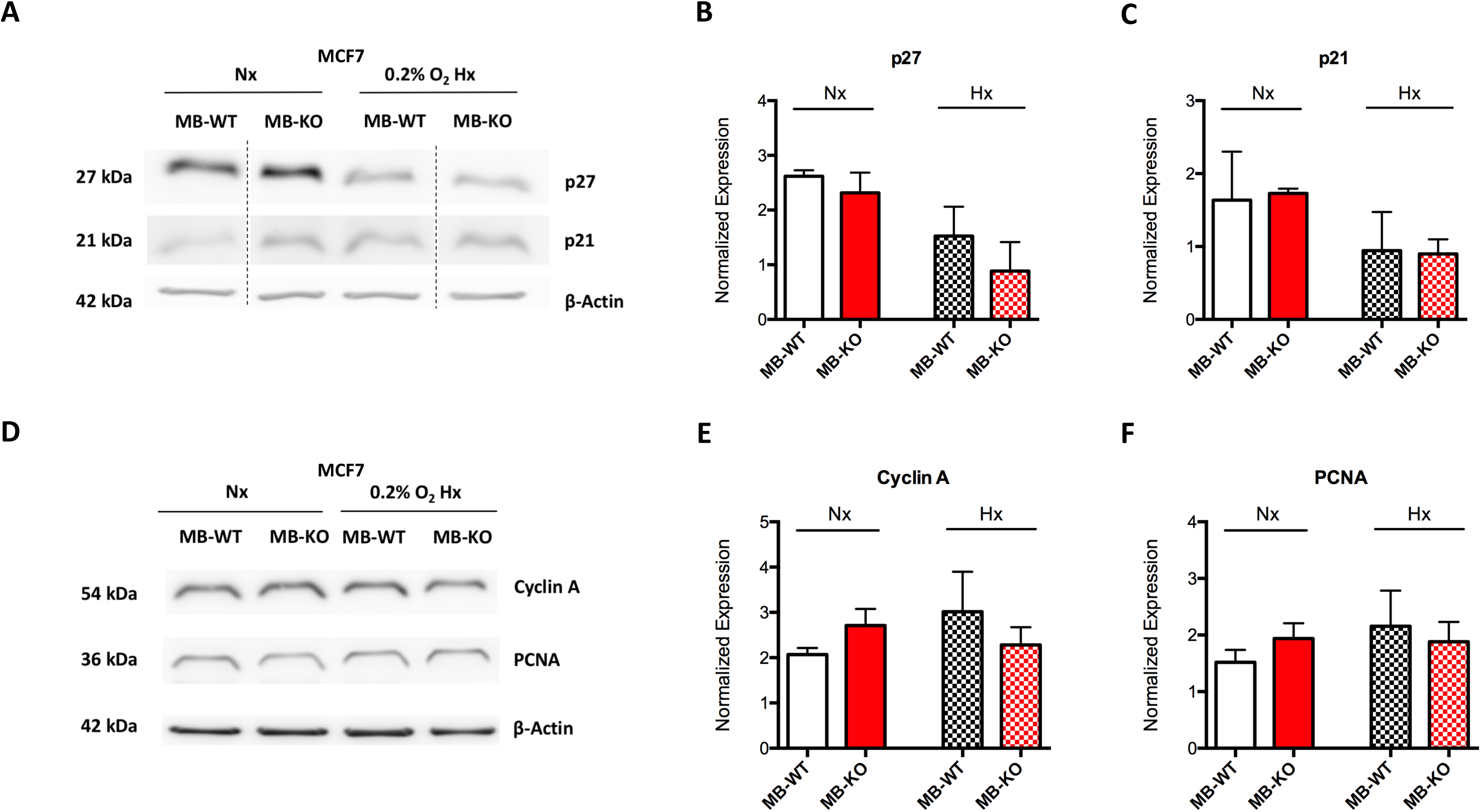

**Supplemental Fig. 4.**
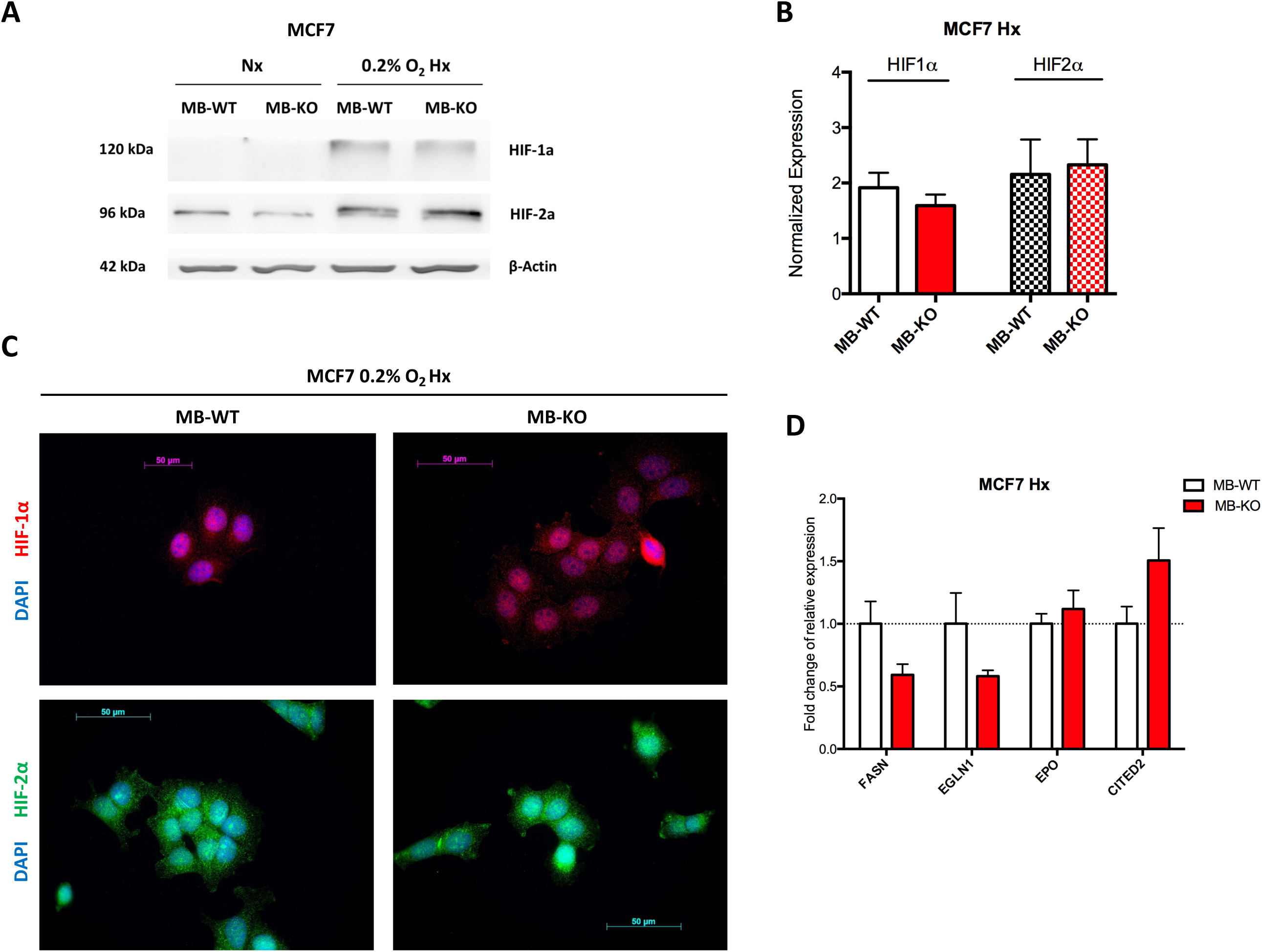

**Supplemental Fig. 5.**
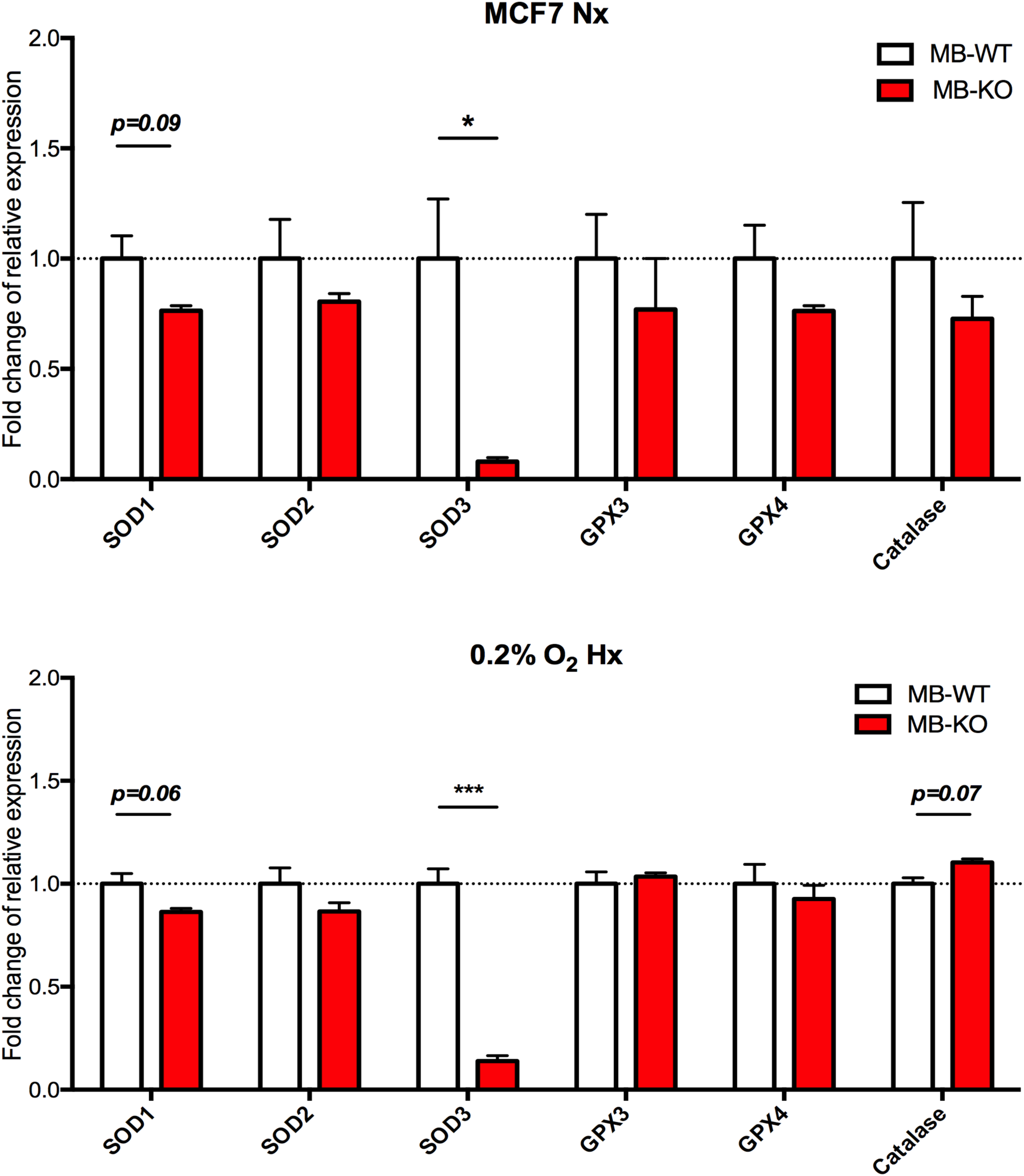

**Supplemental Fig. 6.**
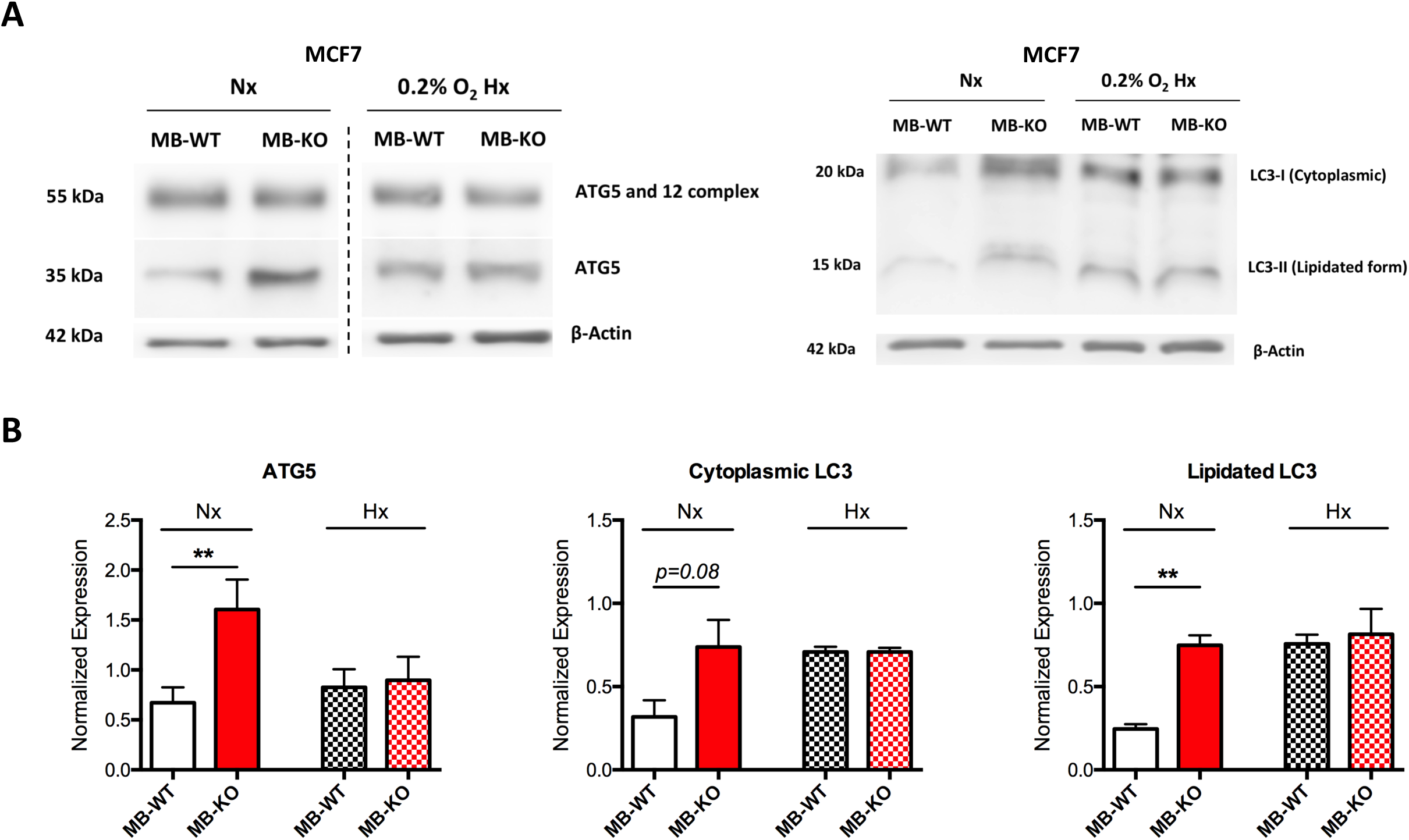

