## Supplemental Material for "Pro-apoptotic and anti-invasive properties underscore the tumor suppressing impact of myoglobin on subset of human breast cancer cells"

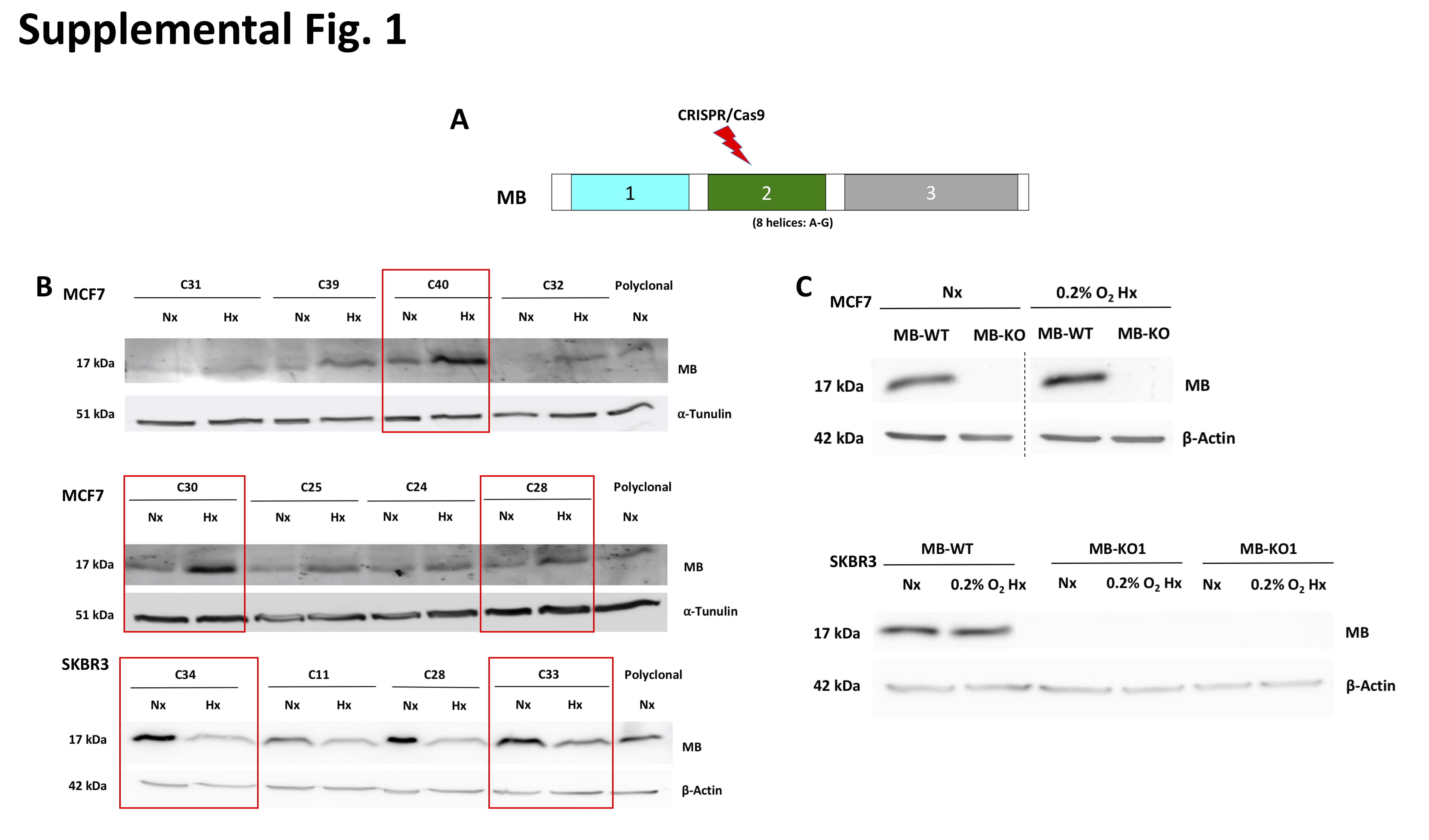
**Supplementary Figures**

**Supplemental Figure 1. Generation of MB knockout clones from MCF7 and SKBR3 cells**

**(A)** Representation of human *MB* gene composed of 3 exons. Exon 2 was targeted to knockout the gene by CRISPR/Cas9 to obtain loss of function. **(B)** Representative Western blot to examine basal MB expression at normoxia (Nx) and (Hx, 0.2% O_2_) in different monoclonal cell lines (denoted as C: clone and number) produced from the paternal MCF7 or SKBR3. **(C)** Representative Western blot stained for MB (17 kDa) to verify the MB knockout (MB-KO) in MCF7 and SKBR3 cells Cells were exposed to normoxia (Nx) or hypoxia (Hx, 0.2% O_2_) for 72 before protein extraction. (n=4).

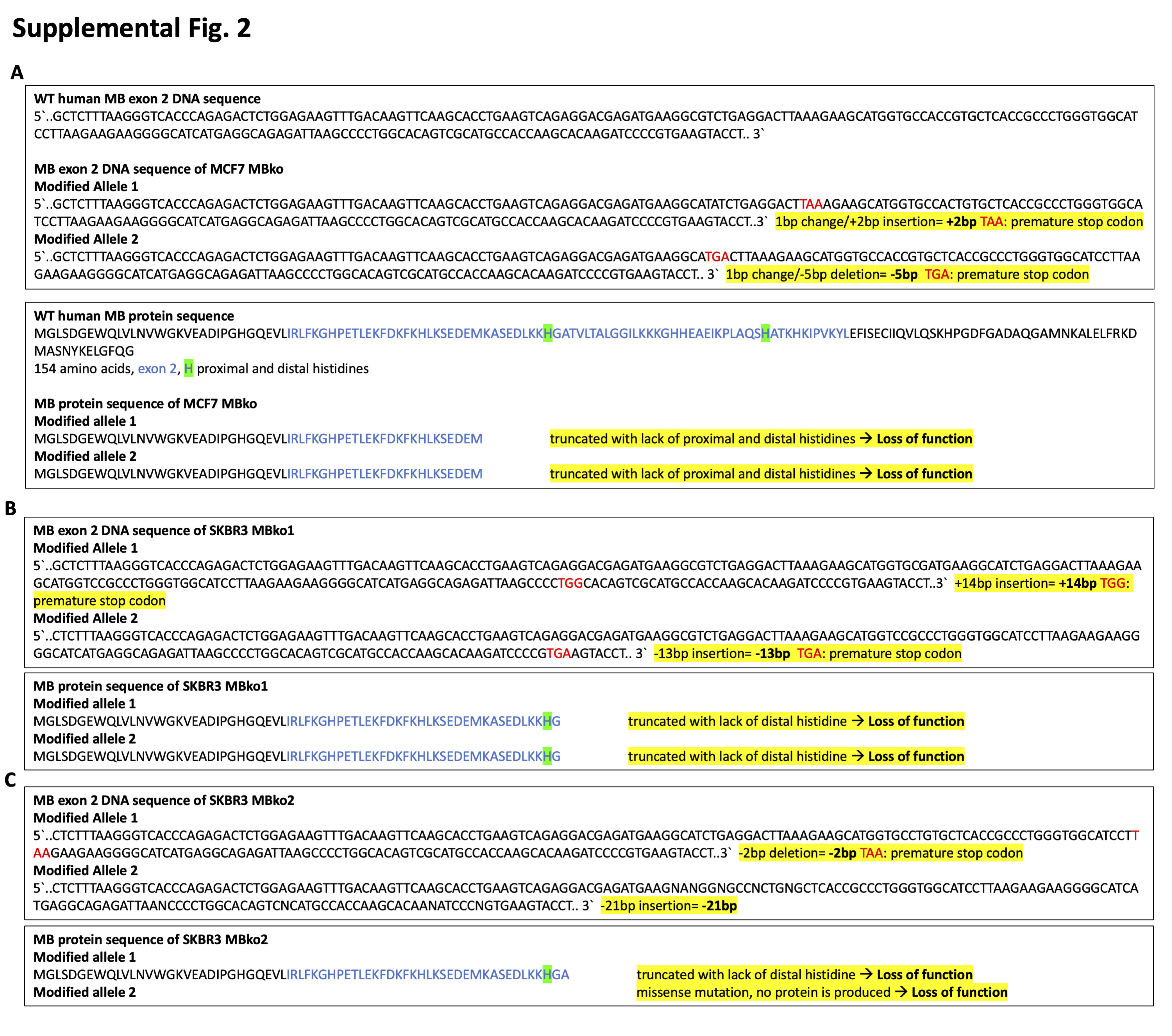
**Supplemental Figure 2. Sequencing of MCF7 and SKBR3 MB-knockout cells shows genomic alterations**

(**A**) DNA and protein sequences of wildtype human *MB* exon 2, showing induced modification with resulting genetic alteration and loss of function proteins in MCF7 MB-KO cells. (**B**) and (**C**) same as (**A**) but in SKBR3 MB-KO cells.

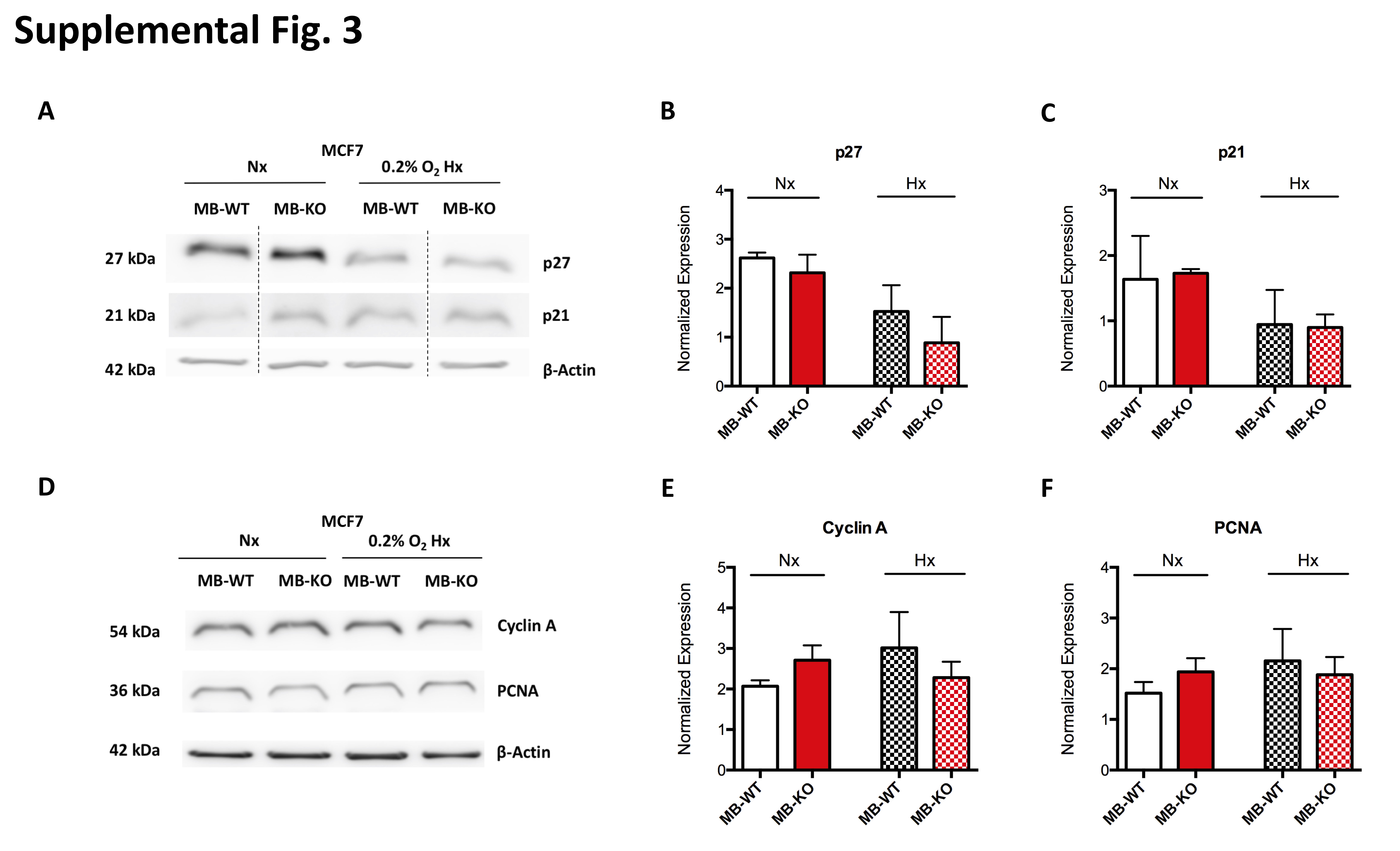
**Supplemental Figure 3. Myoglobin does not impact p27, p21, Cyclin A and PCNA proteins expressions**

(**A**) and (**D**) Representative Western Blotting image of whole tissue lysate of MB-WT and MB-KO clones of MCF7 cells cultured at normoxia Nx or 0.2% O_2_ Hx for 72h and stained for (**A**) p27 (27 kDa), p21 (21 kDa) and β-actin (42 kDa) used as loading control or (**D**) Cyclin A (54 kDa), PCNA (36kDa) and β-actin (42 kDa) used as loading control. Panels (**B**), (**C**), (**E**) and (**F**) Band intensity of p27, p21, cyclin A and PCNA proteins, respectively, after Western Blotting, from MB-WT (white) and MB-KO (red) cells of MCF7 at Nx (empty bars) and 0.2%O_2_ Hx (dashed bars), was quantified using MCID Analysis 7.0 and normalized to β-actin (n=3). Data are shown as bar graph with mean and standard error of mean and analyzed by Student`s t-test

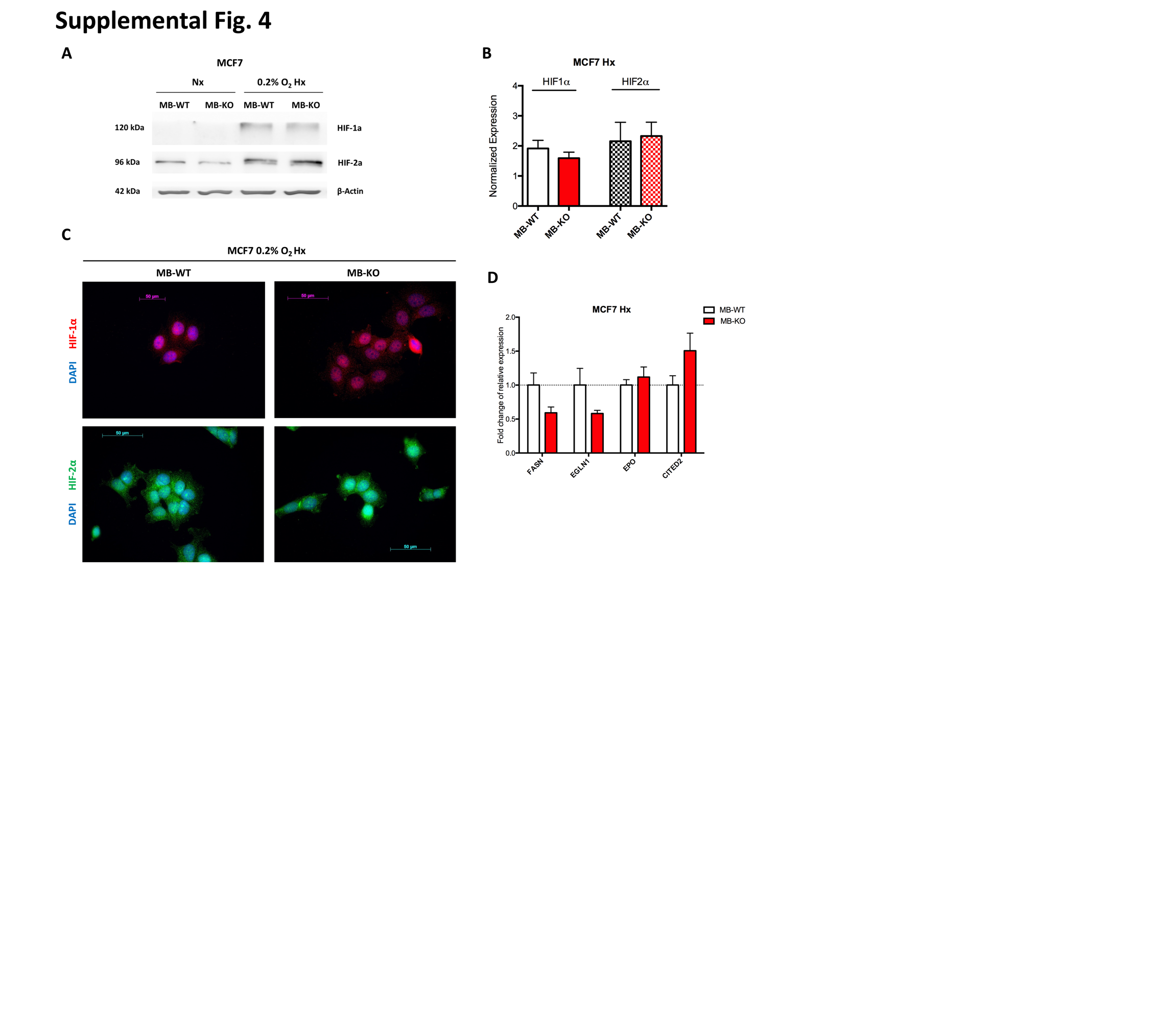

**Supplemental Figure 4. Endogenously expressed MB in BrCa cells has no impact on cellular response to hypoxia.**

(**A**) Representative Western Blotting image (n=3) of whole tissue lysate of MB-WT and MB-KO clones of MCF7 cells cultured at normoxia Nx or 0.2% O_2_ Hx for 72h and stained for hypoxia inducible factor-1a (HIF-1α) (120 kDa), hypoxia inducible factor-2a (HIF-2α) (96 kDa) and β-actin (42 kDa) used as loading control. (**B**) Band intensity of HIF-1α (empty bars) and HIF-2α (dashed bars) proteins after Western Blotting, from MB-WT (white) and MB-KO (red) cells at 0.2%O_2_ Hx for 72h, was quantified using MCID Analysis 7.0 and normalized to β-actin (n=3). (**C**) Representative immunocytochemistry images of MB-WT and MB-KO MCF7 cells stained for HIF-1α (red), HIF-2α (green) and DAPI (blue) after culturing at 0.2% O_2_ hypoxia for 72h. Scale bar is 50µm. (**D**) Relative mRNA expression levels of downstream target genes of hypoxia inducible factors: *FASN, EGLN1, EPO and CITED2*, quantified by qPCR and normalized to β-actin (*ACTB*) mRNA expression levels, from MB-WT and MB-KO clones of MCF7 cells cultured at 0.2% O_2_ Hx, for 72h. (n=3 per group). Data are shown as bar graph and are presented as mean and standard error of mean and analyzed by Student t-test ***p<0.001

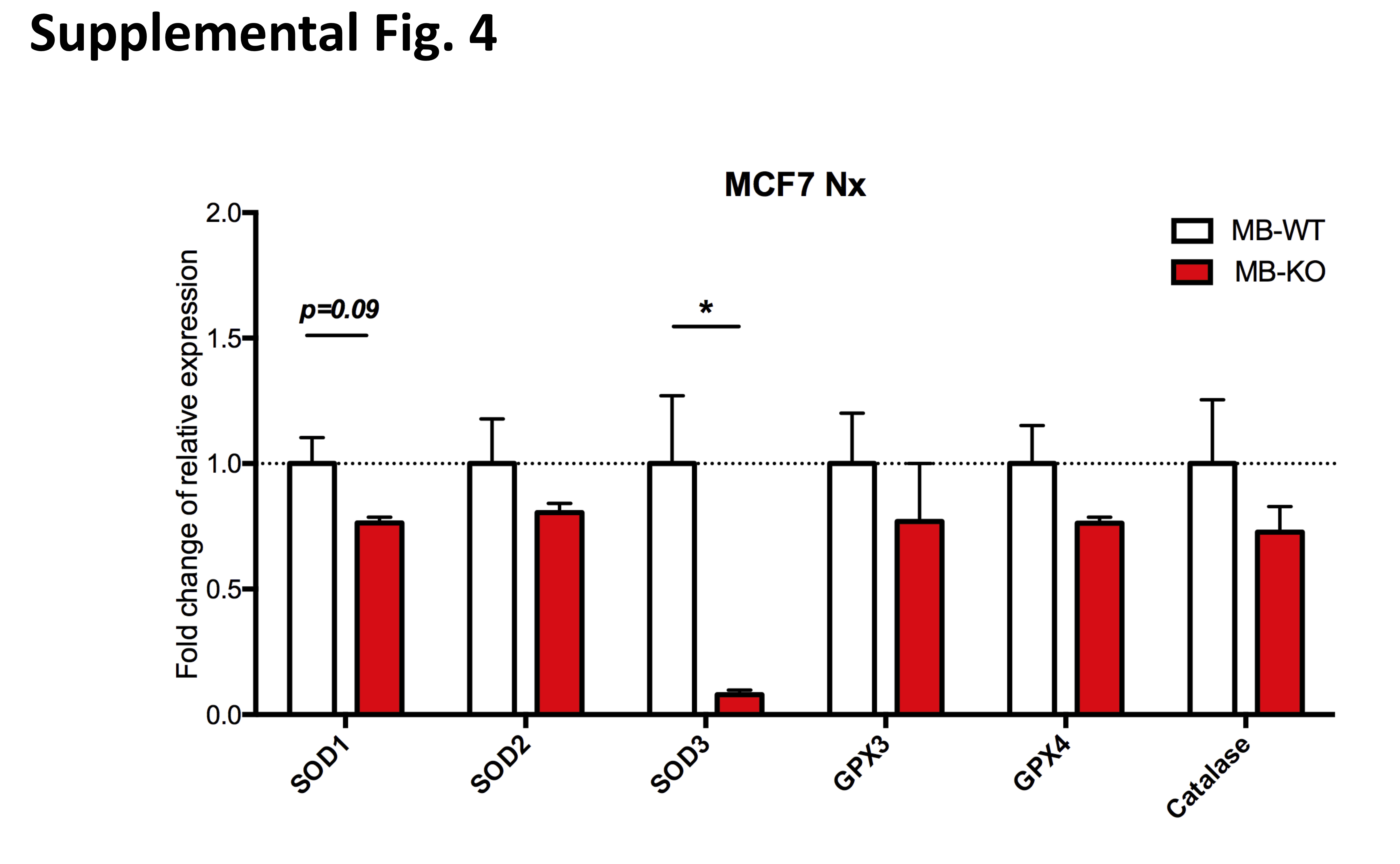

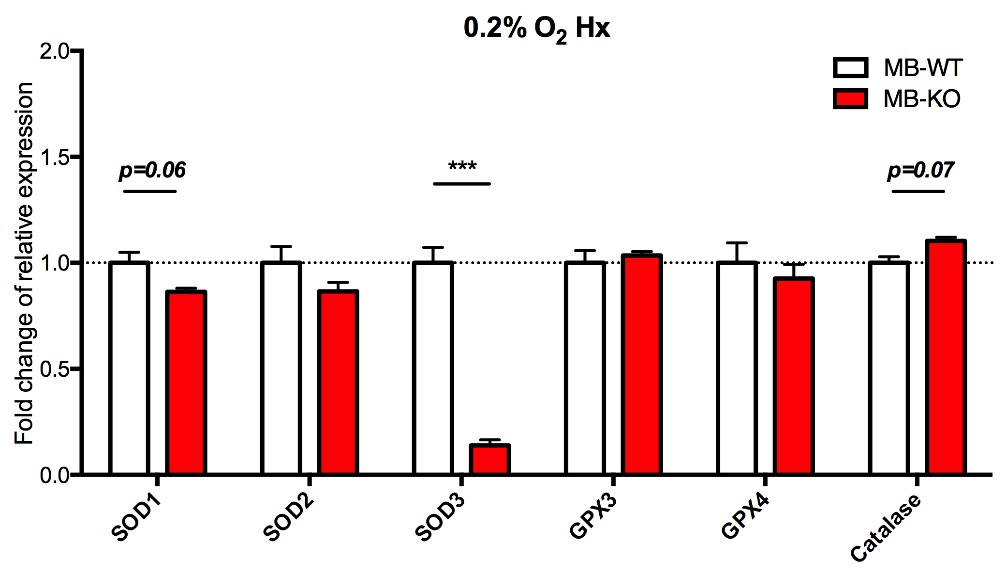

**Supplemental Figure 5. Loss of Myoglobin downregulate SOD3 antioxidant gene expression in breast cancer cells**

Relative mRNA expression levels of antioxidants genes: superoxide dismutase 1-3 (*SOD1, SOD2, SOD3*) Glutathione peroxidase 3 and 4 (*GPX3, GPX4*) and Catalase, quantified by qPCR and normalized to β-actin (*ACTB*) mRNA expression levels, from MB-WT (white bars) and MB-KO (red bars) clones of MCF7 cells cultured at normoxia (upper panel) and 0.2% O_2_ hypoxia (lower panel) for 72h (n=3 per group). Data are shown as bar graph and are presented as mean and standard error of mean and analyzed by Student’s t-test *p<0.05

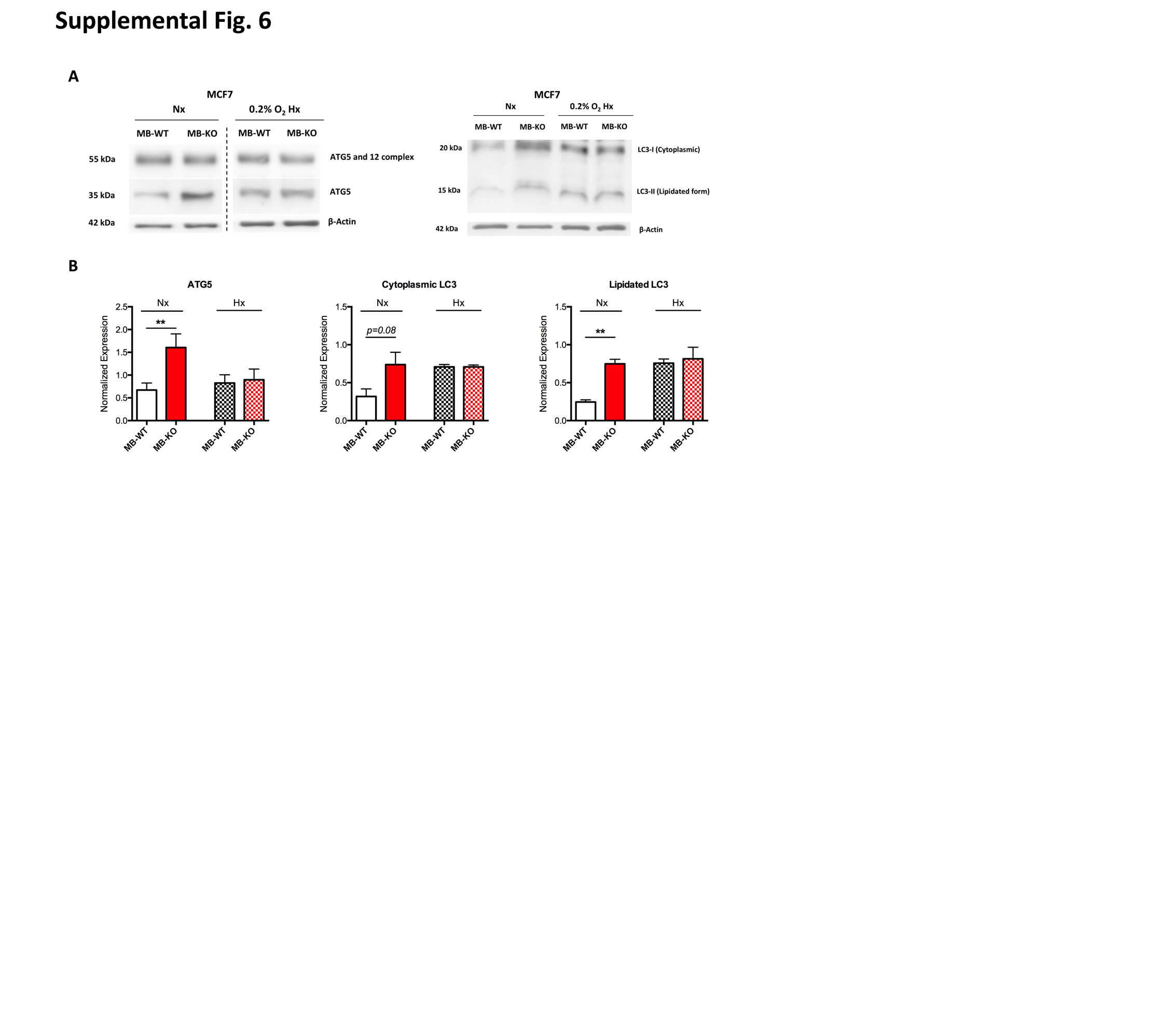

**Supplemental Figure 6. Myoglobin impacts autophagy related proteins expression in breast cancer cells.**

(**A**) Representative Western Blotting image (n=3) of whole tissue lysate of MB-WT and MB-KO clones of MCF7 cells cultured at normoxia Nx or 0.2% O_2_ Hx for 72h and stained for autophagy related protein 5 (ATG5) (35 kDa), ATG5-ATG12 complex (55 kDa), Light Chain 3 protein: cytoplasmic fraction LC3-I (20 kDa), lipidated fraction LC3-II (15 kDa) and β-actin (42 kDa) used as loading control. (**B**) Band intensity of ATG5, LC3-I and LC3-II proteins after Western Blotting, from MB-WT (white) and MB-KO (red) cells of MCF7 at Nx (empty bars) and 0.2%O_2_ Hx (dashed bars), was quantified using MCID Analysis 7.0 and normalized to β-actin (n=3). Data are shown as bar graph with mean and standard error of mean and analyzed by Student`s t-test *p<0.05; **p<0.01

**Supplementary Tables**

**Supplementary table 1.** List of all altered genes for the normoxia dataset.

| Ensembl_ID | GeneName | log2FoldChange | padj |
| --- | --- | --- | --- |
| ENSG00000136859 | ANGPTL2 | 6.497238777 | 8.29E-19 |
| ENSG00000085276 | MECOM | 1.908155009 | 1.19E-16 |
| ENSG00000170689 | HOXB9 | 2.370505195 | 2.13E-14 |
| ENSG00000102271 | KLHL4 | 2.27965335 | 4.09E-14 |
| ENSG00000186583 | SPATC1 | -2.367750837 | 2.13E-11 |
| ENSG00000003436 | TFPI | 1.877245487 | 2.13E-11 |
| ENSG00000254535 | PABPC4L | 2.067369791 | 2.13E-11 |
| ENSG00000047634 | SCML1 | 2.108920653 | 2.13E-11 |
| ENSG00000204791 | SMPD5 | -2.450028613 | 3.42E-11 |
| ENSG00000147256 | ARHGAP36 | 2.010617272 | 1.84E-09 |
| ENSG00000111799 | COL12A1 | -1.514412518 | 8.15E-09 |
| ENSG00000254337 | - | 1.186863904 | 2.37E-08 |
| ENSG00000170509 | HSD17B13 | 3.383238673 | 5.00E-08 |
| ENSG00000286322 | - | 7.950572644 | 2.33E-07 |
| ENSG00000069020 | MAST4 | 1.166467697 | 2.93E-07 |
| ENSG00000116729 | WLS | 2.472628126 | 8.19E-07 |
| ENSG00000099204 | ABLIM1 | 1.272415642 | 1.07E-06 |
| ENSG00000100867 | DHRS2 | -1.348192716 | 1.15E-06 |
| ENSG00000205426 | KRT81 | -1.605372399 | 1.33E-06 |
| ENSG00000156463 | SH3RF2 | 1.30697382 | 1.58E-06 |
| ENSG00000115221 | ITGB6 | 1.179023437 | 2.10E-06 |
| ENSG00000165025 | SYK | 0.816603319 | 2.19E-06 |
| ENSG00000177283 | FZD8 | 1.224943999 | 2.68E-06 |
| ENSG00000101986 | ABCD1 | -1.097831462 | 3.11E-06 |
| ENSG00000006747 | SCIN | 1.468030029 | 3.72E-06 |
| ENSG00000204442 | FAM155A | 1.436506374 | 5.09E-06 |
| ENSG00000142609 | CFAP74 | 1.169826068 | 5.59E-06 |
| ENSG00000164125 | GASK1B | 1.202933479 | 5.96E-06 |
| ENSG00000164749 | HNF4G | 2.461734879 | 5.96E-06 |
| ENSG00000169851 | PCDH7 | -1.282026173 | 6.53E-06 |
| ENSG00000138386 | NAB1 | 1.279430505 | 8.01E-06 |
| ENSG00000164649 | CDCA7L | 0.852166284 | 9.64E-06 |
| ENSG00000078053 | AMPH | 1.130350158 | 1.22E-05 |
| ENSG00000112246 | SIM1 | 2.237232903 | 2.13E-05 |
| ENSG00000230221 | - | 5.245912348 | 2.38E-05 |
| ENSG00000090661 | CERS4 | -0.966670875 | 2.75E-05 |
| ENSG00000106688 | SLC1A1 | -1.096463336 | 4.56E-05 |
| ENSG00000164690 | SHH | -0.783392357 | 4.56E-05 |
| ENSG00000084444 | FAM234B | -0.82863806 | 4.65E-05 |
| ENSG00000168542 | COL3A1 | 1.785512546 | 4.65E-05 |
| ENSG00000196139 | AKR1C3 | 1.277371363 | 4.97E-05 |
| ENSG00000140465 | CYP1A1 | -1.799779936 | 5.61E-05 |
| ENSG00000216740 | ANXA2P3 | 4.807368885 | 8.17E-05 |
| ENSG00000235524 | MTCO1P23 | 5.734933587 | 9.14E-05 |
| ENSG00000169403 | PTAFR | -1.204234084 | 9.25E-05 |
| ENSG00000172575 | RASGRP1 | 0.896350457 | 0.000121639 |
| ENSG00000144366 | GULP1 | 1.219525523 | 0.000133636 |
| ENSG00000242100 | RPL9P32 | 4.008219705 | 0.000148665 |
| ENSG00000105388 | CEACAM5 | 1.310571185 | 0.000177306 |
| ENSG00000249264 | EEF1A1P9 | 2.398701091 | 0.000177306 |
| ENSG00000219747 | RPL32P16 | 3.985066689 | 0.000192152 |
| ENSG00000244192 | - | 4.016477241 | 0.000207848 |
| ENSG00000019549 | SNAI2 | 1.914653463 | 0.000230149 |
| ENSG00000174473 | GALNTL6 | -1.056506923 | 0.000230266 |
| ENSG00000135111 | TBX3 | 1.579923431 | 0.000246576 |
| ENSG00000242353 | RPL12P30 | 4.488389205 | 0.000303016 |
| ENSG00000177728 | TMEM94 | -1.172667797 | 0.000339375 |
| ENSG00000260290 | - | 2.938784122 | 0.000339375 |
| ENSG00000177707 | NECTIN3 | 1.088425619 | 0.000386683 |
| ENSG00000145819 | ARHGAP26 | 0.806239319 | 0.000438049 |
| ENSG00000218426 | - | 3.952504531 | 0.000438049 |
| ENSG00000236570 | RAD23BP1 | 4.668381483 | 0.000438049 |
| ENSG00000146674 | IGFBP3 | 1.103187633 | 0.000456235 |
| ENSG00000224897 | POT1-AS1 | 1.32339549 | 0.000471726 |
| ENSG00000110934 | BIN2 | 2.175411642 | 0.000486887 |
| ENSG00000198189 | HSD17B11 | 1.364153758 | 0.000529167 |
| ENSG00000235962 | RPL7AP53 | 3.518718481 | 0.000533354 |
| ENSG00000250241 | - | 1.091674409 | 0.000560154 |
| ENSG00000099994 | SUSD2 | -1.083682308 | 0.000605718 |
| ENSG00000196932 | TMEM26 | 1.514357332 | 0.000605718 |
| ENSG00000221923 | ZNF880 | 1.313214424 | 0.000628899 |
| ENSG00000152661 | GJA1 | -1.396003408 | 0.000684844 |
| ENSG00000115590 | IL1R2 | 2.302150766 | 0.000684844 |
| ENSG00000211689 | TRGC1 | -1.589460016 | 0.000719756 |
| ENSG00000198363 | ASPH | 0.990306265 | 0.00073226 |
| ENSG00000085831 | TTC39A | -0.643551654 | 0.000735714 |
| ENSG00000130700 | GATA5 | 3.662812649 | 0.000735714 |
| ENSG00000138650 | PCDH10 | 0.707766674 | 0.000760099 |
| ENSG00000130827 | PLXNA3 | -0.880510779 | 0.000760418 |
| ENSG00000253506 | NACA2 | 3.230536532 | 0.000775503 |
| ENSG00000139793 | MBNL2 | 0.791411677 | 0.00078536 |
| ENSG00000218283 | MORF4L1P1 | 2.186681861 | 0.00078536 |
| ENSG00000226396 | - | 2.586801472 | 0.00078536 |
| ENSG00000147689 | FAM83A | -1.052500264 | 0.00079754 |
| ENSG00000182575 | NXPH3 | -0.881844501 | 0.00079754 |
| ENSG00000250148 | KRT8P31 | 3.137297838 | 0.00079754 |
| ENSG00000167925 | GHDC | -1.122806256 | 0.000806164 |
| ENSG00000152137 | HSPB8 | -0.919015119 | 0.000843577 |
| ENSG00000183049 | CAMK1D | 0.869017519 | 0.000862106 |
| ENSG00000105971 | CAV2 | 1.455190949 | 0.000876284 |
| ENSG00000144824 | PHLDB2 | 1.356770507 | 0.000881501 |
| ENSG00000189212 | DPY19L2P1 | 1.863555495 | 0.000890374 |
| ENSG00000223803 | RPS20P14 | 4.192687179 | 0.000922288 |
| ENSG00000188460 | ACTBP11 | 3.113010507 | 0.00105501 |
| ENSG00000250859 | HNRNPKP1 | 2.552120777 | 0.001265679 |
| ENSG00000143341 | HMCN1 | 0.928470086 | 0.001314871 |
| ENSG00000228499 | TMSB10P1 | 4.757400682 | 0.001345375 |
| ENSG00000180764 | PIPSL | 2.729762359 | 0.001390165 |
| ENSG00000233668 | - | 3.289256155 | 0.001390165 |
| ENSG00000179862 | CITED4 | -1.256261148 | 0.001407471 |
| ENSG00000204253 | HNRNPCP2 | 3.046251339 | 0.001407471 |
| ENSG00000108018 | SORCS1 | -1.481608293 | 0.001437959 |
| ENSG00000115641 | FHL2 | -0.796786116 | 0.001437959 |
| ENSG00000196083 | IL1RAP | 0.841637095 | 0.001452918 |
| ENSG00000232385 | RPS3AP25 | 5.005671527 | 0.001469175 |
| ENSG00000196344 | ADH7 | 3.961766052 | 0.001478951 |
| ENSG00000074855 | ANO8 | -0.65265462 | 0.001525634 |
| ENSG00000231767 | RPS27AP5 | 3.609865418 | 0.00156887 |
| ENSG00000170786 | SDR16C5 | 1.344517441 | 0.001711233 |
| ENSG00000183044 | ABAT | -0.485585083 | 0.0017634 |
| ENSG00000249092 | PPIAP77 | 4.691641365 | 0.001769923 |
| ENSG00000134668 | SPOCD1 | -1.359573419 | 0.001814703 |
| ENSG00000122644 | ARL4A | 1.123161768 | 0.001814703 |
| ENSG00000178715 | - | 3.325417444 | 0.00182207 |
| ENSG00000135549 | PKIB | -0.693523525 | 0.001835685 |
| ENSG00000143878 | RHOB | -0.650569354 | 0.001977912 |
| ENSG00000146648 | EGFR | 0.76305769 | 0.00198893 |
| ENSG00000112655 | PTK7 | -0.632518228 | 0.002021276 |
| ENSG00000160867 | FGFR4 | -0.827727153 | 0.00204014 |
| ENSG00000184809 | B3GALT5-AS1 | -1.137284769 | 0.002097184 |
| ENSG00000121552 | CSTA | -1.075593353 | 0.002172879 |
| ENSG00000237433 | RPSAP11 | 4.753033069 | 0.002174838 |
| ENSG00000145632 | PLK2 | -0.501831328 | 0.002209346 |
| ENSG00000228981 | - | 3.402845934 | 0.002209346 |
| ENSG00000233830 | EIF4HP1 | 3.583713708 | 0.002296601 |
| ENSG00000185973 | TMLHE | -0.647402841 | 0.002425823 |
| ENSG00000092758 | COL9A3 | -1.65140853 | 0.002511586 |
| ENSG00000135333 | EPHA7 | 0.945804706 | 0.002511586 |
| ENSG00000223873 | SAP18P2 | 3.909606545 | 0.002511586 |
| ENSG00000147255 | IGSF1 | 1.296410742 | 0.002537293 |
| ENSG00000146757 | ZNF92 | 0.92664326 | 0.002765914 |
| ENSG00000069667 | RORA | 1.382603025 | 0.002958463 |
| ENSG00000247763 | TUBAP14 | 3.251048421 | 0.002958463 |
| ENSG00000239622 | - | 3.598251135 | 0.002958463 |
| ENSG00000240051 | RPL23AP10 | 5.27111787 | 0.002958463 |
| ENSG00000095321 | CRAT | -0.779497345 | 0.002967373 |
| ENSG00000277443 | MARCKS | -0.591580403 | 0.00308335 |
| ENSG00000248415 | GAPDHP61 | 2.704600268 | 0.003105153 |
| ENSG00000214193 | SH3D21 | -0.5724357 | 0.00314674 |
| ENSG00000111058 | ACSS3 | 0.912031399 | 0.003302889 |
| ENSG00000249855 | EEF1A1P19 | 2.068143946 | 0.003302889 |
| ENSG00000111052 | LIN7A | 1.18096491 | 0.003378816 |
| ENSG00000230629 | RPS23P8 | 2.045316003 | 0.003993038 |
| ENSG00000013619 | MAMLD1 | -0.965448116 | 0.004011967 |
| ENSG00000102024 | PLS3 | 2.381712125 | 0.004011967 |
| ENSG00000179083 | FAM133A | -1.245266069 | 0.004079696 |
| ENSG00000228929 | RPS13P2 | 3.705910164 | 0.004090232 |
| ENSG00000240882 | - | 4.42688634 | 0.004727095 |
| ENSG00000245205 | EEF1A1P4 | 2.042503359 | 0.004751799 |
| ENSG00000283041 | - | 1.424514358 | 0.004901767 |
| ENSG00000230783 | RPS3AP13 | 3.658027714 | 0.004928121 |
| ENSG00000168843 | FSTL5 | -1.158745389 | 0.005002829 |
| ENSG00000254038 | - | 1.554115143 | 0.005116486 |
| ENSG00000164850 | GPER1 | -0.959181296 | 0.005178702 |
| ENSG00000232346 | - | 3.551134423 | 0.005178702 |
| ENSG00000116774 | OLFML3 | -1.100370809 | 0.005186402 |
| ENSG00000228834 | ATP5MFP2 | 4.925243198 | 0.005196716 |
| ENSG00000196636 | SDHAF3 | -1.514943742 | 0.005303141 |
| ENSG00000254241 | MTCO1P47 | 4.498863591 | 0.005303141 |
| ENSG00000255450 | - | 3.368729641 | 0.005451173 |
| ENSG00000235065 | RPL24P2 | 2.207403318 | 0.005484245 |
| ENSG00000233057 | EEF1A1P14 | 2.382147084 | 0.005599921 |
| ENSG00000213293 | - | 3.777378219 | 0.005599921 |
| ENSG00000099954 | CECR2 | 1.329473206 | 0.005992344 |
| ENSG00000117525 | F3 | 1.470959893 | 0.005992344 |
| ENSG00000234782 | TPT1P9 | 1.940233589 | 0.00637564 |
| ENSG00000237709 | EEF1A1P28 | 2.592000439 | 0.006622458 |
| ENSG00000139239 | RPL14P1 | 2.163965933 | 0.006667098 |
| ENSG00000219932 | RPL12P8 | 4.579332181 | 0.006851717 |
| ENSG00000143995 | MEIS1 | 0.803993311 | 0.007115745 |
| ENSG00000217889 | KRT18P48 | 2.746072725 | 0.007340391 |
| ENSG00000118922 | KLF12 | 0.813612547 | 0.007397212 |
| ENSG00000128928 | IVD | -0.502307335 | 0.007399525 |
| ENSG00000137193 | PIM1 | 0.698145337 | 0.007399525 |
| ENSG00000183346 | CABCOCO1 | 2.05425382 | 0.007399525 |
| ENSG00000240121 | RPS27P20 | 5.933269813 | 0.007399525 |
| ENSG00000154553 | PDLIM3 | 1.694268806 | 0.007428032 |
| ENSG00000103485 | QPRT | -1.678353536 | 0.007468798 |
| ENSG00000215835 | - | 3.324637792 | 0.007909283 |
| ENSG00000128422 | KRT17 | -2.224527163 | 0.007920521 |
| ENSG00000185607 | ACTBP7 | 3.136074151 | 0.008004505 |
| ENSG00000111339 | ART4 | 1.569154834 | 0.008200124 |
| ENSG00000240522 | RPL7AP10 | 2.067016399 | 0.008433607 |
| ENSG00000198753 | PLXNB3 | -1.041520769 | 0.008439501 |
| ENSG00000227578 | RPS3AP53 | 5.565143512 | 0.008439501 |
| ENSG00000058091 | CDK14 | 0.936951985 | 0.008668802 |
| ENSG00000226543 | MYL6P1 | 5.863341966 | 0.008838427 |
| ENSG00000119699 | TGFB3 | -0.678634827 | 0.008871155 |
| ENSG00000226243 | RPL37AP1 | 2.78872679 | 0.008883673 |
| ENSG00000101188 | NTSR1 | -0.842785383 | 0.009242164 |
| ENSG00000213750 | - | 4.868422039 | 0.009242164 |
| ENSG00000266602 | - | -1.7324115 | 0.009329989 |
| ENSG00000099889 | ARVCF | -0.696148033 | 0.009329989 |
| ENSG00000240622 | RPL7P15 | 3.249642894 | 0.009329989 |
| ENSG00000170545 | SMAGP | 0.827212584 | 0.009375689 |
| ENSG00000219507 | FTH1P8 | 1.774593631 | 0.009394364 |
| ENSG00000205978 | NYNRIN | -0.959789906 | 0.009413676 |
| ENSG00000253341 | PCBP2P2 | 2.186444986 | 0.009413676 |
| ENSG00000203971 | - | 5.776230155 | 0.009413676 |
| ENSG00000099256 | PRTFDC1 | 0.795507266 | 0.009494976 |
| ENSG00000216365 | RPL37P15 | 4.920851347 | 0.009520006 |
| ENSG00000238116 | RAD21P1 | 3.831686229 | 0.010033925 |
| ENSG00000127585 | FBXL16 | -0.729623046 | 0.010242732 |
| ENSG00000179918 | SEPHS2 | -0.512902904 | 0.010377263 |
| ENSG00000107518 | ATRNL1 | 0.994050619 | 0.010659626 |
| ENSG00000227121 | LINC02672 | 1.615020976 | 0.011095342 |
| ENSG00000240828 | RPL21P4 | 5.095907035 | 0.011485106 |
| ENSG00000259032 | ENSAP2 | 3.100431495 | 0.011713151 |
| ENSG00000215218 | UBE2QL1 | -1.025684813 | 0.012520138 |
| ENSG00000204287 | HLA-DRA | -1.323224749 | 0.013111032 |
| ENSG00000187017 | ESPN | -0.675830086 | 0.013111032 |
| ENSG00000111348 | ARHGDIB | 1.165789744 | 0.013111032 |
| ENSG00000169253 | - | 4.020029354 | 0.013111032 |
| ENSG00000063601 | MTMR1 | -0.470223816 | 0.01331066 |
| ENSG00000168269 | FOXI1 | -1.418331634 | 0.013537674 |
| ENSG00000164292 | RHOBTB3 | 0.782931885 | 0.013644114 |
| ENSG00000147955 | SIGMAR1 | -0.916056083 | 0.014080568 |
| ENSG00000254416 | LINC02732 | -0.935101571 | 0.015055435 |
| ENSG00000173846 | PLK3 | -0.838196617 | 0.015241081 |
| ENSG00000241258 | CRCP | 0.423543218 | 0.015241081 |
| ENSG00000076356 | PLXNA2 | 0.964900772 | 0.015241081 |
| ENSG00000139211 | AMIGO2 | 1.043845458 | 0.015346655 |
| ENSG00000107957 | SH3PXD2A | 0.69644889 | 0.015574357 |
| ENSG00000071859 | FAM50A | -0.63153829 | 0.01642141 |
| ENSG00000104549 | SQLE | -0.427854588 | 0.01642141 |
| ENSG00000187122 | SLIT1 | -1.215226808 | 0.016833947 |
| ENSG00000197008 | ZNF138 | 0.597633962 | 0.017120482 |
| ENSG00000228818 | - | 2.682027264 | 0.017120482 |
| ENSG00000106609 | TMEM248 | 0.370311854 | 0.017159749 |
| ENSG00000228247 | UBBP2 | 2.957209864 | 0.017533719 |
| ENSG00000242607 | RPS3AP34 | 6.097316364 | 0.017622727 |
| ENSG00000173852 | DPY19L1 | 0.678400585 | 0.017636161 |
| ENSG00000196437 | ZNF569 | -1.731047377 | 0.017639448 |
| ENSG00000228462 | RPS19P7 | 4.15142014 | 0.017639448 |
| ENSG00000100346 | CACNA1I | -0.939490968 | 0.017905363 |
| ENSG00000049089 | COL9A2 | -0.716433064 | 0.017905363 |
| ENSG00000236946 | HNRNPA1P70 | 4.220827587 | 0.017905363 |
| ENSG00000269516 | CYP4F23P | -1.355200061 | 0.018176628 |
| ENSG00000228232 | GAPDHP1 | 1.926244389 | 0.018249316 |
| ENSG00000234067 | RPL5P10 | 5.256489928 | 0.018249316 |
| ENSG00000157303 | SUSD3 | -0.785217633 | 0.01826698 |
| ENSG00000224899 | LINC02830 | 1.378909107 | 0.018290874 |
| ENSG00000127954 | STEAP4 | -1.10247217 | 0.018468408 |
| ENSG00000224411 | HSP90AA2P | 2.835160673 | 0.018468408 |
| ENSG00000215388 | ACTG1P3 | 3.173071442 | 0.018468408 |
| ENSG00000102312 | PORCN | -0.650335557 | 0.018669646 |
| ENSG00000183778 | B3GALT5 | -0.983935411 | 0.018819701 |
| ENSG00000254270 | ERHP1 | 4.157242204 | 0.018819701 |
| ENSG00000259078 | PTBP1P | 5.687193611 | 0.018819701 |
| ENSG00000197121 | PGAP1 | 1.001153218 | 0.018853073 |
| ENSG00000103197 | TSC2 | -0.522476006 | 0.01919956 |
| ENSG00000213790 | OLA1P1 | 2.498760921 | 0.01919956 |
| ENSG00000102125 | TAZ | -0.66520731 | 0.019442471 |
| ENSG00000197774 | EME2 | -0.614885155 | 0.019442471 |
| ENSG00000184271 | POU6F1 | 1.17600574 | 0.019442471 |
| ENSG00000214460 | TPT1P6 | 2.410441562 | 0.019442471 |
| ENSG00000226549 | SCDP1 | 3.315406452 | 0.019442471 |
| ENSG00000227615 | - | 3.403343099 | 0.019442471 |
| ENSG00000280195 | - | -0.765248941 | 0.019539965 |
| ENSG00000227968 | BUB3P1 | 4.316965811 | 0.020335929 |
| ENSG00000127955 | GNAI1 | -1.059385072 | 0.020356509 |
| ENSG00000270706 | PRMT1P1 | 3.454672059 | 0.020449478 |
| ENSG00000231181 | - | 2.984186863 | 0.020823463 |
| ENSG00000240535 | - | 4.887889583 | 0.020823463 |
| ENSG00000198406 | BZW1P2 | 2.054100395 | 0.021161179 |
| ENSG00000242411 | - | 2.791384264 | 0.021165677 |
| ENSG00000071553 | ATP6AP1 | -0.615086467 | 0.021198717 |
| ENSG00000230807 | - | 4.185157487 | 0.021198717 |
| ENSG00000114450 | GNB4 | 1.449291301 | 0.021323853 |
| ENSG00000236480 | PKMP1 | 2.497019359 | 0.021323853 |
| ENSG00000230916 | MTCO1P53 | 1.366620002 | 0.02175371 |
| ENSG00000242327 | - | 2.885314351 | 0.021781039 |
| ENSG00000277043 | EEF1A1P42 | 3.046001874 | 0.021865622 |
| ENSG00000234332 | BCAS2P2 | 3.019850599 | 0.021904441 |
| ENSG00000152217 | SETBP1 | 1.292835226 | 0.022309564 |
| ENSG00000228887 | EEF1DP1 | 2.1392636 | 0.022392867 |
| ENSG00000259706 | HSP90B2P | 1.863842508 | 0.022407127 |
| ENSG00000113594 | LIFR | 0.949861413 | 0.022720959 |
| ENSG00000005001 | PRSS22 | -0.718451213 | 0.023054112 |
| ENSG00000214078 | CPNE1 | -0.625904009 | 0.023204676 |
| ENSG00000113580 | NR3C1 | 0.733982059 | 0.023717783 |
| ENSG00000225356 | - | 2.570596956 | 0.023792208 |
| ENSG00000183665 | TRMT12 | -0.446000179 | 0.023802567 |
| ENSG00000164236 | ANKRD33B | 1.295901116 | 0.024034587 |
| ENSG00000213891 | RPL3P6 | 2.910977628 | 0.024034587 |
| ENSG00000223668 | EEF1A1P24 | 1.390348206 | 0.02405663 |
| ENSG00000232054 | NPM1P34 | 4.027888676 | 0.024209386 |
| ENSG00000149573 | MPZL2 | 0.624298682 | 0.024327882 |
| ENSG00000163513 | TGFBR2 | 0.687717999 | 0.025182507 |
| ENSG00000172974 | VDAC2P5 | 3.040203602 | 0.02557041 |
| ENSG00000059122 | FLYWCH1 | -0.535597772 | 0.026359973 |
| ENSG00000132329 | RAMP1 | -1.151709045 | 0.026496047 |
| ENSG00000124762 | CDKN1A | -0.816297241 | 0.026867409 |
| ENSG00000230146 | SEPHS1P4 | 2.824122165 | 0.027318018 |
| ENSG00000162004 | CCDC78 | -0.609386531 | 0.027888671 |
| ENSG00000244503 | - | 2.927566729 | 0.027888671 |
| ENSG00000213613 | RPL11P3 | 2.073794128 | 0.027954729 |
| ENSG00000162066 | AMDHD2 | -0.746617285 | 0.028088891 |
| ENSG00000263266 | RPS7P1 | 3.304406215 | 0.028088891 |
| ENSG00000146700 | SSC4D | -1.077006632 | 0.028297426 |
| ENSG00000165905 | LARGE2 | -0.70083752 | 0.028297426 |
| ENSG00000196715 | VKORC1L1 | 0.523888854 | 0.028297426 |
| ENSG00000223529 | EEF1A1P8 | 1.565219032 | 0.028297426 |
| ENSG00000223739 | RPS15AP15 | 2.48829157 | 0.028297426 |
| ENSG00000248373 | - | -1.915693033 | 0.028361297 |
| ENSG00000145451 | GLRA3 | 0.953610434 | 0.028377457 |
| ENSG00000165895 | ARHGAP42 | 0.763993188 | 0.028740261 |
| ENSG00000234882 | EIF3EP1 | 1.805475621 | 0.02932064 |
| ENSG00000187957 | DNER | -1.689575438 | 0.029559212 |
| ENSG00000218582 | GAPDHP63 | 2.333091969 | 0.029736285 |
| ENSG00000233111 | RAB1C | 2.929796171 | 0.029736285 |
| ENSG00000224773 | HSPA8P7 | 3.521345274 | 0.029736285 |
| ENSG00000250363 | KRT18P21 | 2.892056218 | 0.029934936 |
| ENSG00000233476 | EEF1A1P6 | 2.372856072 | 0.030843593 |
| ENSG00000225568 | - | 3.690815517 | 0.031658775 |
| ENSG00000283057 | - | 4.528258581 | 0.031999332 |
| ENSG00000100842 | EFS | -1.330019831 | 0.032803633 |
| ENSG00000171798 | KNDC1 | -0.573539678 | 0.03305499 |
| ENSG00000213704 | EEF1A1P15 | 2.384751765 | 0.034213815 |
| ENSG00000227051 | C14orf132 | 1.229861066 | 0.034315347 |
| ENSG00000225536 | STIP1P3 | 3.815814155 | 0.034629087 |
| ENSG00000117859 | OSBPL9 | -0.437030595 | 0.035140735 |
| ENSG00000250144 | - | 1.898506926 | 0.035140735 |
| ENSG00000262152 | LINC00514 | -0.824478928 | 0.035352576 |
| ENSG00000218175 | - | 2.440967361 | 0.035352576 |
| ENSG00000230391 | RPSAP23 | 3.711109477 | 0.035352576 |
| ENSG00000234589 | - | 3.606304299 | 0.035479072 |
| ENSG00000146072 | TNFRSF21 | 0.424860354 | 0.035623349 |
| ENSG00000179715 | PCED1B | 1.022584521 | 0.035623349 |
| ENSG00000101276 | SLC52A3 | -0.682977192 | 0.036331725 |
| ENSG00000069974 | RAB27A | 0.763178171 | 0.036331725 |
| ENSG00000231494 | RPL21P35 | 4.719297597 | 0.036435016 |
| ENSG00000170153 | RNF150 | 0.776678978 | 0.037092688 |
| ENSG00000135766 | EGLN1 | -0.601030685 | 0.037286484 |
| ENSG00000257616 | - | 3.292847871 | 0.037286484 |
| ENSG00000064787 | BCAS1 | 1.968671816 | 0.037702376 |
| ENSG00000267398 | - | 4.86359512 | 0.037870512 |
| ENSG00000197044 | ZNF441 | -2.33601766 | 0.037981597 |
| ENSG00000123472 | ATPAF1 | -0.355171675 | 0.037981597 |
| ENSG00000151846 | PABPC3 | 2.307863355 | 0.037981597 |
| ENSG00000185825 | BCAP31 | -0.597966213 | 0.038173178 |
| ENSG00000179698 | WDR97 | -0.734068206 | 0.038768233 |
| ENSG00000242683 | RPL12P21 | 4.259733953 | 0.039259136 |
| ENSG00000115129 | TP53I3 | -0.781413595 | 0.039368471 |
| ENSG00000137573 | SULF1 | 0.773326094 | 0.039368471 |
| ENSG00000223810 | KRT8P28 | 5.037443263 | 0.039391096 |
| ENSG00000184226 | PCDH9 | 0.520495288 | 0.039856514 |
| ENSG00000240480 | RPL29P2 | 2.753668271 | 0.04048991 |
| ENSG00000132746 | ALDH3B2 | -0.774910521 | 0.040972325 |
| ENSG00000167962 | ZNF598 | -0.482599097 | 0.041518364 |
| ENSG00000197043 | ANXA6 | -0.733185227 | 0.041586235 |
| ENSG00000213820 | RPL13P2 | 2.88606589 | 0.041586235 |
| ENSG00000004948 | CALCR | 1.150273569 | 0.041707306 |
| ENSG00000242291 | RPL36AP51 | 2.544627727 | 0.041707306 |
| ENSG00000227309 | - | 3.213063253 | 0.041707306 |
| ENSG00000251333 | RTN3P1 | 3.435185086 | 0.041707306 |
| ENSG00000243064 | ABCC13 | 5.059031564 | 0.041707306 |
| ENSG00000167768 | KRT1 | 2.738495237 | 0.041949884 |
| ENSG00000216713 | MTND4P13 | 5.327619491 | 0.041949884 |
| ENSG00000100003 | SEC14L2 | -0.800313034 | 0.042679684 |
| ENSG00000167996 | FTH1 | 0.665237636 | 0.042785064 |
| ENSG00000178028 | DMAP1 | -0.373834026 | 0.043141034 |
| ENSG00000196814 | MVB12B | 0.665158879 | 0.043141034 |
| ENSG00000242299 | - | 1.646650805 | 0.043141034 |
| ENSG00000224333 | GAPDHP20 | 3.544311814 | 0.043141034 |
| ENSG00000270553 | - | 4.02133657 | 0.043141034 |
| ENSG00000220472 | - | 3.019992559 | 0.043271716 |
| ENSG00000265480 | KRT18P55 | 3.122907463 | 0.043271716 |
| ENSG00000243547 | HNRNPKP4 | 1.841565268 | 0.043302499 |
| ENSG00000213601 | KRT18P19 | 2.467937009 | 0.04362776 |
| ENSG00000151692 | RNF144A | 0.817176493 | 0.04378853 |
| ENSG00000117385 | P3H1 | -0.674163707 | 0.043794529 |
| ENSG00000168140 | VASN | -0.893799166 | 0.043943537 |
| ENSG00000234785 | EEF1GP5 | 2.010451742 | 0.044073584 |
| ENSG00000260711 | - | 1.022344849 | 0.044077435 |
| ENSG00000225971 | RPS3AP51 | 3.451834984 | 0.044077435 |
| ENSG00000102683 | SGCG | -0.941271962 | 0.044318398 |
| ENSG00000092929 | UNC13D | -0.552547603 | 0.045166763 |
| ENSG00000132561 | MATN2 | -0.459894097 | 0.046008225 |
| ENSG00000236686 | BZW1P1 | 3.134549276 | 0.046008225 |
| ENSG00000232042 | - | 4.68501998 | 0.046226247 |
| ENSG00000232493 | RPL12P11 | 2.716425319 | 0.046356949 |
| ENSG00000167967 | E4F1 | -0.505564179 | 0.046711014 |
| ENSG00000138821 | SLC39A8 | -1.005540128 | 0.04717101 |
| ENSG00000154274 | C4orf19 | 0.537463886 | 0.04726818 |
| ENSG00000145824 | CXCL14 | -2.432576793 | 0.047492552 |
| ENSG00000108679 | LGALS3BP | -0.575857397 | 0.047492552 |
| ENSG00000008256 | CYTH3 | 0.401236013 | 0.047492552 |
| ENSG00000232883 | - | 2.077703722 | 0.047492552 |
| ENSG00000261557 | EEF1A1P38 | 1.817046237 | 0.047551752 |
| ENSG00000184100 | BRD7P2 | 4.445268476 | 0.047685988 |
| ENSG00000089820 | ARHGAP4 | -0.590067909 | 0.047949638 |
| ENSG00000160293 | VAV2 | -0.392153137 | 0.047949638 |
| ENSG00000226581 | LINC02848 | 2.313287097 | 0.047949638 |
| ENSG00000255642 | PABPC1P4 | 1.8582801 | 0.048064447 |
| ENSG00000258162 | - | 2.992999721 | 0.048532021 |
| ENSG00000165731 | RET | -0.656388536 | 0.048731744 |
| ENSG00000236937 | PTGES3P4 | 3.698104157 | 0.048731744 |
| ENSG00000268282 | - | 3.090519817 | 0.048913995 |
| ENSG00000151572 | ANO4 | 4.560214531 | 0.048937848 |
| ENSG00000254387 | MYL12AP1 | 4.626925986 | 0.04906469 |
| ENSG00000181085 | MAPK15 | -0.682094944 | 0.049123345 |
| ENSG00000173917 | HOXB2 | 0.954410081 | 0.049346147 |
| ENSG00000141527 | CARD14 | -0.521667592 | 0.049377932 |
| ENSG00000163017 | ACTG2 | -1.044928379 | 0.049958593 |
| ENSG00000090674 | MCOLN1 | -0.602387236 | 0.049958593 |
| ENSG00000115594 | IL1R1 | 0.895353232 | 0.049958593 |
| ENSG00000232187 | FTH1P7 | 1.874467012 | 0.049958593 |
| ENSG00000243094 | RPL32P2 | 2.443087852 | 0.049958593 |

**Supplementary table 2.** List of all altered genes for the hypoxia dataset.

| Ensembl_ID | GeneName | log2FoldChange | padj |
| --- | --- | --- | --- |
| ENSG00000115221 | ITGB6 | 1.887537323 | 1.17E-22 |
| ENSG00000003436 | TFPI | 2.172972248 | 6.06E-17 |
| ENSG00000146674 | IGFBP3 | 1.773300934 | 5.30E-13 |
| ENSG00000165025 | SYK | 1.163683373 | 5.66E-13 |
| ENSG00000154229 | PRKCA | 1.163129691 | 2.21E-10 |
| ENSG00000164683 | HEY1 | 1.481845222 | 6.36E-10 |
| ENSG00000172575 | RASGRP1 | 1.265181902 | 8.15E-10 |
| ENSG00000169851 | PCDH7 | -1.647079526 | 8.38E-10 |
| ENSG00000085276 | MECOM | 1.51082496 | 2.20E-08 |
| ENSG00000136859 | ANGPTL2 | 8.129998244 | 3.77E-08 |
| ENSG00000164749 | HNF4G | 2.636267727 | 3.91E-08 |
| ENSG00000196083 | IL1RAP | 1.197383809 | 5.39E-08 |
| ENSG00000142609 | CFAP74 | 1.399515499 | 5.67E-08 |
| ENSG00000146072 | TNFRSF21 | 0.774901922 | 7.61E-08 |
| ENSG00000116729 | WLS | 2.827810124 | 7.61E-08 |
| ENSG00000154783 | FGD5 | 1.965559587 | 1.06E-07 |
| ENSG00000111799 | COL12A1 | -1.540203088 | 1.13E-07 |
| ENSG00000186583 | SPATC1 | -1.931076585 | 1.41E-07 |
| ENSG00000047634 | SCML1 | 1.65194411 | 1.28E-06 |
| ENSG00000146648 | EGFR | 0.948714015 | 1.55E-06 |
| ENSG00000144824 | PHLDB2 | 1.727146526 | 1.61E-06 |
| ENSG00000164292 | RHOBTB3 | 1.207442733 | 4.87E-06 |
| ENSG00000198918 | RPL39 | 1.450868842 | 4.87E-06 |
| ENSG00000012779 | ALOX5 | 1.597723393 | 1.13E-05 |
| ENSG00000058085 | LAMC2 | 0.806616786 | 1.14E-05 |
| ENSG00000121552 | CSTA | -1.416173976 | 1.34E-05 |
| ENSG00000006747 | SCIN | 1.420886012 | 1.34E-05 |
| ENSG00000231298 | MANCR | 2.51434657 | 1.80E-05 |
| ENSG00000183696 | UPP1 | 1.58692632 | 2.45E-05 |
| ENSG00000182795 | C1orf116 | 1.600709084 | 4.38E-05 |
| ENSG00000115919 | KYNU | -1.042762853 | 4.89E-05 |
| ENSG00000107159 | CA9 | -1.421993728 | 4.93E-05 |
| ENSG00000169071 | ROR2 | 1.417465636 | 4.93E-05 |
| ENSG00000135111 | TBX3 | 1.598241824 | 4.93E-05 |
| ENSG00000064042 | LIMCH1 | 1.267690171 | 5.04E-05 |
| ENSG00000152137 | HSPB8 | -1.134438072 | 5.14E-05 |
| ENSG00000178726 | THBD | 1.626756766 | 5.14E-05 |
| ENSG00000107263 | RAPGEF1 | 0.671545037 | 5.76E-05 |
| ENSG00000285969 | - | 2.097187313 | 5.76E-05 |
| ENSG00000286322 | - | 6.518467235 | 6.50E-05 |
| ENSG00000163513 | TGFBR2 | 1.007679091 | 6.65E-05 |
| ENSG00000140465 | CYP1A1 | -1.56107289 | 9.77E-05 |
| ENSG00000076641 | PAG1 | 1.563945474 | 0.000111832 |
| ENSG00000235123 | DSCAM-AS1 | -0.790080967 | 0.000113593 |
| ENSG00000057657 | PRDM1 | 2.329038489 | 0.000145353 |
| ENSG00000204054 | LINC00963 | 0.783798549 | 0.000179915 |
| ENSG00000204791 | SMPD5 | -1.633123343 | 0.000188298 |
| ENSG00000111052 | LIN7A | 1.33683013 | 0.000188298 |
| ENSG00000170689 | HOXB9 | 1.905768603 | 0.000197864 |
| ENSG00000124225 | PMEPA1 | 0.850369061 | 0.000206504 |
| ENSG00000205426 | KRT81 | -1.316479878 | 0.00021194 |
| ENSG00000046604 | DSG2 | 0.787711198 | 0.000225622 |
| ENSG00000113532 | ST8SIA4 | -1.050562365 | 0.000264609 |
| ENSG00000074527 | NTN4 | 1.006042339 | 0.000264609 |
| ENSG00000080031 | PTPRH | 1.042134877 | 0.000281654 |
| ENSG00000266602 | - | -2.018787766 | 0.00028389 |
| ENSG00000168843 | FSTL5 | -1.800256286 | 0.000357038 |
| ENSG00000019549 | SNAI2 | 1.698272988 | 0.000362594 |
| ENSG00000187122 | SLIT1 | -1.620709612 | 0.000375597 |
| ENSG00000167925 | GHDC | -1.333917599 | 0.000395162 |
| ENSG00000138386 | NAB1 | 1.082406341 | 0.000442487 |
| ENSG00000101188 | NTSR1 | -1.042061506 | 0.000550814 |
| ENSG00000128052 | KDR | 1.534894225 | 0.000623533 |
| ENSG00000138640 | FAM13A | 1.182037366 | 0.000662563 |
| ENSG00000198363 | ASPH | 0.979805823 | 0.0007227 |
| ENSG00000108018 | SORCS1 | -1.489535649 | 0.00077885 |
| ENSG00000147010 | SH3KBP1 | 0.572321065 | 0.00077885 |
| ENSG00000091136 | LAMB1 | 1.191092795 | 0.000899757 |
| ENSG00000254038 | - | 3.003212661 | 0.001010642 |
| ENSG00000173930 | SLCO4C1 | -1.288079259 | 0.001082896 |
| ENSG00000221923 | ZNF880 | 1.335263788 | 0.001082896 |
| ENSG00000168542 | COL3A1 | 1.602811449 | 0.001679115 |
| ENSG00000106031 | HOXA13 | 1.016363848 | 0.001943552 |
| ENSG00000137193 | PIM1 | 0.769454184 | 0.001981872 |
| ENSG00000053747 | LAMA3 | 0.907185051 | 0.001981872 |
| ENSG00000105559 | PLEKHA4 | -1.466672125 | 0.00198545 |
| ENSG00000116285 | ERRFI1 | 0.685058114 | 0.002008162 |
| ENSG00000205413 | SAMD9 | 1.418108679 | 0.002008162 |
| ENSG00000080200 | CRYBG3 | 1.682785382 | 0.002071108 |
| ENSG00000249267 | LINC00939 | -3.075744737 | 0.002190921 |
| ENSG00000166949 | SMAD3 | 0.584936666 | 0.002190921 |
| ENSG00000145819 | ARHGAP26 | 0.739434387 | 0.002190921 |
| ENSG00000106609 | TMEM248 | 0.440441705 | 0.002273963 |
| ENSG00000113594 | LIFR | 1.206832965 | 0.002273963 |
| ENSG00000167767 | KRT80 | 0.649454647 | 0.002723742 |
| ENSG00000156103 | MMP16 | -1.525939945 | 0.002925458 |
| ENSG00000176903 | PNMA1 | 0.501869333 | 0.002925458 |
| ENSG00000258676 | - | 1.54566665 | 0.002925458 |
| ENSG00000106258 | CYP3A5 | 1.55131228 | 0.002925458 |
| ENSG00000260604 | - | 1.204819417 | 0.003056491 |
| ENSG00000224897 | POT1-AS1 | 1.22132436 | 0.003094074 |
| ENSG00000127329 | PTPRB | 1.921897774 | 0.003094074 |
| ENSG00000130700 | GATA5 | 5.496109822 | 0.003113568 |
| ENSG00000162004 | CCDC78 | -0.773688636 | 0.003140284 |
| ENSG00000115935 | WIPF1 | 1.458618106 | 0.003261989 |
| ENSG00000168280 | KIF5C | -1.028103024 | 0.003296233 |
| ENSG00000140479 | PCSK6 | 0.669343311 | 0.00333866 |
| ENSG00000086548 | CEACAM6 | 1.109644795 | 0.003627968 |
| ENSG00000145623 | OSMR | 0.716953336 | 0.0040397 |
| ENSG00000197261 | C6orf141 | -1.012853368 | 0.004144454 |
| ENSG00000156535 | CD109 | 0.848080826 | 0.004144454 |
| ENSG00000118523 | CCN2 | 1.482978194 | 0.004161113 |
| ENSG00000112655 | PTK7 | -0.629473806 | 0.00420601 |
| ENSG00000100346 | CACNA1I | -0.963970906 | 0.004270001 |
| ENSG00000151632 | AKR1C2 | -1.016893989 | 0.004483458 |
| ENSG00000150961 | SEC24D | 0.545709369 | 0.005477395 |
| ENSG00000227121 | LINC02672 | 0.909713024 | 0.005531037 |
| ENSG00000164120 | HPGD | 1.771775904 | 0.005531037 |
| ENSG00000171827 | ZNF570 | -1.077090746 | 0.005565237 |
| ENSG00000120708 | TGFBI | 0.695960814 | 0.005565237 |
| ENSG00000198189 | HSD17B11 | 1.246374525 | 0.005565237 |
| ENSG00000145779 | TNFAIP8 | 0.746490847 | 0.005749465 |
| ENSG00000168874 | ATOH8 | 0.848818574 | 0.005901578 |
| ENSG00000174473 | GALNTL6 | -1.001236023 | 0.005902948 |
| ENSG00000266074 | BAHCC1 | -0.64739927 | 0.005919479 |
| ENSG00000204740 | MALRD1 | -1.200847317 | 0.005925588 |
| ENSG00000092969 | TGFB2 | 1.190774446 | 0.005937038 |
| ENSG00000180185 | FAHD1 | -0.553978655 | 0.006076574 |
| ENSG00000106635 | BCL7B | 0.658565814 | 0.006180558 |
| ENSG00000183049 | CAMK1D | 0.740456408 | 0.006628651 |
| ENSG00000076356 | PLXNA2 | 1.060161503 | 0.007071145 |
| ENSG00000116016 | EPAS1 | 0.710660721 | 0.007310133 |
| ENSG00000177707 | NECTIN3 | 0.928397449 | 0.008001667 |
| ENSG00000164236 | ANKRD33B | 1.298113022 | 0.008438208 |
| ENSG00000159840 | ZYX | 0.59033282 | 0.008713077 |
| ENSG00000196814 | MVB12B | 0.82873454 | 0.008713077 |
| ENSG00000185442 | FAM174B | 0.654451371 | 0.009151961 |
| ENSG00000152270 | PDE3B | 0.715891537 | 0.00947575 |
| ENSG00000069667 | RORA | 1.162401449 | 0.009672809 |
| ENSG00000105137 | SYDE1 | 0.649403117 | 0.009693199 |
| ENSG00000286523 | - | -2.077053571 | 0.009696514 |
| ENSG00000181104 | F2R | 1.173707777 | 0.009853825 |
| ENSG00000103485 | QPRT | -1.653053249 | 0.009875314 |
| ENSG00000198948 | MFAP3L | -1.016024851 | 0.009875314 |
| ENSG00000069020 | MAST4 | 0.736737671 | 0.010420293 |
| ENSG00000185736 | ADARB2 | 2.422037065 | 0.010472457 |
| ENSG00000139793 | MBNL2 | 0.666241554 | 0.010781585 |
| ENSG00000082684 | SEMA5B | 1.027668116 | 0.010781585 |
| ENSG00000224899 | LINC02830 | 1.527838579 | 0.010781585 |
| ENSG00000116774 | OLFML3 | -1.304608755 | 0.010805549 |
| ENSG00000147955 | SIGMAR1 | -1.157881403 | 0.010805549 |
| ENSG00000180938 | ZNF572 | -0.619719575 | 0.011983847 |
| ENSG00000076716 | GPC4 | 0.918676078 | 0.011983847 |
| ENSG00000150907 | FOXO1 | 0.752117822 | 0.012268421 |
| ENSG00000164850 | GPER1 | -0.96672716 | 0.012499516 |
| ENSG00000214293 | APTR | 0.66005604 | 0.012852333 |
| ENSG00000157404 | KIT | -1.328505763 | 0.012897939 |
| ENSG00000141527 | CARD14 | -0.598080812 | 0.012995421 |
| ENSG00000171617 | ENC1 | 0.563968494 | 0.012995421 |
| ENSG00000144290 | SLC4A10 | -1.159528989 | 0.013028973 |
| ENSG00000145451 | GLRA3 | 1.055015513 | 0.013028973 |
| ENSG00000237187 | NR2F1-AS1 | 3.764906575 | 0.014042161 |
| ENSG00000090661 | CERS4 | -0.758894532 | 0.014576441 |
| ENSG00000101986 | ABCD1 | -0.759319392 | 0.01484599 |
| ENSG00000003096 | KLHL13 | 1.071467939 | 0.01484599 |
| ENSG00000197696 | NMB | 1.008653941 | 0.015601172 |
| ENSG00000083720 | OXCT1 | -0.816172872 | 0.015972041 |
| ENSG00000214193 | SH3D21 | -0.504225322 | 0.015972041 |
| ENSG00000161642 | ZNF385A | -0.473793931 | 0.015972041 |
| ENSG00000020181 | ADGRA2 | 1.462585829 | 0.015972041 |
| ENSG00000153993 | SEMA3D | 2.385145542 | 0.015972041 |
| ENSG00000159167 | STC1 | -0.742604055 | 0.01603399 |
| ENSG00000128039 | SRD5A3 | 0.777479384 | 0.016155137 |
| ENSG00000082781 | ITGB5 | 0.741839788 | 0.016451458 |
| ENSG00000102699 | PARP4 | 0.576630835 | 0.016743787 |
| ENSG00000188959 | C9orf152 | -1.372582578 | 0.017195686 |
| ENSG00000168679 | SLC16A4 | -1.372917814 | 0.017261245 |
| ENSG00000151718 | WWC2 | 0.738561796 | 0.017261245 |
| ENSG00000136238 | RAC1 | 0.575233958 | 0.017618152 |
| ENSG00000091409 | ITGA6 | 1.16162003 | 0.017817829 |
| ENSG00000185924 | RTN4RL1 | -0.553046234 | 0.018107152 |
| ENSG00000257671 | KRT7-AS | 1.165481696 | 0.01907405 |
| ENSG00000134504 | KCTD1 | 0.624744539 | 0.019107605 |
| ENSG00000127955 | GNAI1 | -1.027798495 | 0.019242308 |
| ENSG00000112183 | RBM24 | -0.990531229 | 0.019242308 |
| ENSG00000095321 | CRAT | -0.865522502 | 0.019242308 |
| ENSG00000128274 | A4GALT | -0.829521154 | 0.019828877 |
| ENSG00000196636 | SDHAF3 | -1.397373821 | 0.020109986 |
| ENSG00000172794 | RAB37 | -0.977071402 | 0.020109986 |
| ENSG00000214078 | CPNE1 | -0.656533189 | 0.020109986 |
| ENSG00000166833 | NAV2 | 1.535642143 | 0.020119476 |
| ENSG00000253276 | CCDC71L | 0.563147043 | 0.020319347 |
| ENSG00000131746 | TNS4 | -1.171019047 | 0.020401967 |
| ENSG00000198246 | SLC29A3 | 0.598720205 | 0.020600083 |
| ENSG00000172551 | MUCL1 | -0.924446266 | 0.022069823 |
| ENSG00000139289 | PHLDA1 | 0.748078573 | 0.022069823 |
| ENSG00000162496 | DHRS3 | 1.124026074 | 0.022069823 |
| ENSG00000115290 | GRB14 | -0.591415986 | 0.022820118 |
| ENSG00000230882 | - | 1.058086599 | 0.022820118 |
| ENSG00000249700 | SRD5A3-AS1 | 0.999653999 | 0.023737661 |
| ENSG00000164932 | CTHRC1 | -1.184322986 | 0.025845308 |
| ENSG00000106617 | PRKAG2 | 0.63438887 | 0.026066954 |
| ENSG00000117525 | F3 | 1.150824104 | 0.026129253 |
| ENSG00000106638 | TBL2 | 0.540525634 | 0.026634815 |
| ENSG00000127948 | POR | 0.628814201 | 0.028129504 |
| ENSG00000136383 | ALPK3 | 0.946763384 | 0.028129504 |
| ENSG00000166974 | MAPRE2 | 1.020963791 | 0.028129504 |
| ENSG00000196428 | TSC22D2 | 0.570377694 | 0.028296915 |
| ENSG00000232533 | - | 0.871876528 | 0.028296915 |
| ENSG00000144366 | GULP1 | 0.872570518 | 0.02856424 |
| ENSG00000149573 | MPZL2 | 0.616046997 | 0.02874668 |
| ENSG00000079805 | DNM2 | 0.50576267 | 0.029296616 |
| ENSG00000058091 | CDK14 | 0.863235517 | 0.029529631 |
| ENSG00000111058 | ACSS3 | 0.877215967 | 0.029529631 |
| ENSG00000153904 | DDAH1 | -0.750222768 | 0.029722896 |
| ENSG00000105877 | DNAH11 | 1.069564245 | 0.03040863 |
| ENSG00000139352 | ASCL1 | -1.227245488 | 0.030814065 |
| ENSG00000186187 | ZNRF1 | 0.518411577 | 0.030894038 |
| ENSG00000071859 | FAM50A | -0.607014788 | 0.031228581 |
| ENSG00000079257 | LXN | 0.619899203 | 0.031929235 |
| ENSG00000123095 | BHLHE41 | 0.828581841 | 0.033039427 |
| ENSG00000145824 | CXCL14 | -4.897880799 | 0.033334657 |
| ENSG00000117868 | ESYT2 | 0.415393014 | 0.033334657 |
| ENSG00000232931 | LINC00342 | 1.023885812 | 0.033334657 |
| ENSG00000143878 | RHOB | -0.526929328 | 0.03444086 |
| ENSG00000283538 | - | 1.063049627 | 0.034882785 |
| ENSG00000258667 | HIF1A-AS3 | 0.937183134 | 0.034945109 |
| ENSG00000230221 | - | 3.954877082 | 0.034999533 |
| ENSG00000154274 | C4orf19 | 0.568342414 | 0.037211785 |
| ENSG00000164266 | SPINK1 | 2.19454636 | 0.037527932 |
| ENSG00000185033 | SEMA4B | 0.573268786 | 0.037692183 |
| ENSG00000136830 | NIBAN2 | 0.610188212 | 0.037692183 |
| ENSG00000085831 | TTC39A | -0.494439943 | 0.039322054 |
| ENSG00000241258 | CRCP | 0.406772042 | 0.039396988 |
| ENSG00000163364 | LINC01116 | 0.801399256 | 0.039396988 |
| ENSG00000178821 | TMEM52 | 0.803856407 | 0.039396988 |
| ENSG00000127129 | EDN2 | 0.879138707 | 0.039396988 |
| ENSG00000189212 | DPY19L2P1 | 1.729940482 | 0.039396988 |
| ENSG00000085741 | WNT11 | 1.758298528 | 0.039396988 |
| ENSG00000168269 | FOXI1 | -1.251919043 | 0.03986553 |
| ENSG00000121671 | CRY2 | -0.552356356 | 0.03986553 |
| ENSG00000149929 | HIRIP3 | -0.526245111 | 0.03986553 |
| ENSG00000105711 | SCN1B | 0.641812557 | 0.03986553 |
| ENSG00000177283 | FZD8 | 0.812610501 | 0.040268156 |
| ENSG00000035664 | DAPK2 | 0.407928743 | 0.041586398 |
| ENSG00000132535 | DLG4 | -1.299819499 | 0.041646165 |
| ENSG00000181588 | MEX3D | 0.670043208 | 0.041865833 |
| ENSG00000146151 | HMGCLL1 | -1.144578957 | 0.042257282 |
| ENSG00000100842 | EFS | -1.633628557 | 0.043628975 |
| ENSG00000179083 | FAM133A | -1.014674402 | 0.043628975 |
| ENSG00000142871 | CCN1 | 0.561051523 | 0.043628975 |
| ENSG00000146757 | ZNF92 | 0.752250317 | 0.043628975 |
| ENSG00000087053 | MTMR2 | 0.901623924 | 0.043831175 |
| ENSG00000151655 | ITIH2 | -1.536930979 | 0.044104959 |
| ENSG00000156463 | SH3RF2 | 0.767569119 | 0.044104959 |
| ENSG00000008311 | AASS | 2.265648385 | 0.044348026 |
| ENSG00000123836 | PFKFB2 | 0.608542308 | 0.044884674 |
| ENSG00000260401 | - | 0.768734559 | 0.045235838 |
| ENSG00000181085 | MAPK15 | -0.725214109 | 0.04634039 |
| ENSG00000086619 | ERO1B | 0.835003061 | 0.046831615 |
| ENSG00000136828 | RALGPS1 | 0.58962825 | 0.046972194 |
| ENSG00000006576 | PHTF2 | 0.584285468 | 0.047111228 |
| ENSG00000203971 | - | 4.945148389 | 0.047615473 |
| ENSG00000071575 | TRIB2 | 0.812972534 | 0.047985894 |
| ENSG00000185973 | TMLHE | -0.534667382 | 0.048110254 |
| ENSG00000157502 | PWWP3B | -1.519630917 | 0.048310049 |
| ENSG00000259527 | LINC00052 | -2.729850582 | 0.048942107 |
| ENSG00000182985 | CADM1 | 0.705305357 | 0.048942107 |
| ENSG00000149591 | TAGLN | 0.832459222 | 0.048942107 |
| ENSG00000115602 | IL1RL1 | 1.750644752 | 0.049525138 |
| ENSG00000186197 | EDARADD | 1.300889719 | 0.049677344 |

**Supplementary table 3.** Sequences of sgRNA used for CRISPR/Cas9 as well as primers used for real time PCR

| **Gene** | **Primer Pair** |
| --- | --- |
| sgRNA1(Cas9wt) | F: 5’-**CACCg**GGACGAGATGAAGGCGTCTG-3’  R: 5’-**AAAC**CAGACGCCTTCATCTCGTCCc-3’ |
| sgRNA2.1 (Cas9n) | F: 5’-**CACCg**TCATCTCGTCCTCTGACTTC-3’  R: 5’-**AAAC**GAAGTCAGAGGACGAGATGAc-3’ |
| sgRNA2.2 (Cas9n) | F: 5’-**CACCg**TGCCACCGTGCTCACCGCCC-3’  R: 5’-**AAAC**GGGCGGTGAGCACGGTGGCAc-3’ |
| genomic MB | F: 5'-TGGGAAGACAGGGAGCTAAA-3'  R: 5'-GCTCTGCCATTATCCACCTC-3' |
| TWIST | F: 5'-GGAGTCCGCAGTCTTACGAG-3'  R: 5'-TCTGGAGGACCTGGTAGAGG-3' |
| SNAIL | F: 5'-GACCACTATGCCGCGCTCTT-3'  R: 5'-TCGCTGTAGTTAGGCTTCCGATT-3' |
| SLUG | F: 5'-TTCGGACCCACACATTACCT-3'  R: 5'-GCAGTGAGGGCAAGAAAAAG-3' |
| FN1 | F: 5'-TCGCCATCAGTAGAAGGTAGCA-3'  R: 5'-TACTTTCTTGATTTTCTTCCACAGCATA-3' |
| ZEB1 | F: 5'-TACAGAACCCAACTTGAACGTCACA-3'  R: 5'-GATTACACCCAGACTGCGTCACA-3' |
| MMP3 | F: 5'-CAACAAGAGCTAAGTAAAGCCAGTGG-3'  R: 5'-CTAGATATTTCTGAACAAGGTTCATGCT-3' |
| PI3K CA | F: 5'-AGTAGGCAACCGTGAAGAAAAG-3'  R: 5'-GAGGTGAATTGAGGTCCCTAAGA-3' |
| PI3K CB | F: 5'-CTGCCTGCGACAGATGAGTG-3'  R: 5'-TCCGATTACCAAGTGCTCTTTC-3' |
| FASN | F: 5'-CATCCAGATAGGCCTCATAGAC-3'  R: 5'-CTCCATGAAGTAGGAGTGGAAG-3' |
| EGLN1 | F: 5'-CAAATGGAGATGGAAGATGTGTG-3'  R: 5'-AATGTCAGCAAACTGGGCTTT-3' |
| EPO | F: 5'-ATGTGGATAAAGCCGTCAGT-3'  R: 5'-AGTGATTGTTCGGAGTGGAG-3' |
| CITED2 | F: 5'-ACCATCACCCTGCCCACC-3'  R: 5'-CGTAGTGTATGTGCTCGCCCA-3' |
| SOD1 | F: 5'-TACAAAGACAGGAAACGCTGG-3'  R: 5'-CCTCAGACTACATCCAAGGGAA-3' |
| SOD2 | F: 5'-AGGTGACTCTAACTTCCCTGGC-3'  R: 5'-CCCACAAGCACAGAAATAAAGGAGA-3' |
| SOD3 | F: 5'-CGTTCCTGGGCTGGCTGGGT-3'  R: 5'-ATGGCTGGAGTCGGGCACCTTT-3' |
| GPX3 | F: 5'-AGGTATGCGTGATTGTGTGTGT-3'  R: 5'-GGAGAACTGGAGAGAAAGGGTTG-3' |
| GPX4 | F: 5'-CGCTGTGGAAGTGGATGAAGA-3'  R: 5'-CTTGTCGATGAGGAACTTGGTGAA-3' |
| CAT | F: 5'-GACATTACCAAATACTCCAAGGCAA-3'  R: 5'-AACCCGATTCTCCAGCAACA-3' |
| ACTB | F: 5'-CTGGAACGGTGAAGGTGACA-3'  R: 5'-AAGGGACTTCCTGTAACAACGA-3' |
